# ExploreTurns: A web tool for the exploration, analysis, and classification of beta turns and structured loops in proteins; application to beta-bulge and Schellman loops, Asx helix caps, beta hairpins and other hydrogen-bonded motifs

**DOI:** 10.1101/2024.01.01.573820

**Authors:** Nicholas E. Newell

## Abstract

The most common type of protein secondary structure after the alpha helix and beta sheet is the four-residue beta turn, which plays many key structural and functional roles. Existing tools for the study of beta turns operate in backbone dihedral-angle (Ramachandran) space, which presents challenges for the visualization, comparison and analysis of the wide range of turn conformations. In this work, a new turn-local coordinate system and structural alignment, together with a set of geometric descriptors for turn backbone shape, are incorporated into ExploreTurns, a web facility for the exploration, analysis, geometric tuning and retrieval of beta turns and their contexts which combines the advantages of Ramachandran- and Euclidean-space representations. Due to the prevalence of beta turns in proteins, this facility, supported by its interpreter for a new general nomenclature which classifies H-bonded loop motifs and beta hairpins, serves as an exploratory browser and analysis tool for most loop structure. The tool is applied to the detection of new H-bonded loops, including short and “double” Schellman loops, a large family of beta-bulge loops with a range of geometries and H-bond topologies, and other motifs. Other applications presented here include the mapping of sequence preferences in Asx helix N-caps and an investigation of the depth dependence of beta-turn geometry. ExploreTurns, available at www.betaturn.com, should prove useful in research, education, and applications such as protein design, in which an enhanced Euclidean-space picture of turn and motif structure and the ability to identify and tune structures suited to particular requirements may improve performance.

## 1 Introduction

Of the five reported types of tight turns^1^ in proteins, the four-residue beta turn, first described by Venkatachalam^2^ in 1968, is by far the most common; it represents the most common type of protein secondary structure after the alpha helix and beta sheet and constitutes close to two-thirds of all loop residues^3^. Beta turns, which can occur either individually or in overlapping multiples^4^, exhibit a wide variety of backbone (BB) conformations and side-chain (SC) interactions, and they play key roles in multiple contexts in proteins. Either alone or as components of short loops, turns link helices/strands together into supersecondary structures which contribute to a protein’s tertiary fold, and turns are also common in the extended loops of intrinsically disordered regions, where they may form phase-separation nucleation points^5^.

Beta turns are also features of beta sheets, for which they can form entrance or exit structures or all or part of the chain-reversing loops between the antiparallel strands of beta hairpins^6,7,8^, and turns also occur in beta bulges^9,10^, which add irregularities at the edges of sheets that hinder the aggregation associated with the plaque diseases^11^. Beta turns are also found in beta-bulge loops^12^ (BBLs), which occur when beta bulges are found at the loop ends of beta hairpins, but also exist as independent structures. The occurrence of beta turns in multiple contexts associated with beta sheets suggests that mutations within turns may be a driver of plaque formation, a possibility highlighted by the finding that cross-alpha crystals can be converted to cross-beta fibrils by flipping a beta turn’s type^13^.

Beta turns are associated with helices as well as sheets; they are found at both ends of helices, where they can “cap” unsatisfied BB polar groups with BB or SC H-bonds^14,15^. The six-residue Schellman loop^16,17^ is a common C-terminal capping structure that can incorporate or overlap a beta turn, and a seven-residue “wide” form^18^ has also been identified. Beta turns can also interrupt helices, forming helix kinks^19^ that introduce design flexibility and allow a greater variety of SC placements^20^.

Other structures in which beta turns occur include beta helices^21^ and other solenoidal folds, beta^22^ and TIM^23^ barrels, beta propellers^24^, and multiple local motifs such as nests^25^, Asx and ST turns/motifs^26,27^ and ST staples^28^. Beta turns are components of all reported short H-bonded loop motifs, including the alpha-beta^28^ and crown bridge^29^ motifs in addition to the Schellman, wide Schellman and beta-bulge loops.

Beta turns may play important roles in protein folding, including as nucleation sites, established by stable intra-turn interactions, which help initiate folding and enforce chain registration^30–34^, and it has been shown that interactions involving beta-turn SCs can make a substantial contribution to overall protein stability^35^, and that the stability of proteins can be increased via amino acid (AA) substitutions in turns^36^.

Since beta turns often change the direction of the BB as it reaches the protein surface, they are frequently exposed to the surrounding solvent, and polar, hydrophilic SCs are overrepresented in turns. The common occurrence of turns at the protein surface also positions them well to interact with external ligands, including other proteins, and turns also participate in ligand binding within the protein body. Key beta-turn binding partners include metal ions, ADP/ATP, nucleic acids and heme groups. Beta turns play important roles in determining the conformation and specificity of enzyme active sites and antibody-combining sites^37^.

Beta turns are found in peptides as well as proteins; they can provide chain-reversal in the beta hairpins of cyclic antimicrobial bioactive peptides^38^, and it has been postulated that in many cases they represent the bioactive conformation of linear peptides^39^. Chemically modified beta-turn mimics can serve as asymmetric catalysts^40^. Beta turns and their mimics are employed in drug development studies^41^; it has been demonstrated that beta-turn mimics can inhibit plaque formation^42^.

Since their identification, beta turns have been the subject of a large body of work that has evolved their definition and classification (see brief reviews in^3,43^). The beta-turn definition used here^3^ describes a four-residue BB segment in which the alpha carbons of the first and last residues are separated by no more than 7 Å and the two central turn residues lie outside helices or strands; in about half of these structures the fourth turn residue donates a BB H-bond to the first residue (4→1 H-bond). ExploreTurns employs three beta-turn classification systems which specify turn geometry at different levels of precision in the Ramachandran space of BB dihedral angles; in increasing order of precision, these systems are referred to here as the “classical” types^44^, BB clusters^3^ and Ramachandran types^3,45,46^ (see the ExploreTurns online help for definitions).

As far as the author is aware, the only currently available tools that specifically support the study of beta turns are the web facility Motivated Proteins^28^ and its associated stand-alone application Structure Motivator^47^, which present turns as one of multiple small H-bonded motifs, including SC structures. ExploreTurns has a different focus, taking a beta-turn centered approach to the exploration of loop structure in a redundancy-screened, PDB-derived database. The tool incorporates multiple new features, including a turn-local Euclidean-space coordinate system and structural alignment for all beta turns^48^ (see Sections 5.2), a set of geometric descriptors that characterize the bulk BB shapes of turns^48^ (Section 5.3), statistical tools for the detection of SC motifs, and other features. In addition, ExploreTurns is the first tool to encompass beta turns without the 4→1 H-bond, which comprise about half of the database and include classical types VIa2, VIb and VIII, as well as 13 of the 18 BB clusters. Also new are ExploreTurns’ integration of structure selection, profiling, and 3D viewing onto a single web page and its support for rapid comparative structure browsing.

ExploreTurns’ turn-local coordinate system provides a common framework for the visualization and comparison of the wide range of beta-turn BB geometries, and also supports analyses of the many recurrent contexts that incorporate turns, including local BB structures, ligand-binding and active sites and supersecondary structures. The tool’s incorporation of geometric turn descriptors enhances the Euclidean-space representation of turns, characterizing meaningful modes of structural variation not explicitly captured by Ramachandran-space classification systems and complementing those systems^48^. Descriptors enable structural discrimination within the classical types or BB clusters, yielding major improvements in the specificity of the measurements of sequence preferences in turns (which the tool computes), and enabling the Euclidean-space tuning of turn geometry for compatibility with key SC interactions and contexts.

Since beta turns constitute almost two-thirds of all residues in loops^3^, and ExploreTurns also encompasses the four-residue BB segments N- and C-terminal to turns, the tool gives access to most loop structure in proteins. In addition to beta turns, turns of all lengths can be explored, along with short loops shaped by multiple BB H-bonds into local structural motifs such as the Schellman, alpha-beta and beta-bulge loops. This work introduces a general description of all of these motifs as “compound turns” composed of overlapping “simple” H-bonded turns of all types (lengths), and defines a nomenclature which classifies the motifs by their content of simple turns. ExploreTurns includes an interpreter for this notation which is applied here to explore the structures of H-bonded loop motifs and their variants and detect new motifs, including short and “double” forms of the Schellman loop, a new seven-residue variant of the classic Schellman loop, four- and six-residue analogues of the common five-residue alpha-beta loop^28^, seven-and eight-residue “beta brackets” formed of overlapping beta turns constrained by longer turns, and a large family of motifs which satisfy a new generalized definition of the beta-bulge loop and exhibit a range of geometries and H-bond topologies. The compound-turn nomenclature is also applied to the chain-reversing loops of beta hairpins, where the notation provides a hairpin classification that is more complete than the X:Y hairpin nomenclature^8^, since it describes the H-bond configuration within a hairpin’s loop as well as the configuration at the loop end of its beta ladder.

ExploreTurns is also applied to profile the conformations of helix-terminal Schellman loop/beta-turn combinations, map sequence preference vs. cap geometry in Asx helix N-caps, and investigate the relationship between beta-turn geometry and depth beneath the solvent-accessible surface.

## 2 Tool description

### 2.1 Feature summary

ExploreTurns is a database selection screen, structure profiler and graphical browser, integrated onto a single web page, which supports the analysis and retrieval of structures from a redundancy-screened, PDB^49^-derived dataset of 102,192 beta turns and their four-residue BB neighborhoods (their “tails”), using a wide range of selection criteria. Criteria include turn BB geometry classified in the Ramachandran space of BB dihedral angles (the classical types, Ramachandran types and BB clusters), ranges of the geometric turn descriptors (“span”, N- and C-terminal “half-spans”, “bulge”, “aspect”, “skew”, and “warp“; see Section 5.3), inclusionary/exclusionary patterns of BB H-bonds, BB dihedral angles, the *cis/trans* status of the turn’s peptide bonds, sequence motif content, the electrostatic energy of the turn’s potential 4→1 H-bond, DSSP^50^ secondary structure codes, the approximate orientations of the tails with respect to the turn (the “context vectors”), and the depth of the turn beneath the protein’s solvent-accessible surface. Required patterns of BB H-bonds in structures can be specified either in the position frame of the beta turn at the center of the 12-residue ExploreTurns structural window, in the frame of an H-bonded loop motif via the compound-turn notation, or by descriptive shorthand names for loop motifs. A summary of all ExploreTurns selection criteria, with examples of their use and links to the relevant help pages, is accessible from the tool page via the “Feature Summary/Examples” button.

ExploreTurns supports the comprehensive exploration and analysis of structure and interaction in beta turns and their BB contexts and the identification, tuning and retrieval of structures tailored to suit particular requirements. Browsing with the tool reveals the characteristic geometries of the classical turn types and BB clusters and the modes of variation characterized by the geometric turn descriptors. The tool supports the analysis and retrieval of sets of examples of any BB motif that can be defined by the tool’s selection criteria operating on its twelve-residue window centered on a beta burn, including beta hairpins from the four classes^6,7,8^, supersecondary structures, some beta bulges^9,10^ and nests^25^, and all H-bonded loop motifs that occur within four residues of a beta turn, including all previously reported loops along with newly-identified types.

The tool also gives access to sequence motifs, which are defined here as the specification of one or more AAs at particular positions in the turn/tails, whether or not they are associated with recurrent SC structures (which are termed “SC motifs”). The most common SC motifs in turns include Asx/ST turns/motifs^26,27^ and Asx/ST helix caps^14,15,26,27^, but turns and their BB contexts host many recurrent SC structures that can be identified with the aid of ExploreTurns’ motif detection tools, which rank sequence motifs by fractional overrepresentation or statistical significance within the selected set of structures, and include options that focus the search on particular residue positions or position ranges. In addition to H-bonded motifs, SC motifs in turns/tails include salt bridges, hydrophobic interactions, a wide range of aromatic-Pro and pi-stacking interactions, and cation-pi and pi-(peptide bond) relationships.

ExploreTurns supports the further investigation of SC motifs by providing direct access to the motif maps generated by the MapTurns tool^51^ (also available at www.betaturn.com), which enable the comprehensive exploration of the BB and SC structure, H-bonding and contexts associated with single-AA, pair, and many triplet sequence motifs in turns and their BB neighborhoods.

ExploreTurns provides three types of distribution plots involving the geometric turn descriptors which cover each classical turn type or BB cluster and support structure selection and analysis. Interpolated histograms of the distributions of the descriptors within types and clusters support structure selection by Euclidean-space turn BB geometry, and browsing with the guidance of these plots reveals the modes of geometric variation characterized by the descriptors (see Supplementary Section 1). Also available, for single-AA and pair sequence motifs with sufficient abundance within the turn and the first two residues of each tail, are plots of sequence motif overrepresentation vs. descriptor value. These plots, which yield major improvements in the precision of measurements of sequence preferences in turns, identify the descriptor regimes most compatible with each sequence motif, and therefore most likely to be compatible with any associated SC motif. Motif overrepresentation plots support the “tuning” of turn BB geometry for compatibility with a particular SC motif, via the selection of turns with descriptor values that maximize the fractional overrepresentation of the associated sequence motif (see Supplementary Section 2). The third type of distribution plot available in ExploreTurns heat-maps the values of the turn descriptors across the BB dihedral-angle (Ramachandran) spaces of the two central residues of the turns in each type, BB cluster, or the global turn set, revealing the bond rotations associated with descriptor variation as well as the conformations that yield the lowest energy for the 4→1 beta-turn H-bond (see Supplementary Section 3).

ExploreTurns supports the tuning of a turn’s context as well as its BB geometry: the “context vectors” may be used to select supersecondary structures in which turns link helices/strands into particular approximate relative orientations (see Supplementary Section 4), or identify the tail vs. turn orientations that optimize interactions, such as helix-capping H-bonds, that occur between turns and adjacent structures (see Section 3.3.1 and Supplementary Section 7).

### 2.2 The ExploreTurns screen

The top half of the ExploreTurns screen (Figure 1) is divided vertically into three sections. The buttons at the top of the screen provide access to extensive online help, including an introduction, a feature summary with examples, an in-depth user guide, and definitions of the compound-turn nomenclature, BBL shorthand notation, turn types, BB clusters and turn descriptors. The *Explore Loop Motifs* button opens a tour of H-bonded loop motifs and beta hairpins which includes schematics of structure and H-bonding as well as links which load sets of motif structures into the tool for browsing.

**Figure 1.**
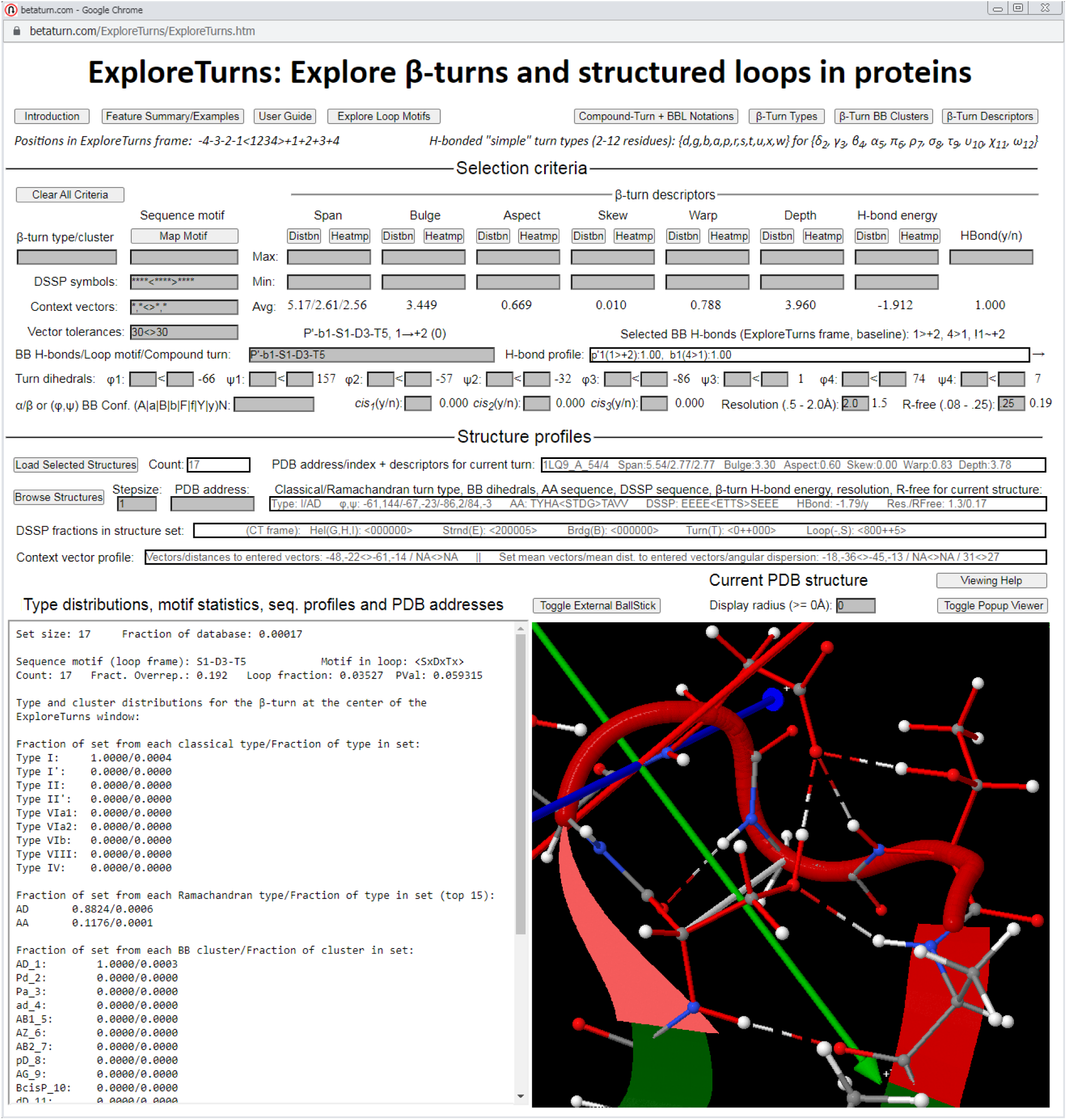
The ExploreTurns screen. The compound-turn label *P’-b1-S1-D3-T5* (see sections 5.4 and 5.5, or click *Compound-Turn + BBL Notations*) has been entered into the *BB H-bonds/Loop motif/Compound-turn* box to select the sequence motif that specifies Ser at position 1, Asp at position 3 and Thr at position 5 in the 6C2 beta-bulge loop (BBL), one of the new BBL motifs described here (Section 3.1.6). The structure is displayed in the turn-local coordinate system, with a white bar drawn between *C*_*⍺*1_ and *C*_*⍺*4_ in the β-turn to show the turn’s span. The loop’s cartoon BB and the sequence motif’s SCs are highlighted in red, with a lighter-red flash at the N terminus to aid orientation. The radius for the display of structures outside the 12-residue ExploreTurns window (which is always centered on a beta turn) has been set to zero in the *Display radius* box to remove clutter.

The *Selection criteria* section in the middle of the top half of the tool page is a database selection screen which accepts input in the shaded boxes. Average values of numerical criteria for the selected structures are displayed beneath or to the right of the input boxes when a set of structures is loaded. Also displayed, in the *H-bond profile* box, are the ranked occurrence frequencies of all BB H-bonds that occur in the turns/tails in the selected set, with the H-bonds labelled by the positions of their donor and acceptor residues expressed in the frame of the beta turn at the center of the ExploreTurns window in the format “M>N”, using the position key: -4-3-2-1<1234>+1+2+3+4 (where the brackets delineate the turn). When an H-bonded loop motif is selected using a compound-turn label (see sections 3.1.4, 5.4 and 5.5), the H-bond profile window displays only the H-bonds that occur within the motif, and the H-bonds are also labelled with their associated simple turn types, which can be copied into the compound-turn label to investigate the structures of the motif’s BB H-bond variants.

The *Structure profiles* section at the bottom of the top half of the tool page profiles the structure from the selected set that is currently displayed in the viewer, reporting values for all selection criteria, and also displays information on the selected set of structures as a whole, including profiles of DSSP secondary structure and context vectors.

At bottom left on the page, the profile window provides additional information on the selected set. The distributions of the turns at the center of the 12-residue ExploreTurns window across the classical types, Ramachandran types and BB clusters are displayed, along with ranked statistics for all single-AA sequence motifs that occur in the window (if no specific motif is entered in the *Sequence motif* box as a selection criterion), statistics for the selected single or multiple-AA motif (if a sequence motif is entered), or ranked statistics for sequence motifs that have been specified by “wild-card” motif entries, which support SC motif detection at particular positions or position ranges in the window (see *Sequence motif selection…/Examples/Ex. 3* in the feature summary). If an H-bonded loop motif is selected with a compound-turn label, ranked statistics for the single-AA and pair sequence motifs that occur in the loop are listed at the top of the profile window, and sequence motif labels can be copied into the compound-turn label to investigate the SC interactions that occur in the loop and their effects on the loop’s geometry. A log-odds sequence profile of the selected set of structures is also provided in the profile window, and a list of PDB addresses for the structures is displayed at the bottom of the window, along with a list of PDBIDs alone, which can be copied to www.rcsb.com/downloads to download the structures from the PDB.

At bottom right on the page, a JSmol viewer displays the structures from the selected set in the turn-local coordinate system (see Section 5.2 and Figure 9), in the contexts of their PDB files (which may be downloaded individually in the viewer). By default, structure in the PDB files external to the ExploreTurns window is displayed out to a radius of 30 Å from the turn, in translucent cartoon only, but the user can change the display radius and add a ball/stick representation.

## 3 Applications

### 3.1 Classifying and exploring H-bonded loop motifs

#### 3.1.1 Roles of H-bonded loop motifs

Helices and beta sheets, stabilized by repetitive patterns of BB H-bonds, provide the extended “scaffolds” of protein structure, but there is also a role in protein architecture for local H-bonded motifs which complement the repetitive structures by shaping the BB into irregular local conformations which can serve as “accessories” to the scaffold. Motifs such as the Schellman, alpha-beta and crown bridge loops are often found at the ends of helices, where they can stabilize the helix and shape its entrance and exit loops, contributing to the geometries of supersecondary structures and, by extension, to a protein’s tertiary fold. Local motifs are also found in association with beta sheets: beta-bulge loops commonly occur at the loop ends of beta hairpins, where they shape the BB for structural roles or support function in contexts such as ligand-binding or active sites (see Figure 8), and motifs such as the alpha-beta, beta-bulge and Schellman loops can form beta-sheet entrance or exit structures. In addition to their associations with helices and sheets, H-bonded loops play independent structural and functional roles; they are versatile motifs which combine the constraints imposed by their BB H-bonds with a degree of conformational flexibility, filling a niche between the simple turns characterized by a single H-bond and the highly constrained repetitive structures.

#### 3.1.2 H-bonded loop motifs as “compound turns”

The BB geometry of a simple H-bonded turn is restricted by the constraint imposed by its H-bond to a set of conformations characteristic of its type (length), and loops formed of H-bonded turns with sufficient overlap are constrained, to some degree, along their full lengths by overlapping H-bonds. All reported H-bonded loop motifs in proteins are formed of overlapping H-bonded turns, and although the component turns in these motifs may be shaped by the H-bonds associated with their overlapping partners, each turn remains an example of its type, since its own constraining H-bond is still present. Accordingly, H-bonded loop motifs can be described as “compound turns” and classified by the types of their component simple turns and the positions of those turns within the motifs. Before formally defining compound turns and introducing a nomenclature which classifies them and supports their exploration in ExploreTurns, it is helpful to review what is known about the simple turns that compose them.

#### 3.1.3 Review of “simple” H-bonded turns

H-bonded “tight” turns are classified into types {δ, γ, β, α, π}, with lengths of two to six residues^1^. The four longest types are divided into sub-types in which the H-bond that links the turn’s terminal residues is donated in either the C→N (“normal”) or N→C (“reverse”) direction (delta turns are found only in the reverse form), and the normal and reverse sub-types are further partitioned by geometry, based on the BB dihedral angles of the turn’s central residues.

Figure 2 presents example structures and schematics for the five types of tight turns. The two-residue, H-bonded delta turn (Figure 2a), found only in reverse form, is extremely rare: just six nonredundant prototype structures were identified in proteins in a 2015 study^52^. Four of these examples exhibit “p*cis*P” Ramachandran-space conformations, with ‘p’ (+φ polyproline) and ‘P’ (-φ polyproline) conformations at their first and second residues respectively, and their single peptide bonds in *cis* conformation (click *β-Turn Types* on the ExploreTurns page for a map of Ramachandran space). The p*cis*P conformation, shown in Figure 2, is also the most common delta turn type in the ExploreTurns database, with four examples out of the 15 turn structures. However, while previous theoretical and observational studies have identified *cis* as the exclusive conformation for the delta-turn peptide bond^52^, six of the delta turns in the ExploreTurns database exhibit *trans* peptide bonds, and three of these (4PDX_A_416, 4QHW_A_253 and 5HDM_A_219) show no outliers in the PDB’s validation plots^53^.

**Figure 2.**
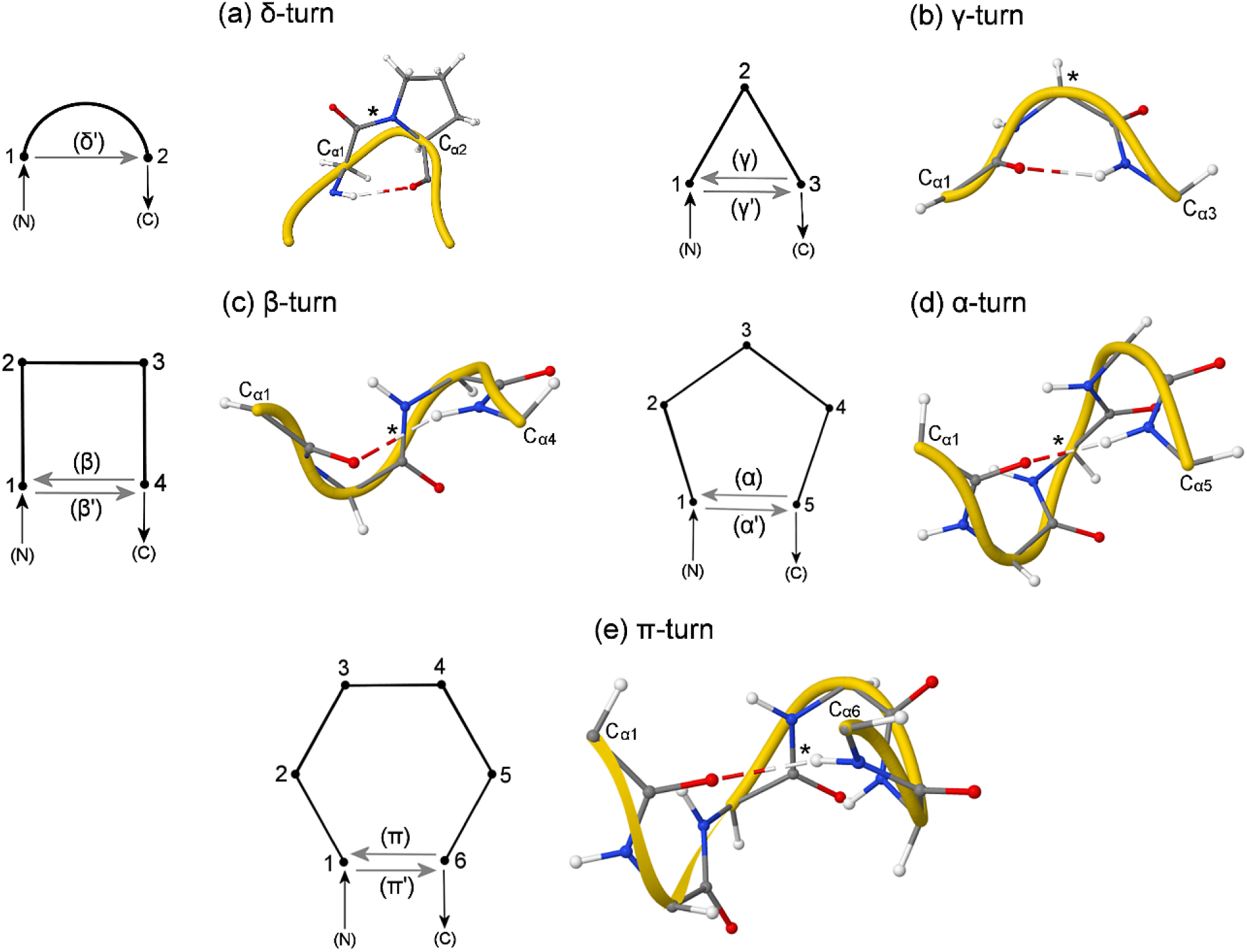
Schematics and example structures for the five types of “simple” H-bonded “tight” turns. Structures are shown for H-bonded turns of types {δ, γ, β, α, π}, with lengths of 2-6 residues. Schematics show H-bonds as arrows oriented in the direction of H-bond donation: arrows pointing left represent the “normal“^1^ turns of each type, with donation in the C→N direction, while those pointing right represent “reverse” (primed) turns, with donation in the N→C direction. With the exception of the δ-turn, which exists only in reverse form, all types are shown in their normal forms. Each turn is labelled at the α-carbons of its terminal residues, and its “turn center”, which lies either in the center of its middle peptide bond (when the turn contains an even number of residues), or at the α-carbon of its central residue (when the turn has an odd length), is marked with an asterisk. A “turn plane“^48^ (not shown) is defined which passes through the three labelled points in each turn, and the relatively flat δ- and γ-turns are viewed from above this plane, while the longer turns are viewed approximately “edge-on” to the plane to best illustrate and compare their conformations. **(a)** The two-residue reverse δ-turn, in its most common p*cis*P conformation (2FYF_A_25). **(b)** The three-residue γ-turn, in its “classic” geometry, with ‘g’ conformation at its central residue (4D6Z_A_263). **(c)** The four-residue β-turn, in its most common geometry (classical type I), with right-handed helical or helix-adjacent conformations at both central residues (3LY1_A_228). **(d)** The five-residue α-turn, in its most common geometry, with right-handed helical conformation at all three central residues (3SUK_A_46). **(e)** The six-residue π-turn, in its most common geometry, with right-handed helical conformation at the first three central residues and left-handed helical conformation at the last central residue, as seen in the Schellman loop (1W07_A_361).

The three-residue, H-bonded gamma turn (Figure 2b) is found in either a “classic” conformation (shown), with its central residue falling in the ‘g’ region of the Ramachandran plot beneath the left-handed helical region, or the “inverse” conformation, in which that residue lies in the ‘G’ region, above the right-handed helical region^54,55^ (’g’ and ‘G’ are related by inversion symmetry).

The four-residue, H-bonded beta turn (Figure 2c) is found in abundance in five of the eight structurally classified classical beta-turn types^44^, including types {I, I’, II, II’, VIa1}. Classical type I (shown in the figure), with its pair of central residues occurring within or adjacent to right-handed helical conformation, contains about 60% of all examples.

Five-residue, H-bonded alpha turns (Figure 2d) have been grouped into three structurally classified types and an unclassified category. Alpha turns of the dominant structurally classified type (shown in the figure), which represents about three-quarters of all examples, exhibit right-handed helical conformation at all three central residues^56^.

Six-residue, H-bonded pi turns (Figure 2e) have been grouped into eleven overall classes based on the conformations of their four central residues^57^. The dominant pi-turn conformation exhibits right-handed helical conformation at the first three central residues and left-handed helical conformation at the fourth^58^, as seen in the Schellman loop.

#### 3.1.4 Compound-turn definition and nomenclature

The compound turn (CT) is defined here as a short loop incorporating two or more H-bonded turns, in which both terminal residues are linked by a BB H-bond to at least one other loop residue, and all covalent bonds in the main-chain path between the loop’s first and last H-bonded atoms lie within the H-bond span of at least one turn. This definition guarantees a degree of constraint across the entire loop by preventing free rotation around any internal covalent bond, while allowing “open” loops (e. g. the “beta bracket” in Figure 3), in which the terminal residues are not linked by a single shared BB H-bond. In the more common, “closed” CTs, one or more shorter H-bonded turns lie embedded within the longest, “framing” H-bonded turn; they may share one terminal residue with that turn.

**Figure 3.**
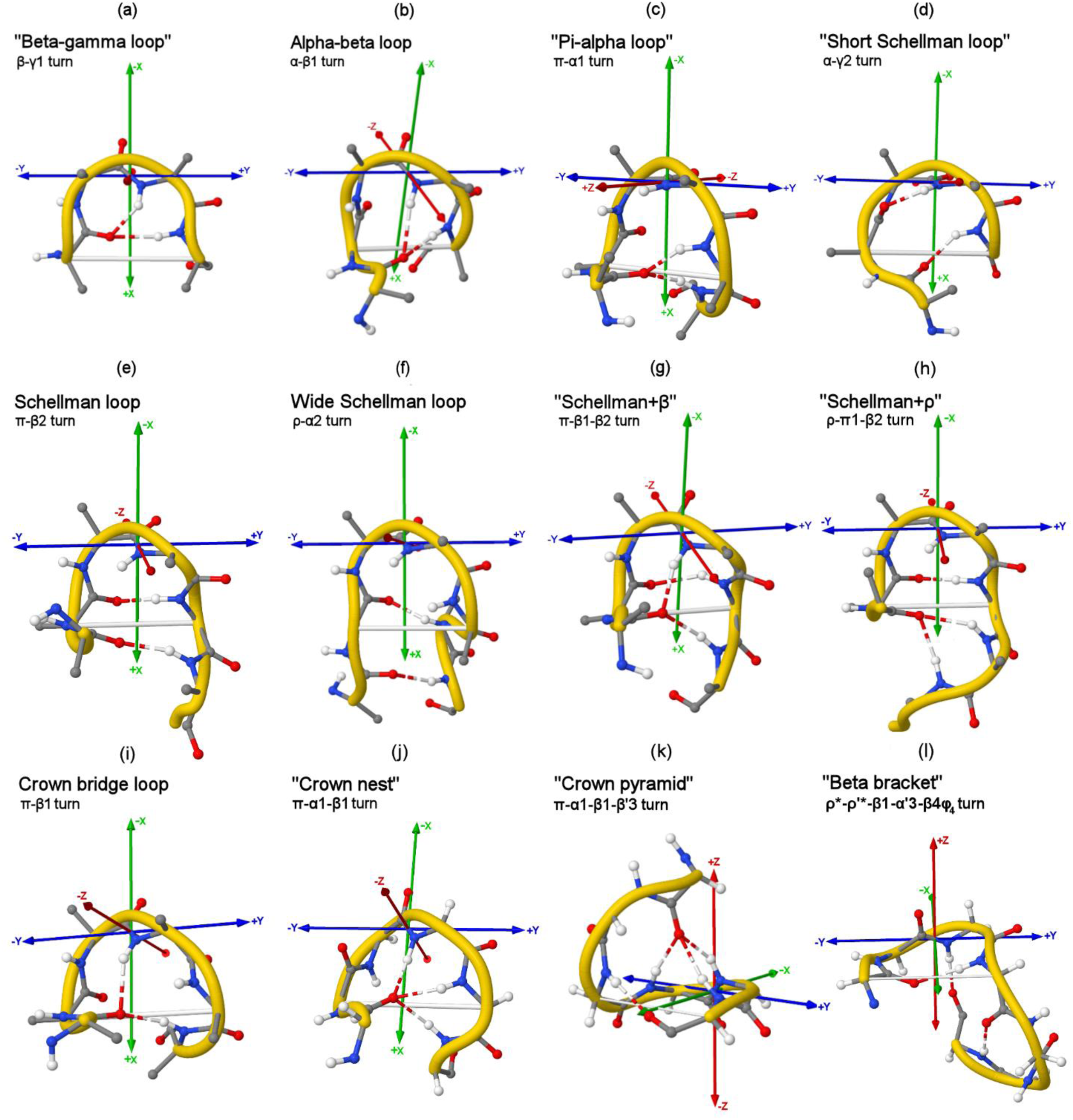
Classifying and comparing H-bonded loop motifs (excluding beta-bulge loops and additional motif variants). Loops are labeled with motif names as well as compound-turn labels that classify the structures by the lengths (types) and start positions of their component H-bonded turns. Simple transformations relating the loops to each other are suggested. **(a)** The four-residue beta-gamma loop shows a high frequency of outliers in PDB validation checks^53^, but some examples pass checks, including 3Q23_A_949, which binds Mn/G2P at the active site of bacteriophage N4 RNA polymerase. **(b)** The common alpha-beta loop^28^ adds a residue inside the beta-gamma loop’s γ1-turn, yielding a β1-turn. **(c)** The pi-alpha loop adds a residue inside the alpha-beta loop’s β1-turn, forming an α1-turn. **(d)** Bond rotations that transition the alpha-beta loop’s β1-turn to a γ2-turn yield the short Schellman loop. **(e)** The classic Schellman loop^16,17^ adds a residue inside the short Schellman’s γ2-turn, forming a β2-turn. **(f)** The wide Schellman loop^18^ adds a residue inside the Schellman’s β2-turn, forming an α2-turn. **(g)** A rotation of the Schellman loop’s fourth BB NH group adds a β1-turn, yielding the Schellman+β loop^28^. **(h)** A ρ-turn H-bond extends the Schellman loop, directing the BB towards -y. **(i)** Bond rotations disrupt the Schellman+β loop’s β2-turn and redirect its final residue, yielding the crown bridge loop^29^. **(j)** A rotation of the crown bridge loop’s fifth BB NH group adds an α1-turn, yielding the crown nest^29^. **(k)** Major conformational changes in the crown nest yield a β’3-turn, forming the crown pyramid^60^, which binds Mg and substrate at the active sites of Mg phosphatases in all three kingdoms. **(l)** A pair of overlapping β-turns is constrained into an approximate right-angle orientation by an α’-turn, forming the beta bracket. See also the “long beta bracket” listed in table S4, which can form a structurally conserved “loop the loop” sheet exit (CT label *u*-u’*-b3-p’5-b6-\E1*) in the alpha/beta hydrolase superfamily.

A CT’s conformation is constrained to a degree which depends on factors including the ratio of the number of BB H-bonds in the loop to the number of loop residues, the degree of overlap of BB H-bonds, and the presence of SC interactions, BB-constraining proline residues, or BB-freeing Gly residues in the loop. Recurrent CTs exhibit “baseline” forms, which contain only the BB H-bonds that define the motif, as well as BB H-bond variants, in which the structures are constrained by additional BB H-bonds, and loop structures may also be shaped by SC interactions into SC variants.

Including previously reported motifs and their BB H-bond variants and those identified here, more than forty H-bonded loops are presented in this work and available in the ExploreTurns loop browser, and the abundance and variety of loop types highlights the need for a nomenclature that classifies all CTs. In the concise notation introduced here (summarized in Section 5.4, or click *Compound-Turn + BBL Notations*), the Greek letters that specify the types (lengths) of a loop’s component H-bonded turns are concatenated with dashes, along with the start positions within the loop of each component turn (except in the case of the loop’s framing turn, which is listed first, without a position specifier). A component turn’s letter is primed if the H-bond that defines it is donated in the N→C (reverse) direction, rather than the C→N (normal) direction, and H-bonds oriented in either direction can be specifically excluded by appending ‘*’ to a component turn’s entry, after any prime character and before the component turn’s position specifier, which terminates the entry. The length of a loop is conveyed by the type of its framing turn, and if this turn is specified with ‘*’ exclusions that rule out both its normal and reverse forms, then it serves only to frame an open loop. To avoid confusion with the names of supersecondary structure motifs (e.g. βαβ), a CT label always concludes with the word “turn”.

Since the existing classification for simple H-bonded turns only encompasses structures with lengths up to six residues, additional turn types are needed to cover motifs such as the wide Schellman loop^18^ (seven residues), double Schellman loop (nine residues), and the longer beta-bulge loops identified below, which contain up to eight residues, with up to ten residues in their “extended” forms. In addition, beta hairpins, which can be classified by the lengths of their chain-reversing loops and the H-bond configurations at the loop ends of their beta-ladders using the X:Y nomenclature^8^, can also be described in the CT notation (see Section 3.2 and the loop motif browser), and the availability of longer component turn types allows hairpins with longer chain-reversing loops to be described by the notation and selectable in ExploreTurns. For these reasons, the Greek letters following π are included in the nomenclature, with the exception of φ and ψ, which are reserved for the specification of dihedral angles (see below). The complete set of turn specifiers is {δ, γ, β, α, π, ρ, σ, τ, υ, χ, ω}, which includes component turns with lengths of 2-12 residues, covering the full ExploreTurns window and enabling the classification of hairpins with loop lengths up to ten residues (the 10:10, or ω-ω’ hairpin).

For loops which show conformational variants for which dihedral information is a useful discriminator (e. g. the Schellman and short Schellman loops, the beta bracket and some BBLs), the CT notation adds dihedral-angle specifiers: ‘ϕ’ or ‘ψ’ is appended to the CT label for each constrained angle, with the constrained residue position(s) (in the loop’s frame) indicated either as superscript(s), if the angle is restricted to positive values, or subscripts for negative values. Loop BB geometry may also be classified in the CT label using the “right-handed” and “left-handed” α/β conformations {αR, αL, βR, βL}, which support (for example) the selection of beta-turn geometries within CTs (see Supplementary Section 11 for the definitions of the α/β conformations applied here).

ExploreTurns includes an interpreter for the CT notation which supports the exploration of H-bonded loop motifs and beta hairpins and the detection of new motifs. The CT interpreter extends the notation to include entries which specify sequence motifs or secondary structures within the loop, or enable the selection of loops that occur in particular position frames with respect to the beta turn at the center of the ExploreTurns window. CT labels are entered in ExploreTurns after translation using the Beta Code^59^ romanization of the Greek script. The CT interpreter is described in section 5.5 (or click *Compound-Turn + BBL Notations*), and Table 1 presents examples of CT labels which demonstrate its features.

**Table 1.** CT examples.

| CT label | Motif | Romanization |
| --- | --- | --- |
| $\gamma$ turn | $\gamma$ turn, most common position frame | g |
| α-β1 turn | <u>Alpha-beta loop:</u><br>All BB H-bond variants, most common frame<br>Heme binding with Cys and His<br>Heme binding, all position frames | a-b1<br>a-b1-C1-C4-H5<br>a-b1-C1-C4-H5_* |
| $\alpha'$-β1 turn | <u>5C1 (type 1) β-bulge loop (BBL):</u><br>All BB H-bond variants, most common frame<br>BB H-bond variant adds a1, a "normal" α-turn<br>Adds D3T5 sequence motif<br>"Extended" 5C1 BBL (\e's exclude β-ladder) | a'-b1<br>a'-a1-b1<br>a'-b1-D3-T5<br>t-a'3-b3-\e1-\e2-\e8-\e9 |
| $\pi'$-α1 turn | <u>6C1 (type 2) BBL:</u><br>2<sup>nd</sup> most common position frame aligns the loop's "upright chair back" with the β-turn at the center of the ExploreTurns window<br>Metal-ion binding in e. g. Zn fingers/ribbons | p'-a1_2<br>p'-a1-C1-C4 |
| $\pi'$-β1 turn | <u>6C2 BBL:</u><br>Baseline (all BB H-bond variants excluded)<br>Excludes only 6C1 BBLs, using a*1<br>"6C2-" variant (\f4), with T4 H-bond bridge<br>Metal-ion binding with an Asp triplet | P'-b1<br>p'-a*1-b1<br>P'-b1-\f4-T4<br>P'-b1-D1-D3-D5 |
| $\rho'$-β1-β4 turn<br>$\rho'$-β1-β4$\alpha^{2,3}\beta^5\alpha_6$ | 7NC3 BBL, baseline, all position frames<br>Selects type I ($\alpha R\alpha R$) and II ($\beta R\alpha L$) β-turns | R'-b1-b4_*<br>R'-b1-b4-\A2-\A3-\B5-\a6 |
| τ-$\tau'$ turn | <u>7:7 β-hairpin:</u><br>BB H-bonds excluded from loop with *1~9<br>Ser, Asp H-bonds shape loop into a "hook" | t-t'-*1~9-\E1-\E9<br>t-t'-S2-D4-*1~9-\E1-\E9 |
| π-β2 turn<br>π-β2$\varphi^{3,4}\varphi_5$ turn | <u>Schellman loop:</u><br>All BB H-bond variants, most common frame<br>Strict baseline, BB H-bond variants excluded<br>Permissive baseline, extern. H-bonds allowed<br>Restricted to "dual" helix caps, all frames<br>"Left-handed", all frames | p-b2<br>P-b2<br>@p-b2<br>p-b2-\H1-\H2_*<br>p-b2-\F3-\F4-\f5_* |
| π-α1-β1-β'3 turn | "Crown pyramid", 3<sup>rd</sup> position frame aligns "base" with the central ExploreTurns β-turn | p-a1-b1-b'3_3 |
| $\rho^*$-$\rho'^*$-β1-$\alpha'$3-β4$\phi_4$ turn | "Beta bracket" (open loop), 2<sup>nd</sup> β-turn orients towards -z in the frame of the first (\f4) | r*-r'*-b1-a'3-b4-\f4 |

**Notes:** When a CT is loaded without specifying a baseline mode, all H-bonded component turns that occur in the loop are listed in the *H-bond profile* box, and turns may be copied into the CT label to investigate the effects of their BB H-bonds on loop structure. The loop's sequence motifs, ranked in the profile window, may also be copied into the label.

#### 3.1.5 Classifying and comparing H-bonded loop motifs

Figure 3 compares the structures of 12 H-bonded loop motifs, including seven previously described loops and five new motifs detected with ExploreTurns using the CT interpreter (the figure does not include beta-bulge loops^12^, as these are treated separately below). Previously reported structures include the six-residue Schellman loop^16,17^, which commonly forms a C-terminal helix cap with its pair of BB H-bonds, along with two related structures: a common Schellman variant^28^, referred to here as “Schellman+β”, which adds a second internal H-bonded beta turn to the “classic” loop, and the wide Schellman loop^18^, which adds a residue inside the classic motif’s beta turn, yielding an alpha turn. Also included are the common five-residue alpha-beta loop^28^ as well as the crown bridge loop^29^ and two variants referred to here as the “crown nest”, which adds an H-bonded alpha turn to the loop, and the “crown pyramid“^60^, which adds a reverse H-bonded beta turn to the crown nest, yielding a motif which binds Mg and substrate at the active sites of Mg phosphatases in all three kingdoms.

Motifs detected with ExploreTurns and displayed in Figure 3 include the “pi-alpha” and “beta-gamma” loops, which add and drop residues from the alpha-beta loop respectively, the “short Schellman loop”, which drops a residue from the Schellman loop’s internal beta turn (forming a gamma turn), a seven-residue variant of the Schellman loop, dubbed “Schellman+ρ”, which extends the motif at its C-terminus with an H-bonded rho turn, and the “beta bracket”, an overlapping pair of beta turns constrained into an approximate right-angle orientation by a reverse alpha turn.

Table S4 in Supplementary Section 5 lists the romanized CT labels for all of the motifs shown in Figure 3, as well as labels for additional loops or variants identified with ExploreTurns, including the “Schellman310” motif, in which the classic loop incorporates a 3_10_ helix, and the “double Schellman loop”, in which two instances of the classic motif (usually right-handed followed by left-handed) combine to form a nine-residue open loop. Table S4 also lists the “long beta bracket”, an eight-residue form of the bracket that substitutes a pi turn for the loop’s alpha turn and can occur as a structurally conserved “loopback” sheet exit motif in the alpha/beta hydrolase superfamily (CT label *u*-u’*-b3-p’5-b6-\E1*). Finally, Table S4 lists the label for the “α+Schellman” motif, a previously identified structure in which the classic loop overlaps a preceding alpha turn^28^.

The loops in Figure 3 are labelled with their motif names and also classified in the CT nomenclature, which facilitates their comparison by explicitly indicating the differences in their component-turn profiles. Simple transformations are suggested in the figure caption which relate the structures to each other by addition or subtraction of loop residues or component turn H-bonds. Loops can be explored and profiled in the ExploreTurns motif browser, where schematics are displayed, or by entering a motif’s name or CT label into the *BB H-bonds/Loop motif/Compound turn* box and clicking *Load Selected Structures* (“loop” is optional in motif names, and Greek letters in a name are translated to English words for entry in the tool).

#### 3.1.6 Generalizing the beta-bulge loop

The “classical” definition of the type 1 beta-bulge loop (BBL1)^12^ describes a five residue motif which is commonly found at the loop ends of beta hairpins, where it can be formed by the insertion of a residue that splits the pair of reciprocal BB H-bonds at the loop end of the C-terminal strand of a 2:2 hairpin, forming a new chain-reversing loop containing an H-bonded beta turn and a “bulge” at the loop’s C-terminus constrained at its ends by the split H-bond pair (to view a 2:2 hairpin, enter *2:2* into the *BB H-bonds/Loop motif/Compound turn* box in ExploreTurns and click *Load Selected Structures*; to view a BBL1 in a hairpin, enter “BBL1” and click *Browse Structures* after loading). The classical BBL definition also describes a second motif, which can be formed by adding a residue inside the BBL1’s beta turn, forming an alpha turn (enter and load “BBL2”).

Although the name “BBL” refers to the beta bulges that occur in beta sheets/hairpins, the definitions of these loops do not depend on the presence of a beta ladder, since the motifs can also be found as independent structures. Accordingly, the most salient feature of BBLs is that they include a loop-terminal residue with a split pair of BB H-bonds, in which one member of the pair links the motif’s terminal residues, closing the loop and defining the structure’s extent, while the other member links to an interior loop residue, simultaneously forming an interior H-bonded turn and a bulge constrained at its ends by the split H-bond pair. In both the type 1 and 2 BBLs, the bulge contains only one added residue and occurs at the loop’s C-terminus, but neither of these conditions is structurally required. These observations motivated a search for structures that would satisfy a generalized BBL definition requiring only the presence of a split pair of BB H-bonds closing a loop and forming a turn and a bulge, without limiting the bulge’s length to one residue or requiring that it occur at the loop’s C-terminus.

A search for generalized BBLs using the CT interpreter and other features of ExploreTurns yielded a large family of motifs with lengths up to eight residues and bulges which add up to four residues and occur at either the N- or C-terminus of the loop. These BBLs are all less common than the classical pair and some are very rare, but like those motifs they occur either at the loop ends of beta hairpins or independently and can be found at ligand-binding and active sites (see Section 3.1.7). The longer BBL types exhibit greater BB freedom in their bulges than that seen in the classic pair, but recurrent approximate geometries can be identified, and the motifs show BB variants constrained by additional BB H-bonds and SC variants shaped by internal SC/BB and SC/SC interactions.

Since the CT nomenclature encompasses all short H-bonded loops, it applies to BBLs and classifies them by their H-bonded component turns. However, since the structures of BBLs have a characteristic general form, it is also possible to classify them in a simple, motif-specific notation: loop types are labelled in a three-component format which specifies the motif’s overall length, the location of the bulge at the N- or C-terminus, and the bulge length (the number of residues added to form the bulge). BBLs can be selected in ExploreTurns by entering either their CT labels or their BBL type names into the *BB H-bonds…* box (or they may be explored in the loop motif browser by clicking links).

Figure 4 displays example structures for four BBLs with C-terminal bulges, including types {5C1, 6C2, 7C3, 6C1}. The 5C1 BBL corresponds to type 1 in the classical nomenclature, while the 6C2 and 7C3 BBLs can be formed by successive insertions of additional residues into the 5C1 BBL’s bulge. As stated above, the 6C1 BBL (classical type 2) can be formed by the insertion of a residue into the 5C1 BBL’s beta turn. Each BBL in Figure 4 can be classified into conformational variants by the sign of the ϕ BB dihedral angle of the last residue in each loop’s beta turn (the second overlapping beta turn in the case of the 6C1 BBL), which correlates with whether the bulge orients above (+) or below (-) the beta-turn’s plane^48^ (see Section 5.2 for the plane’s definition). Table S5 in Supplementary Section 6 supports the exploration of the +/-variants of the BBLs displayed in Figure 4 by listing the compound-turn labels for both conformations of each loop; the variants can also be explored in the tool by entering a BBL’s type name with a “+/-” suffix. Table S5 also lists CT labels for “extended” forms of the 5C1, 6C2 and 6C1 BBLs, in which the motifs lie embedded within longer H-bonded “normal” turns (explore by adding “extended” to the BBL type names). Since the addition of a single “reverse” BB H-bond would result in the classification of extended BBLs as beta hairpins, these motifs may, in some cases at least, represent evolutionary intermediates between beta hairpins and independent BBLs.

**Figure 4.**
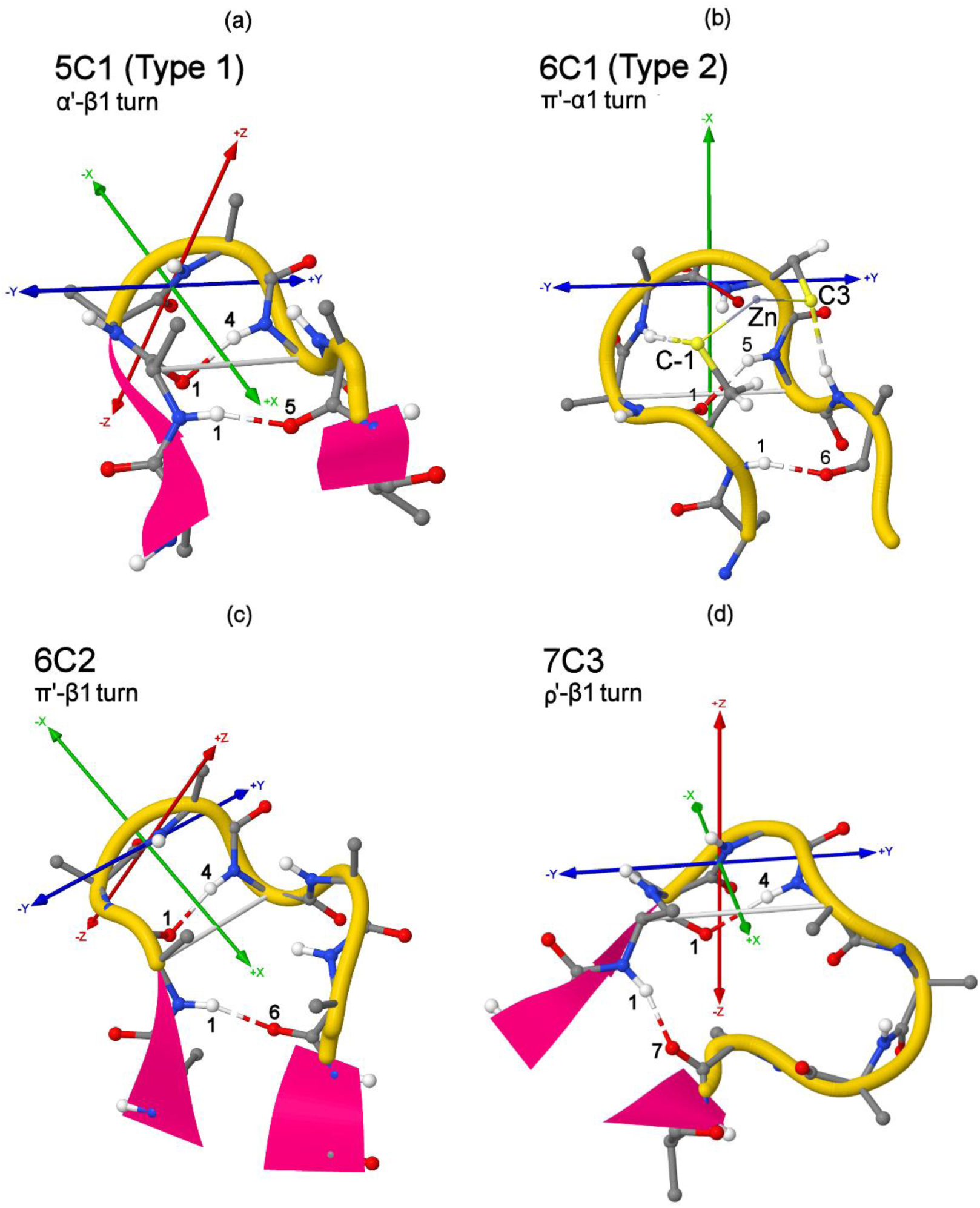
Examples of beta-bulge loops with C-terminal bulges. Structures are labelled using both a BBL-specific notation that specifies overall loop length, bulge location (N/C-terminal) and bulge length, and the compound-turn nomenclature for all H-bonded loop motifs, which classifies structures by the types and start positions of their component H-bonded turns (see Section 5.4). Each motif’s defining H-bonds are labelled at their ends with residue positions in the loop. To view examples of a BBL, enter its type name into the *BB H-bonds/Loop motif/Compound turn* box and click *Load Selected Structures*. **(a)** 5C1 (type 1); common 5C1+ conformational variant. In its 2:2 hairpin context (shown here), this motif can be formed by inserting a residue at the loop end of the C-terminal strand of a 2:2 beta hairpin, splitting the reciprocal pair of beta-ladder H-bonds. As with other BBLs, this motif also occurs outside hairpins. **(b)** 6C1 (type 2); common 6C1+ variant, in its “rubredoxin knuckle“^61^ metal-ion-binding form found in contexts such as Zn fingers/ribbons. This loop adds a residue inside 5C1’s β-turn, forming an α-turn. **(c)** 6C2; 6C2+ variant, adds a residue to 5C1’s bulge. **(d)** 7C3; 7C3-variant, adds a residue to 6C2’s bulge.

The shorthand names used to select the BBLs displayed in Figure 4 are translated in the tool to CT labels with capitalized framing turns that specify the motifs in strict baseline format, which rules out structures with additional BB H-bonds within the loop or between loop residues and other residues in the ExploreTurns window (see section 5.5). However, if a BBL (or any CT) is instead loaded using a CT label with a lowercase entry for its framing turn, then the selected set of structures includes any loop variants with additional BB H-bonds, and the corresponding additional H-bonded component turns are listed in the *H-bond profile* box along with the motif’s defining H-bonded turns. The effects of these additional H-bonds on loop structure can be explored by copying the component-turn labels from the *H-bond profile* box into the CT label in the *BB H-bonds…* box and loading the structures of the resulting BB H-bond variants (see *H-bonded loop motifs/Examples/Ex. 3* in the feature summary).

The 5C1 (type 1) BBL in Figure 4a demonstrates that a CT’s BB H-bond variants may exhibit conformations very different from the loop’s baseline form. This motif’s most common BB variant, which occurs in 8% of loop examples in the database, adds a normal alpha-turn H-bond, yielding a combined 5C1 BBL/alpha-beta loop, and 92% of the structures of this variant show the “5C1-” conformation, compared to only 2% of the structures of the baseline motif. To compare the 5C1 BBL’s baseline form to its most common variant, load the baseline loop with the CT label *A’-b1*, then load the variant with the label *A’-a1-b1*.

The 6C2 BBL in Figure 4c demonstrates that a CT’s conformational variants may exhibit distinct sets of SC interactions. The top ten most overrepresented single-AA sequence motifs in the 6C2+ and 6C2-variants share just one motif in common (proline at loop position 2), and the loop structures illustrate the distinct SC/BB and SC/SC interactions which likely stabilize each conformation. To compare the SC interactions in these variants, load “6C2+” and then “6C2-” in the tool; all single-AA sequence motifs present in each loop and all pair motifs with p-values of.10 or less are ranked in the profile window, and sequence labels can be copied into the CT label in the *BB H-bonds…* box to explore the associated SC structures and their effects on loop conformation (see *H-bonded loop motifs/Examples/Ex. 4* in the feature summary).

Figure 5 presents schematic diagrams of the structures and H-bond topologies of a larger set of BBLs with C-terminal bulges, including four additional types with bulge lengths of up to four residues.

**Figure 5.**
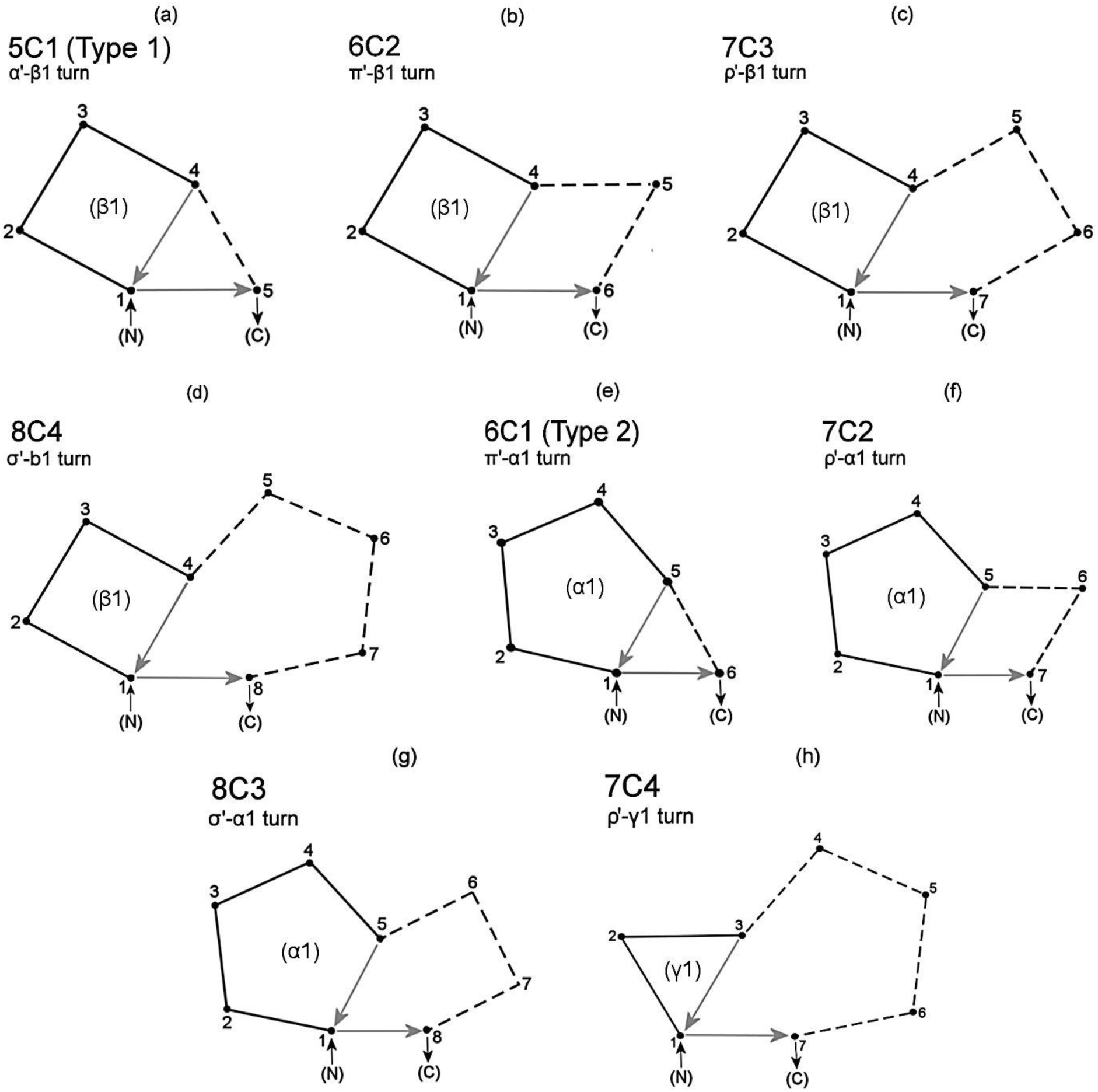
Schematics for eight generalized BBLs with C-terminal bulges. Internal H-bonded turns are drawn with solid lines and bulges are drawn with dashed lines. H-bonds are displayed as arrows oriented in the direction of donation. Loops are labelled using both BBL-specific and compound-turn nomenclatures. To browse a BBL’s structures in ExploreTurns, enter its type name into the *BB H-bonds...* box and click *Load Selected Structures*. BBLs not selected in “baseline” form (with a capitalized framing turn in the romanized CT label) may show additional BB H-bonds. CT labels are listed in Tables S5 and S7 for variants of the BBLs marked with asterisks below. **(a)** 5C1 (type 1) BBL*. **(b-d)** Successive additional insertions into the 5C1’s bulge yield the 6C2-8C4 BBLs*. **(e)** The 6C1 (type 2) BBL* can be formed by the insertion of a residue into the 5C1’s β1-turn, yielding an α1-turn. **(f, g)** Successive insertions of residues into the 6C1’s bulge yield the 7C2 and 8C3 BBLs. **(h)** Conversion of 7C3’s β1-turn to a γ1-turn yields the 7C4 BBL*.

Figure 6 displays example structures for four BBLs with bulges at their N-termini or at both termini, including types {5N1, 6N2, 7N3, 7NC3}. In its hairpin context, the 5N1 BBL can be formed by the insertion of a residue at the loop end of the N-terminal strand of a 2:2 hairpin, while the 6N2 and 7N3 BBLs can result from successive additional insertions into the 5N1 BBL’s bulge. The 7NC3 BBL is classified as both N- and C-type, since it contains split pairs of BB H-bonds at each terminus. Table S6 in Supplementary Section 6 lists the CT labels used to select the BBLs in Figure 6, and includes separate labels for the 5N1 BBL which select or exclude a BB variant (CT label *a’-b2-b’2*) which adds a reverse beta-turn H-bond to the loop; this H-bond is found in about half of all examples of the BBL.

**Figure 6.**
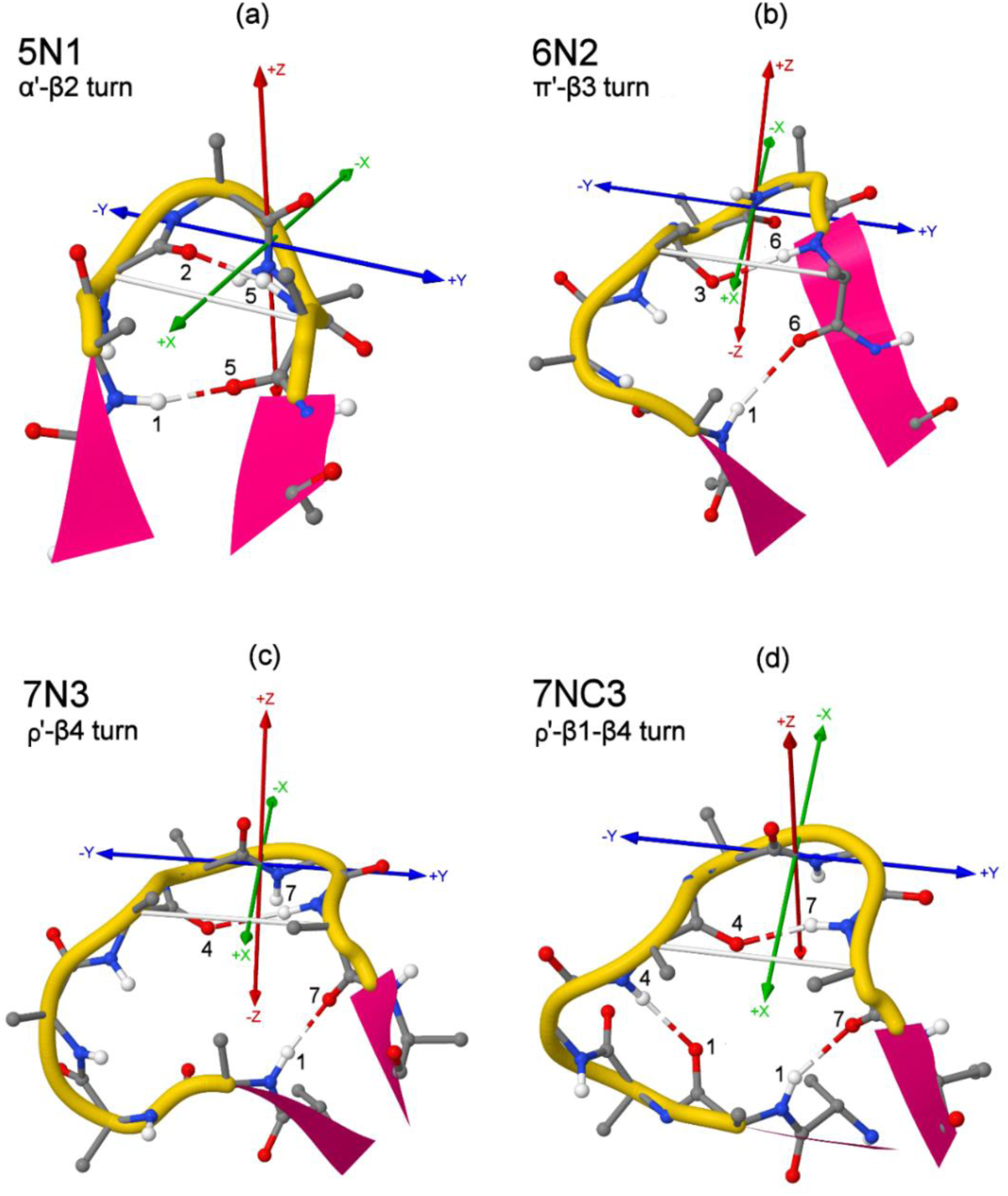
Examples of beta-bulge loops with N-or N/C-terminal bulges. See the Figure 4 caption for loop nomenclature and structure viewing directions. **(a)** In its hairpin context (shown here), the 5N1 BBL can be formed by inserting a residue at the loop end of the N-terminal strand of a 2:2 beta hairpin, splitting the reciprocal pair of beta-ladder H-bonds, but the motif also occurs independently. The baseline motif is shown here, but a common variant (CT label *a’-b2-b’2*) adds a reverse β-turn H-bond. **(b)** The 6N2 BBL adds a residue to 5N1’s bulge. **(c)** The 7N3 BBL adds a residue to 6N2’s bulge. **(d)** The 7NC3 BBL adds a second β-turn H-bond to 7N3. Split pairs of BB H-bonds at both termini classify this BBL as both N- and C-type.

Figure 7 presents schematic diagrams of the structures and H-bond topologies of a larger set of BBLs with N- and N-/C-terminal bulges, including seven additional types with bulge lengths of up to four residues.

**Figure 7.**
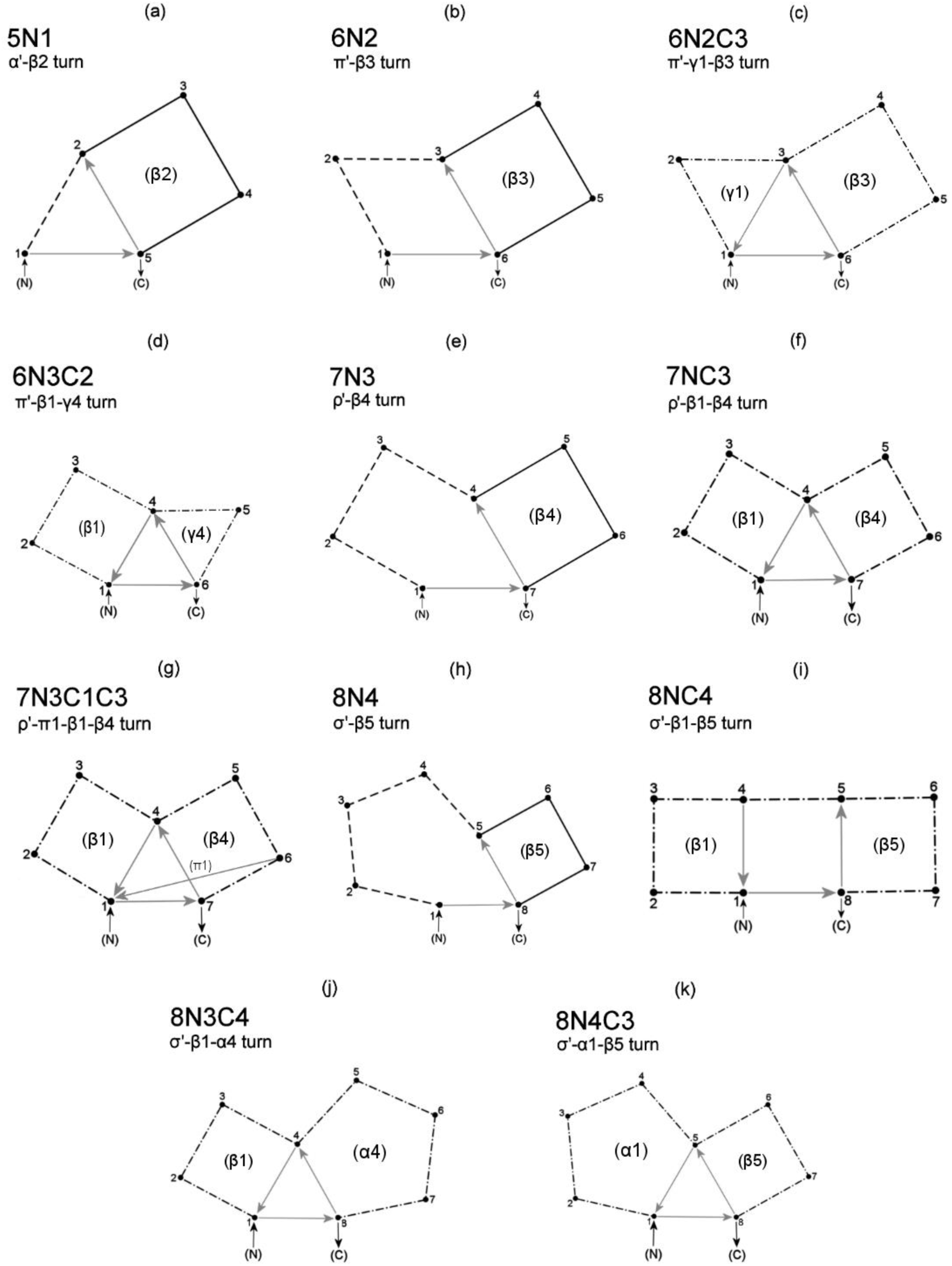
Schematics for eleven generalized BBLs with N- or N/C-terminal bulges. See the Figure 5 caption for display conventions. Dual N/C-type BBLs are drawn entirely with dash-dot lines. Example structures of some BBL types include additional H-bonds; for the BBLs marked with asterisks below, CT labels for BB H-bond or conformational variants are listed in Tables S6 and S7. **(a)** 5N1 BBL*. A common variant (CT label *a’-b2-b’2*) adds a reverse β-turn H-bond. **(b)** The 6N2 BBL* adds a residue to 5N1’s bulge. **(c)** Addition of a γ1 H-bond to 6N2 yields 6N2C3*, with split pairs of BB H-bonds at each terminus that classify it as both N-type, with a two-residue bulge, and C-type with a three-residue bulge. Examples of this motif bind RNA (Figure 8d) and separate DNA strands in the helicase reaction (Figure 8e). **(d)** Addition of a γ4 H-bond to 6C2 (see Figure 4c) yields 6N3C2, classified as both N-type with a three-residue bulge and C-type with a two-residue bulge. **(e)** 7N3 adds a residue to 6N2’s bulge (see DNA-binding example in Figure 8g). **(f)** Addition of a β1 H-bond to 7N3 yields 7NC3, with three-residue N- and C-terminal bulges. **(g)** Addition of a π1 H-bond to 7NC3 yields 7N3C1C3*, with overlapping one- and three-residue C-terminal bulges; the π1 H-bond draws the β-turn “lobes” closer together. DSSP calls 3_10_ helices within the loop in multiple examples of this motif; helices can be excluded with the CT label *r’-p1-b1-b4-\h2*, or the variant *r’-p1-b1-b2-b4* adds a β2 H-bond, yielding helices in all examples. **(h)** 8N4* adds a residue to 7N3’s bulge; the CT label *s’-b*1-b5-\H3* selects a variant which incorporate 3_10_ helices, while the label *s’-b*1-b5-\h3* excludes helices (in each variant, *b*1* excludes 8NC4). (**i**) Addition of a β1 H-bond to 8N4 yields 8NC4 (see DNA-binding example in Figure 8i). **(j)** Conversion of 8NC4’s β5-turn to α4 yields 8N3C4. **(k)** Conversion of 8NC4’s β1-turn to α1 yields 8N4C3.

**Figure 8.**
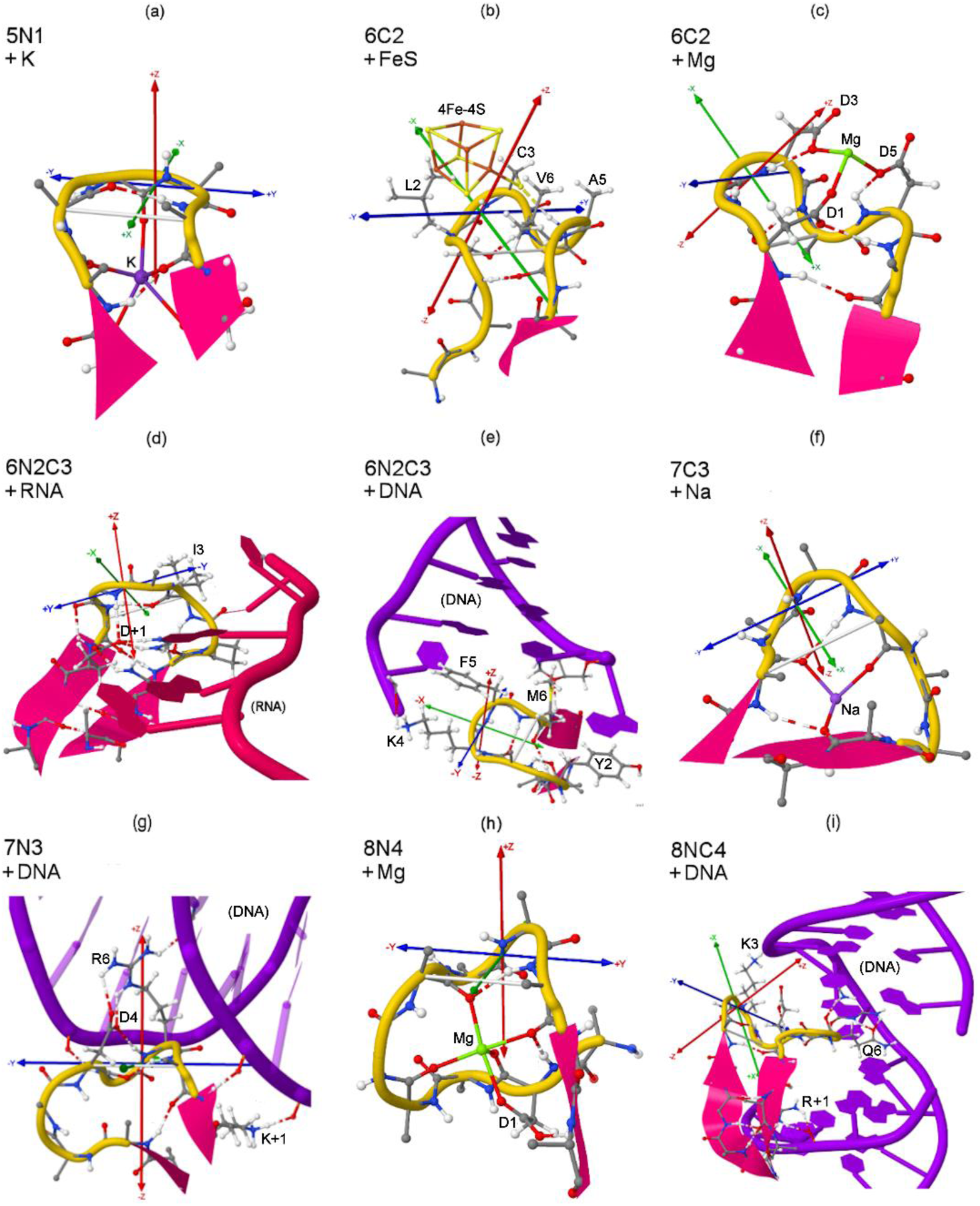
Examples of generalized BBLs at ligand-binding and active sites. Selected SCs are shown, labelled with residue positions in the loop frames. Structures may be viewed by loading their CT labels in ExploreTurns; the SCs specified in the labels serve to uniquely identify the loops in the database, and they may also highlight interactions of interest. **(a)** A 5N1 BBL binds K in the fatty-acid binding protein AtFAP1 (4DOO_A_34, CT label *A’-b2-L1-L2-G3*). **(b)** A 6C2 BBL tethers an active-site 4Fe-4S cluster with Cys3 (C3) in acetyl-CoA synthase/carbon monoxide dehydrogenase. The loop also may help shield the cube from oxidation using its hydrophobic SCs^63^ (1OAO_C_526, *P’-b1-G1-L2-C3*). **(c)** A 6C2 BBL binds Mg with an Asp triplet (D1, D3, D5) at the active site of phosphoglucomutase from Xanthomonas citri (5BMN_A_259, *P’-b1-D1-D3-F4*-D5). **(d)** A 6N2C3 BBL partially “wraps” an RNA base in the HutP antitermination complex, stacking its β3-turn above the base in a likely pi-peptide bond interaction while its γ1-turn forms a BB H-bond with the guanine. Asp (D+1) also H-bonds with the base, and Ile (I3) packs with an adjacent base (1WPU_A_131, *p’-g1-b3-A1-P2*). **(e)** A 6N2C3 BBL at the tip of a “beta wing” motif acts as a “separating knife”, wedging between the first and second base pairs at the blunt end of a DNA duplex in the helicase reaction in human Werner syndrome protein (WRN)^62^ (3AAF_A_1033, *p’-g1-b3-R1*). **(f)** A 7C3 BBL binds Na in methionine R-sulfoxide reductase B1 (3MAO_A_60, *R’-b1-E1-H2*). **(g)** A 7N3 BBL and its hairpin binds DNA with BB H-bonds, a Lys SC following the loop (K+1), and an Asp/Arg salt bridge (D4R6) in RovA, a transcriptional activator of Yersinia (plague) invasin (4AIK_A_81, *r’-b*1-b4-N3-D4-R5-R6*). **(h)** An 8N4 BBL binds Mg with four BB carbonyls and an Asp SC (D1) (4WA0_A_473, *s’-b*1-b5-D1-S5*). **(i)** An 8NC4 BBL and its hairpin bind DNA with BB H-bonds and SCs (K3, Q6, R+1, Y+3) in the loop and following strand in the de novo DNA methyl transferase DNMT3B (5CIY_A_232, *s’-b1-b5-K3-Q6-E8*).

**Figure 9.**
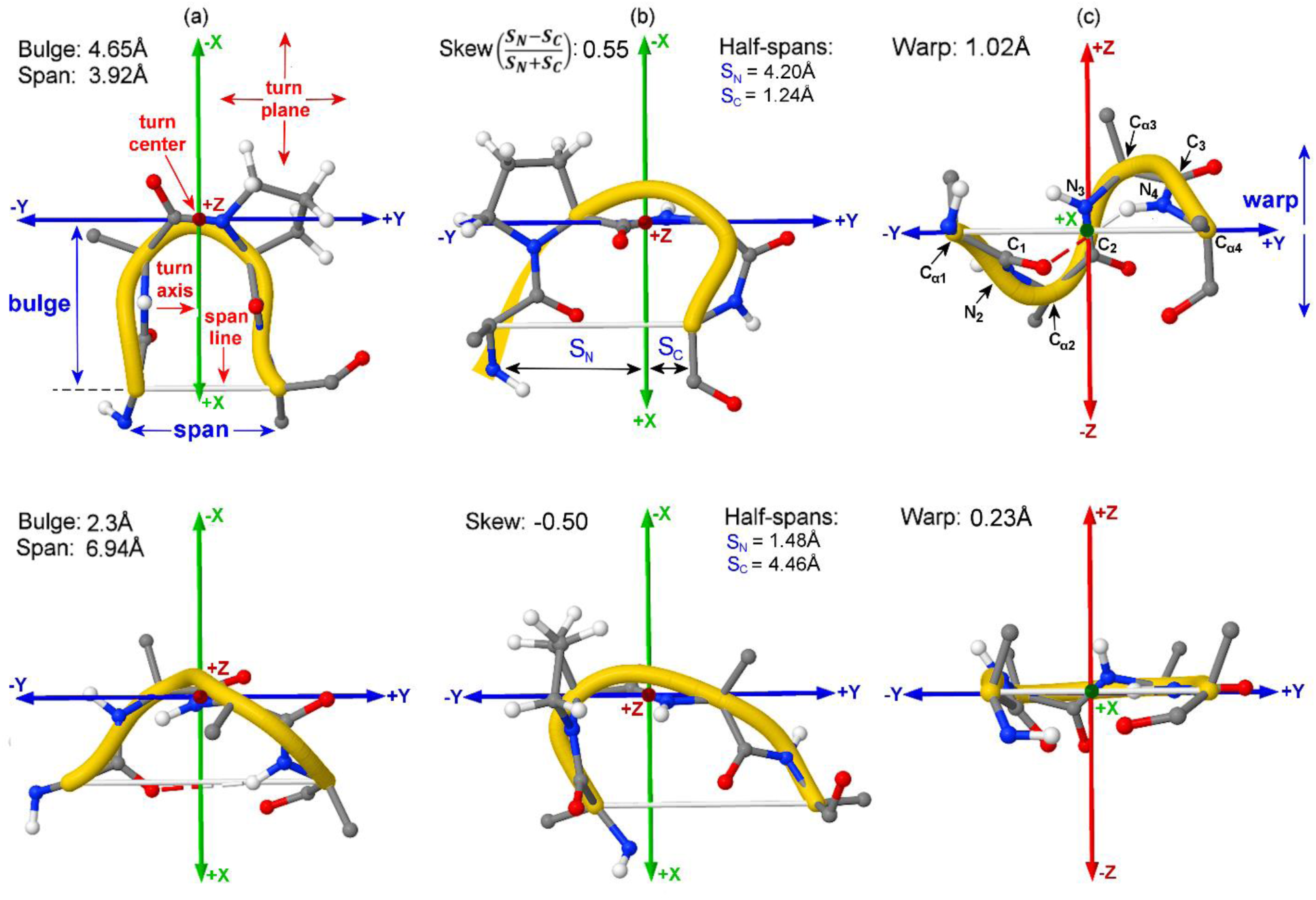
Ranges of the geometric turn descriptors in structures, displayed in the turn-local coordinate system, with geometric definitions labelled in red and descriptors labelled in blue. This figure illustrates the geometric definitions used to establish the turn-local coordinate system, the coordinate system itself, and the geometric descriptors for beta turns. The figure also demonstrates the modes of variation measured by the descriptors, by displaying turn structures with descriptor values near their limits in the dataset. A white bar representing the span line is added to all structures **(a)** 5HKQ_I_74 (top) and 4YZZ_A_239 (bottom) represent the ranges of both bulge and span, since the two descriptors exhibit strong negative correlation. **(b)** 5TEE_A_23 (top), and 3NSW_A_44 (bottom) represent strong positive (C-terminal) and negative (N-terminal) skew. N-and C-terminal half spans are also given. **(c)** 5BV8_A_1285 (top) and 3S9D_B_117 (bottom) represent the limits of warp in classical type I, and demonstrate the wide warp range that occurs within this type. The BB atoms in the main-chain path of the turn between C_α1_ and C_α4_ are labelled; all labelled atoms except C_α1_ and C_α4_ are used to compute warp, as the average distance between these atoms and the turn plane.

#### 3.1.7 Roles of beta-bulge loops

BBLs are hybrid structures that combine the constraints imposed by their defining BB H-bonds with a degree of conformational range configurable by additional BB H-bonds or SC interactions, giving them versatility for structural or functional roles. Figure 8 displays nine examples of generalized BBLs at binding and active sites, with ligands that include metal ions, an FeS cluster, RNA and DNA (x3).

BBLs often form the loop ends of beta-hairpins, and they can be projected by hairpin “arms” into interaction sites, as seen in Figure 8e, where a 6N2C3 BBL at the tip of a “beta wing” motif acts as a “separating knife”, wedging between the first and second base pairs at the blunt end of a DNA duplex in the helicase reaction in human Werner syndrome protein (WRN)^62^ (3AAF_A_1033, CT label *p’-g1-b3-R1*). In another example of BBL projection (Figure 8g), a 7N3 BBL binds DNA with BB H-bonds, a Lys SC, and an Asp/Arg salt bridge in RovA, a transcriptional activator of Yersinia (plague) invasin (4AIK_A_81, *r’-b*1-b4-N3-D4-R5-R6*).

BBLs can bind metal ions for the purposes of catalysis, as in Figure 8c, where a 6C2 BBL binds Mg with an Asp triplet (D1, D3, D5) at the active site of phosphoglucomutase from Xanthomonas citri (5BMN_A_259, *P’-b1-D1-D3-F4*-*D5*). But Figure 8 also shows examples in which metals might be bound in order to help shape or stabilize its structure, as in Figure 8a, where a 5N1 BBL and its hairpin coordinate potassium with 5 covalent bonds in the fatty-acid binding protein AtFAP1 (4DOO_A_34, *A’-b2-L1-L2-G3*), as well as Figure 8f, where A 7C3 BBL coordinates Na with three bonds in methionine R-sulfoxide reductase B1 (the loop is not at the catalytic site) (3MAO_A_60, *R’-b1-E1-H2*). In Figure 8h, an 8N4 BBL is shaped by its coordination of Mg with five covalent bonds as it joins a helix in the “capping” of a beta solenoid in a possible adhesin domain from Caldicellulosiruptor kronotskyensis (4WA0_A_473, *s’-b*1-b5-D1-S5*).

Figure 8b shows another type of interaction between BBLs and metals: a 6C2 BBL tethers an active-site 4Fe-4S cluster with a Cys SC, and it may also “cradle” the cluster with hydrophobic SCs, which could help shield the cube from degradation^63^ (1OAO_C_526, *P’-b1-G1-L2-C3*).

The utility of a BBL’s combination of structural constraint and flexibility is shown to good effect in Figure 8i, in which an 8NC4 BBL and its hairpin bind DNA with BB and SC H-bonds in the de novo DNA methyl transferase DNMT3B. This σ’-β1-β5 turn inserts its β5-turn into the major groove for interaction with the DNA’s bases, while its β1-turn orients at about 90° to the β5-turn, positioning its Lys SC for H-bonding with the DNA backbone from outside the double helix (5CIY_A_232, *s’-b1-b5-K3-Q6-E8*). The versatility of BBLs is also demonstrated in Figure 8d, in which a 6N2C3 BBL (π’-γ1-β3 turn), binding RNA in the HutP antitermination complex, partially “wraps” a guanine base, stacking its β3-turn above the base in a likely pi-peptide bond interaction, while its γ1-turn interacts with the guanine by forming a BB H-bond with the base in its plane (1WPU_A_131, *p’-g1-b3-A1-P2*). These two examples suggest that the H-bonded simple turns in BBLs can impart modularity to the motifs, serving as locally stabilized components that cooperate to accomplish multipart binding or interaction tasks.

### 3.2 Classifying beta hairpins

A beta hairpin can be classified by the length of its chain reversing loop and the H-bond configuration at the loop end of its beta ladder using the nomenclature X:Y, where X indicates the loop’s length according the IUPAC definition of a strand residue and Y is its length according to the DSSP definition^8^. This notation conveys loop length directly and ladder H-bond configuration indirectly, using the contrast between the two conventions. Since the CT nomenclature also describes the length of a loop and the H-bond configuration at its ends (or anywhere in the loop), the notation can also classify hairpins, if it is applied to the hairpin’s chain-reversing turn (the chain-reversing loop inclusive of the pair of H-bonded residues at the loop end of the hairpin’s beta ladder). Since all hairpins classified in the X:Y nomenclature exhibit either a single reverse H-bond or a pair of reciprocal normal and reverse H-bonds at the loop ends of their ladders, all X:Y hairpins with loop lengths up to 10 residues (corresponding to a chain-reversing turn length of 12 residues, the longest covered by the CT notation) are classifiable with CT labels composed of either a single entry for a reverse H-bonded turn, or a pair of entries for normal and reverse turns of the same type, so that the X:Y hairpin types {1:1, 1:3, 2:2, 2:4, 3:3, 3:5,…} become {γ-γ’, γ’, β-β’, β’, α-α’, α’,…} in the CT hairpin notation.

The CT hairpin notation has the disadvantage, compared to the X:Y system, that it requires knowledge of the lengths of the simple turn types. But it has the advantage that it directly specifies H-bonds rather than relying on the contrast between two conflicting conventions (IUPAC and DSSP), and more importantly, the CT notation is more complete, since it describes the H-bond configuration in the hairpin’s chain-reversing loop as well as its ladder. For example, the common structure described in the X:Y notation as a “3:5 hairpin with a type 1 beta-bulge loop” is simply an “α’-β1 hairpin” in the CT hairpin notation, while a “4:4 hairpin with a Schellman loop” is a “π-π’-β2 hairpin”.

The ExploreTurns loop motif browser uses preset CT labels to select sets of examples of 16 types of beta hairpins from the four classes^6^ in “baseline” configuration, with no BB H-bonds in their chain-reversing loops, and the browser includes hairpin schematics and classifies the structures using both the X:Y and CT hairpin notations. Each hairpin schematic is also annotated with the CT label used to select the hairpin, which must specify residues N- and C-terminal to the chain-reversing turn as well as the residues within the turn, in order to specify the required BB H-bonds in the hairpin’s beta ladder. A shorthand CT hairpin notation that automatically embeds a CT label entered by the user into an extended label that includes the required additional beta-ladder H-bonds (e. g. “a’-b1 hairpin” entered by the user is translated to the CT label “t-t’-a’3-b3”, which is then used to select structures) is not yet live in the ExploreTurns CT interpreter; this is a priority for a future ExploreTurns release, with a structural window longer than 12 residues, which can better support the selection of hairpins with longer chain-reversing loops.

### 3.3 General structural analysis of beta turns and their contexts

In addition to extensive support for the exploration and analysis of H-bonded loop motifs via CT labels, ExploreTurns supports the analysis of the structures of beta turns and their contexts in general, via selection criteria that describe the beta turn at the center of the 12-residue ExploreTurns window or specify structure in the window outside the turn (see the feature summary and user guide). The tool has been applied to studies of SC-mediated helix N-capping (Asx N-caps), BB-mediated helix C-capping (Schellman loops), and the relationship between beta-turn BB geometry and depth beneath the solvent-accessible surface.

#### 3.3.1 Mapping Asx N-cap sequence preference vs. cap geometry

At the helical N-terminus, unsatisfied BB NH groups are commonly “capped” by H-bonding with the SCs of Asp or Asn (Asx) or Ser or Thr (ST) in Asx/ST N-caps^14,15,26,27^. Beta turns are also common at the helical N-terminus^15^, and the Asx AAs in helix-terminal turns show greatest overrepresentation when they occur at the third turn position and this position coincides with the NCap residue at the start of the helix. Figures S7 and S8 in Supplementary Section 7 apply the sequence motif, DSSP secondary structure, and “context vector” selection features of ExploreTurns to heat-map the sequence preferences of Asx N-cap motifs in beta turns vs. helix/turn geometry. The maps reveal the cap geometries most compatible with these motifs, and they can be used to guide ExploreTurns investigations of the structure and H-bonding associated with Asx N-caps formed by beta turns.

#### 3.3.2 Profiling Schellman loop/beta-turn conformations

Both Schellman loops and beta turns commonly occur at the helix C-terminus, where they overlap to differing degrees in conformations that depend on both the loop’s registration in the turn’s frame and the turn BB geometry, which determines the orientations of the BB NH groups in the turn which donate the loop’s capping H-bonds. Supplementary Section 8 applies the classical type/BB cluster, DSSP secondary structure, and BB H-bond selection features of ExploreTurns to profile the geometries of Schellman loop/beta-turn combinations at the C-termini of alpha helices. Four principal conformations are identified, and Figure S9 presents example structures and ExploreTurns selection criteria for each.

#### 3.3.3 Investigating the depth dependence of turn geometry

The extent to which beta-turn BB geometry is affected by interactions with adjacent structure in a protein is an open question. If turn geometry is substantially influenced by neighboring structures, then the distribution of turns across BB geometries might be expected to change as the turns become more buried in the protein. Supplementary Section 9 applies the depth-selection and type-profiling features of ExploreTurns to profile the changes that occur in the distribution of beta-turn geometries as depth increases. The proportion of structurally unclassified (type IV) turns is found to increase by more than 50% with depth at the expense of classical type I, reaching almost half of all turns (Figure S10). The apparent transfer of a substantial fraction of all turns, as depth increases, from type I to the structurally unclassified category may reflect geometric distortions due to increasing and/or more impactful contacts with adjacent components of the protein, increasing stress transmitted through the BB by neighboring structures or structures that incorporate beta turns, or increased binding of ligands or prosthetic groups by turns, and it supports the hypothesis that many type IV turns represent structural distortions rather than intrinsically stable turn conformations^45^.

## 4 Discussion

ExploreTurns supports the study of beta turns and most loop structure with multiple new features, including a turn-local coordinate system which provides a useful framework for the analysis of turns and their contexts, a global turn alignment that supports structure comparisons, geometric descriptors for beta turns that complement the Ramachandran-space classification systems, and new nomenclatures that classify H-bonded loop motifs, beta hairpins and BBLs and support the exploration of local backbone structures and the detection of new structures. The compound turns characterized here represent significant new contexts for beta turns, and although many of them are relatively rare in the redundancy-screened dataset, they can nevertheless play key roles, as demonstrated in Figure 8.

The large family of generalized BBLs, which demonstrate a range of geometries and H-bond topologies, could provide models for loop design. BBLs can play structural, regulatory, binding or catalytic roles (Figures 4b, 8a-c, 8f, 8h), and they commonly constitute the chain-reversing loops of beta hairpins, so they may prove particularly useful in the design of these motifs, which are popular components of engineered structures because they play key roles in a wide range of bioactive molecules, from antimicrobial peptides to beta-barrel transporter proteins^7^ (e. g. the 7C3 BBL, CT label *R’-b1-E1-*S3, which forms the L3 hairpin loop in the P. aeruginosa alginate transporter AlgE). BBLs may find application as configurable “tools” at the ends of beta-ladder “arms” that project them into interaction sites, as seen with the 6N2C3 and 7N3 BBLs that interact with DNA in Figures 8e and 8g (CT labels *p’-g1-b3-R1* and *r’-b*1-b4-N3-D4-R5-R6*), or as accessories to beta sheets that support binding or interaction at the sheet edges, as seen in Figure 8i (CT label *s’-b1-b5-K3-Q6-E8*).

CTs are already employed in design; for example, the beta-turn motif used in a recent design of “buttressed” loops for function^64^ was a β-β’ turn (extended CT label S*-s’*-b3-b’3) extracted from a structural database, and as design efforts increasingly encompass detailed, nonrepetitive BB structures tailored to particular contexts and functions, the availability of a library of versatile loop motifs may prove useful. Design efforts involving CTs could focus not only on employing existing motifs, but also on designing new loops; motifs of up to ten residues, constrained by up to five BB H-bonds, are described here, but design work could probe the limits of the size and complexity of local H-bonded BB structures.

A reservoir of new examples of compound turns which could provide much more complete coverage of the BB and SC variants of the CTs described here (and may yield new motifs) might already be available, in the form of the large sets of protein structures predicted from sequence by AlphaFold 2^65^ and ESMFold^66^. AlphaFold 2 has shown good accuracy in the prediction of short loops^67^, supporting this prospect, but the rarity and/or complexity of some CTs (e.g. the 7N3C1C3 BBL in Figure 7g, the crown pyramid^60^ in Figure 3k, or the double Schellman loop in Table S4, CT label t*-t’*-p1-b2-p4-b5_*) may challenge prediction tools. If all tools cannot accurately predict all loops, a set of loops could be used in benchmark testing to rank the tools by their ability to predict irregular H-bonded motifs, and this CT library could support the development of improvements in the prediction of local structures. The conformations of any CTs that cannot be accurately predicted could be further explored with computational modelling like that applied by Venkatachalam to beta turns^2^, and the structures and energetics of all the newly-identified loops could be investigated experimentally using combinatorial mutagenesis and other methods.

Other potential applications of the CT nomenclature and the motif library presented here include structure annotation, structure selection, protein classification, and the study of protein evolution. The nomenclature could be applied to catalog the H-bonded loop motifs in all protein structures, providing a detailed picture of structured loops which could be added to tools such as the PDB’s secondary structure annotator. The CT interpreter could be incorporated into the front-ends of structural databases to support the selection of loop motifs and the beta hairpins that incorporate them. Finally, CT profiles could be computed for protein families classified by structure or function, and these profiles could shed light on the relationships between local H-bonded structure and fold or function as well as the evolution of loop motifs.

## 5 Methods

### 5.1 The ExploreTurns database

Beta turns are defined here as four-residue BB segments with a distance of no more than 7 Å between the alpha carbons of the first and fourth residues and central residues with DSSP codes outside the set {H, G, I, E} that specifies helix or strand^3^. Turn/tail structures were extracted from their PDB files^49^ with the aid of a list of peptide chains with a maximum mutual identity of 25% obtained from PISCES^68^. Since sequence motif detection identified multiple motif “artifacts” generated by remaining local redundancy in the data, an additional step of turn-local redundancy screening was applied: structures were clustered and filtered to reduce redundancy within five residues of turns to 40% or less; this threshold was chosen because it effectively reduced artifacts while preserving a dataset of sufficient size.

Turn structures were screened for quality by requiring that there be no missing residues within 5 residues of any turn and excluding structures with bond lengths that constituted outliers from established values. The final dataset of 102,192 structures was compiled with resolution and R-value cutoffs of 2.0 Å and 0.25 respectively; average values in the dataset are 1.6 Å and .20.

### 5.2 Turn-local coordinate system and alignment

A common local Euclidean-space coordinate system for beta turns was established via a set of geometric definitions^48^ (see Figure 9). The *span line* is the line between a turn’s first and last alpha carbons (*C*_*⍺*1_and *C*_*⍺*4_), and the span, already used in the beta-turn definition, is its length. The *turn center* is the midpoint of (*C*_2_→*N*_3_), the middle peptide bond in the turn. The *turn plane* contains the span line and passes through the turn center, and the perpendicular dropped from the turn center to the span line defines the *turn axis*. Using these definitions, a local, orthogonal right-handed coordinate system is established. The system’s origin lies at the turn center, while the x axis lies in the turn plane, collinear with the turn axis, with its positive sense oriented from the center towards the span line. The y axis lies in the turn plane, parallel to the span, with its positive sense oriented towards the C-terminal half of the turn. Finally, the z axis is perpendicular to the turn plane, with its positive sense determined by the cross-product of the x and y axes.

The atomic coordinates of each turn are transformed from the global protein system in the turn’s PDB file to the turn-local system, implicitly aligning the structures. The alignment’s accuracy was tested by a comparison between implicit pairwise turn alignments and the best alignments generated by an RMSD-based perturbative search procedure. In this process, the average distances between the corresponding BB atoms of randomly chosen, implicitly aligned turn pairs were compared to the distances obtained for the best alignments generated by a perturbative search applied to the pairs which seeks the minimum average interatomic distances by introducing small variations in the relative position and orientation of the two turns. This procedure was applied to a random sample of 100 million turn pairs to generate an estimate of the average positional error of the implicit alignments. Since this estimate (0.17 Å) is comparable to the errors expected for the positions of well-determined atoms in well-refined structures (0.1 Å - 0.2 Å; overall RMSDs for independently determined 2 Å structures have been measured as 0.5 Å - 0.8 Å)^69^, the accuracy of the implicitly established alignment was judged to be satisfactory.

### 5.3 Geometric turn descriptors

Geometric descriptors for beta turns^48^ are defined here and illustrated in Figure 9.

#### 5.3.1 Span and half-spans

The pre-existing span descriptor measures the distance [Å] between *C*_*⍺*1_and *C*_*⍺*4_in the turn, and a 7 Å threshold is part of the beta-turn definition. Two new span-related descriptors are defined: the N- and C-terminal half-spans measure the distances between the turn axis and *C*_*⍺*1_ and *C*_*⍺*4_ respectively, and quantify the separate “widths” of the N- and C-terminal halves of the turn, providing an additional degree of freedom in the measurement of span. Span values in the dataset range from a minimum of 3.13 Å to the 7.00 Å limit set by the beta-turn definition, while N-terminal half-spans range from 0.00 to 4.76 Å, and C-terminal half-spans from 0.76 Å to 4.82 Å.

#### 5.3.2 Bulge

Bulge measures the magnitude of the excursion that the BB makes from the turn span as it bends to form the turn. Bulge is defined as the distance [Å] from the turn center to the span line. Bulge values in the dataset range from 1.28 Å to 4.70 Å.

#### 5.3.3 Aspect

Aspect, which approximates the aspect ratio of the projection of the turn’s BB onto the turn plane, is computed as the ratio of bulge to span.

#### 5.3.4 Skew

Skew is a dimensionless measure of the asymmetry of the projection of a beta turn’s BB onto the turn plane. Skew is defined as the directed displacement along the span line between the line’s center and the point of intersection between the turn axis and the line, divided by half the span. Skew is negative when the axis intersects the span line in its N-terminal half, indicating a “lean” in the N-terminal direction, or positive when the axis intersects the line in its C-terminal half, producing a C-terminal lean. Skew is expressed in terms of the N- and C-terminal half-spans (*S*_*N*_ and *S*_*C*_) as:

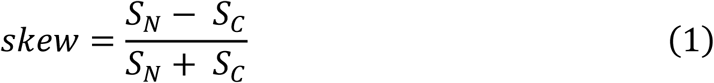

Skew is positive when *S*_*N*_ is larger than *S*_*C*_, negative when the reverse is true, or zero when the half-spans are equal. A turn that skews so strongly N-terminally that its axis intersects the span line at the position of *C*_*⍺*1_(*S*_*N*_= 0) would exhibit a skew of -1, while a turn skewed C-terminally so that its axis intersects the line at *C*_*⍺*4_(*S*_*C*_ = 0) would show a skew of +1. Skew values in the dataset range from -1.00 to 0.78.

#### 5.3.5 Warp

Warp measures the departure from flatness of a beta turn’s BB in the z dimension of the turn coordinate system, perpendicular to the turn plane. Warp [Å] is defined as the average excursion of the turn’s BB above or below the turn plane between *C*_*⍺*1_ and *C*_*⍺*4_, excluding BB carbonyl oxygen atoms as well as *C*_*⍺*1_and *C*_*⍺*4_themselves (since they lie within the plane).

Warp is computed as the average absolute value of the z-components of the BB atoms (C_1_, N_2_, C_*⍺*2_, C_2_, N_3_, C_*⍺*3_, C_3_, N_4_). Warp values in the dataset range from 0.14 Å to 1.25 Å.

### 5.4 The compound-turn nomenclature for loop classification

The compound-turn (CT) nomenclature treats all short H-bonded loop motifs as overlapping H-bonded turns, and classifies them by the types (lengths) of these component turns and the start positions of the turns in the loops. H-bonded loop motifs are classified by CT labels composed of entries separated by dashes. The first entry in a label, which indicates the length of the “framing” turn that embodies the entire loop, has the form:

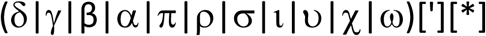

where the parentheses are used here for grouping and the brackets indicate optional components; neither punctuation appears in the notation. The Greek letters, separated by ‘OR’ bars here, indicate the component turn types, which have lengths of 2-12 residues. The single-quote character is used when a component turn’s defining H-bond is donated in the N→C direction, indicating a “reverse” turn, rather than the default C→N direction seen in “normal” turns^1^, while ‘*’ is used when the turn’s defining H-bond is to be excluded from the loop rather than included. If a CT’s label contains entries which exclude both normal and reverse H-bonds in its framing turn (e.g. π*-π’*), then it describes an “open” loop, with termini not linked by a single shared H-bond.

Subsequent dash-separated entries in the CT label specify the loop’s internal H-bonded turns (which may share a terminus with the framing turn), using the form:

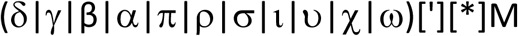

which adds M, an integer that specifies a component turn’s start position in the loop.

Constraints that require positive or negative values for the (φ, ψ) dihedral angles of a CT’s residues may be appended to its dash-separated list of component-turn specifiers, using the forms:

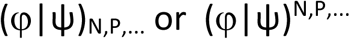

where the subscripts constrain the dihedral angles of loop residues (N, P, …) to negative values, while the superscripts constrain the angles to positive values. Dihedral constraints are concatenated to the end of the list of component-turn entries in a label without dashes or spaces.

A CT’s BB conformation may also be specified using the “right-handed” and “left-handed” α/β categories {αR, αL, βR, βL}, which provide a useful simplification of Ramachandran space and support (for example) the approximate but effective specification of the types of β-turns within CTs (click *β-Turn Types* to see the α/β categories of classical turn types {I, I’, II, II’} as well as the category definitions; the definitions are also given in Supplementary Section 11). The α/β conformational constraints take the form:

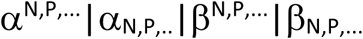

where superscript(s) indicate the loop position(s) which exhibit the right-handed form of the conformation, while subscripts(s) indicate position(s) with the left-handed form. Like (φ, ψ) constraints, α/β conformations are concatenated to the CT label without dashes or spaces.

To avoid confusion with the names of supersecondary structure motifs (e.g. βαβ), a compound-turn label is always followed by the word “turn”.

### 5.5 The ExploreTurns CT interpreter

ExploreTurns includes a CT interpreter which adds features to the notation that facilitate structure selection and searching. A CT label is entered into the *BB H-bonds/Loop motif/Compound turn* box as a list of dash-separated entries, after translation from the Greek alphabet using the Beta Code^59^ romanization (see examples in Table 1). The entry for a loop’s framing turn takes the form:

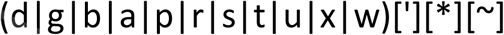

where the parentheses, brackets and “OR” bars are not entered. The single-quote character specifies a ‘reverse’, N→C H-bonded turn, while the asterisk excludes the specified normal or reverse turn. The tilde neither requires nor excludes an H-bond in the turn; a framing turn entered with a tilde serves only to specify the length of the selected BB segment, and may be used to extend the CT label without indicating H-bond configuration, for the purpose of profiling, selecting, or excluding structure in a motif’s context (see Section 5.6 and examples 5 and 6 in *H-bonded loop motifs* in the feature summary). The framing turn may be capitalized to specify a “baseline” loop (see below).

The entries for a loop’s internal H-bonded turns take the form:

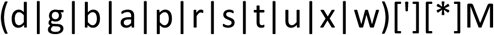

The integer M, which specifies the start position of a component turn in the loop, is required for internal turns, which must be specified in lowercase. The tilde option is omitted for internal turns, since it would serve no purpose.

The CT interpreter supports the selection of loops that contain particular sequence motifs, which are defined as the requirement for one or more specific AAs at one or more loop positions. An AA is specified at loop position M with an entry of the form:

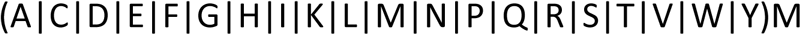

Any α/β or (φ, ψ) conformational constraints in a CT label are translated, using the romanizations (α, β) → (A, B) and (φ, ψ) → (F, Y), into dash-separated entries of the form:

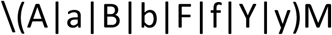

where uppercase α/β entries indicate the corresponding right-handed conformations and lowercase left-handed, while uppercase (φ, ψ) entries indicate positive dihedral angles and lowercase negative, at loop position M. The backslash is required as an “escape” character which signals the tool not to interpret a letter as an AA.

Optional, escaped dash-separated entries of the form:

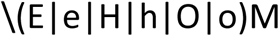

apply secondary-structure conditions to loop residues, which either require (uppercase) or exclude (lowercase) residues of strand type (E/e), helix type (H/h, for helices of any type), or any repetitive (strand or helix, “ordered”) structure (O/o) at loop position M.

Capitalization of the letter specifying the first (framing) turn in a CT specifies the loop in its strict baseline form, without additional BB H-bonds either within the loop or linking loop residues to any other residue in the 12-reside ExploreTurns window. An alternative, more permissive baseline loop definition, which only excludes structures with additional BB H-bonds that occur entirely within the loop, may be specified by prepending ‘@’ to the CT label. If a loop is commonly in contact with other secondary structure, the choice of baseline format can have a large impact on the number of selected structures: the Schellman loop, for example, frequently caps helix C-termini and receives additional H-bonds from the helix, and the strict baseline CT label “P-b2” reduces the size of the selected set by 73% compared to the unrestricted label “p-b2”, while the permissive baseline label “@p-b2” reduces the set size by just 24%.

Loop variants with additional BB H-bonds can be ruled out more selectively using entries of the form:

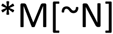

which exclude examples with additional BB H-bonds donated or received at loop position M, or (when ∼N is included) within the position range from M to N inclusive.

CTs can occur at any position within the 12-residue ExploreTurns window, which is always centered on a beta turn. By default, a CT label selects the structures found in the loop’s most common position frame, but alternative frames can be selected by appending the suffix:

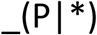

to the label to select either the set of structures from the frame ranked P in the list of frames ordered by loop abundance, or (with “_*”) select the merged set of unique loop structures from all frames (the same structures can occur in multiple frames due to overlapping turns/tails). Alternative position frames show the loop in different structural contexts, or (if a loop contains overlapping beta turns) they may be used to select loop examples that are oriented in a more natural way in the turn-local coordinate system. Note that all types of entries included in a loop’s CT label can affect the ordering of a loop’s position frames, and may shift the default frame.

A loop’s position in the ExploreTurns window is indicated in the text displayed above the *BB H-bonds/Loop motif/Compound turn* box after the loop is loaded, in the format “M→N (P)”, where M and N indicate the start and end positions of the loop in the 12-residue window centered on a beta turn, and P indicates the loop’s offset from the central β-turn. If structures are selected in a single frame, then the loop’s position is the same for all structures in the set, but if the merged set of all structures is selected (with “_*”), then the position is in general different for each structure, and the text above the *BB H-bonds…* box updates as each loop example is loaded.

The complete, romanized CT notation for entry into ExploreTurns is summarized, using single entries of each type, as:

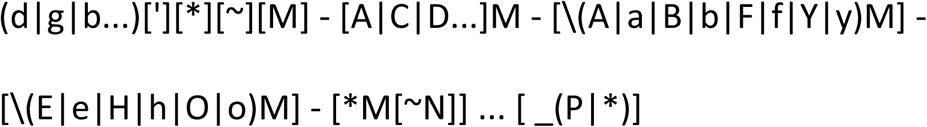

where the parentheses, brackets and bars are not entered, and white space is optional. The residue position specifier M is omitted from the entry for the framing turn (which must be listed first), but all other entries (except the frame-selection suffix) must include M. Tilde is an option only for the framing turn. Capitalization of the framing turn applies the strict baseline format, while prepending ‘@’ specifies the permissive form. If only a framing turn is specified, then a “simple” H-bonded turn is selected (note that the longer types of simple turns commonly incorporate additional internal H-bonds). Apart from the entry that specifies the framing turn and the frame-selection suffix, entries may be listed in any order, but a convention followed here is that internal component turns are ordered by increasing position in the loop followed by decreasing turn length, so that a β2-turn is listed before a β3-turn, which is itself listed before a (shorter) γ3-turn.

### 5.6 Extending CT labels

CT labels specify structure only within a loop’s framing turn, and when a loop is loaded all selection criteria must be entered within the label, since criteria entered in input boxes other than the *BB H-bonds…* box refer to positions in the 12-residue ExploreTurns window, and the registration of a CT in this window is not known in advance. Although other ExploreTurns input boxes may not be used with CTs, structure outside a motif can be specified or excluded by extending the CT label to cover neighboring residues. This technique can be used to search for variants with additional BB H-bonds that extend the structure of a given loop. For example, many 5C1 (type 1) BBLs (α’-β1 turns) are embedded within nine-residue H-bonded ι-turns, since these BBLs commonly occur in the chain-reversing loops of 3:5 beta hairpins, and the ι-turn in the hairpin’s beta-ladder encloses the BBL. However, since 5C1 BBLs enclosed by ι-turns also occur independently of beta hairpins, the loop which includes the BBL within the turn qualifies as a structural motif in its own right, and the extended CT label “t-a’3-b3-\e1-\e2-\e8-\e9” adds two residues to each end of the BBL, selecting this motif and specifying that its termini do not overlap a hairpin’s beta ladder. Other loop motifs can also be extended at both termini in this way, including the 6C1 (type 2) and 6C2 BBLs and the Schellman loop, or loops may be extended at just one terminus, as in the Schellman+ρ or α+Schellman motifs (see the loop browser).

CT labels may also be extended to select loops that exclude particular BB H-bonds in a motif’s BB context, or the ‘∼’ option for the framing turn may be applied to extend the loop without regard for H-bond configuration, for the purpose of profiling, selecting or excluding DSSP secondary structure in a motif’s context, or selecting sequence motifs or BB conformation in the context (see examples 5 and 6 in *H-bonded loop motifs* in the feature summary).

### 5.7 Exploring BB and SC variants of CTs

Two ExploreTurns display features support the exploration of CT variants. When a loop is loaded with a CT label that does not specify a baseline mode, all H-bonded component turns that occur in the set of loop structures are listed in the *H-bond profile* box, and turns not already specified in the loop’s definition may be copied into the CT label to investigate the structures of the motif’s BB H-bond variants (see *H-bonded loop motifs/Examples/Ex. 3* in the feature summary). In addition, ranked statistics for the single-AA and pair sequence motifs that occur in the loop are displayed at the top of the profile window when a CT is loaded, and sequence motif labels can be copied from the window into the CT label to investigate the loop’s SC structures and their effects on its BB geometry (see *H-bonded loop motifs/Examples/Ex. 4*).

### 5.8 CT examples

Table 1 provides a set of CT labels that demonstrate the features of the notation and its interpreter. Labels are given in Greek form for classification as well as romanized form for entry into ExploreTurns. To load a set of motif structures, copy a romanized CT label into the *BB H-bonds/Loop motif/Compound turn* box and click *Load Selected structures*.

### 5.9 Beta-bulge loop nomenclature

Since beta-bulge loops (BBLs) have a characteristic general form (Section 3.1.6), they can be classified with a motif-specific notation in addition to the CT nomenclature. BBL types are labelled in the three-component format:

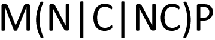

where M specifies the overall length of the loop, {N, C, NC} specify the location of the bulge(s) at the N-terminus, C-terminus, or both (for loops which have split pairs of BB H-bonds at both termini), and P specifies bulge length (the number of residues added to form the bulge, by splitting a reciprocal pair of BB H-bonds). If an NC-type BBL has bulges of different lengths (e.g. 6N2C3), or a bulge defined by an overlapping pair of split BB H-bonds (e. g. 7N3C1C3), then the length of each bulge is given separately.

### 5.10 H-bond definition

BB H-bonds are defined by both electrostatic (dipole-dipole) energy and geometric criteria. The electrostatic energy is computed as:

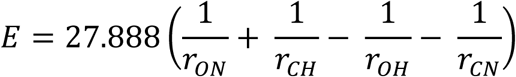

and the DSSP^50^ threshold of -.5 kcal/mol is applied, while the geometric criteria specify a maximum distance between the hydrogen atom donated by the BB nitrogen and its carbonyl oxygen receptor (H-bond length) of 2.6Å, a minimum nitrogen→hydrogen→oxygen angle (H-bond angle) of 100°, and a minimum hydrogen→oxygen→carbonyl carbon angle (H-bond acceptor angle) of 90°.

### 5.11 Statistical methods

The methods used to compute sequence motif overrepresentation and p-value^15,48,70,71^ are described in Supplementary Section 10.

### 5.12 Languages and tools used

Biopython^72^ was used to extract the structural data from PDB^49^ files. The depths of beta turns beneath the protein’s solvent-accessible surface were computed with Depth^73^. ExploreTurns was written in HTML/CSS and Javascript, with an embedded JSmol viewer^74^. The tool is tested for compatibility with the Chrome, Edge, and Firefox browsers; for best performance, browse with Chrome or Edge.

## Supporting information

ExploreTurns supplementary sections

## Declarations

### Availability of data and materials

ExploreTurns is available at www.betaturn.com. All data supporting the conclusions of this article are available from the Protein Data Bank at www.rcsb.org.

### Competing interests

The author declares that he has no competing interests.

## Funding

This work was funded entirely by the author.

## Acknowledgements

The author wishes to thank Athena Newell for her work as a research associate and Kit Newell for his work as a research assistant. The constructive suggestions of the attendees of ISMB/3DSig 2019 (Basel) and the 2019 and 2023 Annual Symposia of the Protein Society (Seattle and Boston) also contributed to the development of this project.

## Notes

https://www.betaturn.com

### Competing Interest Statement

The authors have declared no competing interest.

### Summary of Updates

Updates to the text, including more extensive treatment of the analysis of Euclidean-span beta-turn geometry with geometric descriptors. New features added the tool, including the ability to select alpha/beta BB conformations within H-bonded loop motifs and DSSP profiles of loop motifs. New loop motifs identified, including a "double" Schellman loop.

