## Supplementary material for "ExploreTurns: A web tool for the exploration, analysis, and classification of beta turns and structured loops in proteins; application to beta-bulge and Schellman loops, Asx helix caps, beta hairpins and other hydrogen-bonded motifs": ExploreTurns supplementary sections

#### Supplementary sections


1 | Exploring beta-turn geometry with Euclidean-space descriptors 2

Figure S1. Distributions of the geometric descriptors in the classical turn types 3

Table S1. Selecting turns with geometric descriptors near their limits 4

2 | Tuning turn geometry for compatibility with SC motifs 5

Figure S2. Sequence motif overrepresentation plots for single-AA motifs 6

Figure S3. Sequence motif overrepresentation plots for pair motifs 7

Table S2. Selecting sequence motifs tuned by warp 8

3 | Ramachandran-space descriptor heatmaps 9

Figure S4. Examples of heatmaps 10

4 | Selecting supersecondary structures with context vectors 11

Table S3. Selection criteria for examples of supersecondary structures 11

Figure S5. Examples of supersecondary structures 12

5 | Classifying and exploring non-BBL H-bonded loop motifs and variants 13

Table S4. Selection criteria for non-BBL H-bonded loops and variants 14

6 | Classifying and exploring beta-bulge loops 15

Table S5. Beta-bulge loops with C-terminal bulges 15

Table S6. Beta-bulge loops with N- or N/C-terminal bulges 16

Table S7. Additional beta-bulge loops and variants 17

7 | Mapping Asx N-cap sequence preference vs. cap geometry 18

Figure S6. Example of an Asp3 helix N-cap 19

Figure S7. Mapping Asp3 N-cap sequence preference vs. cap geometry 20

Figure S8. Mapping Asn3 N-cap sequence preference vs. cap geometry 21

8 | Profiling Schellman loop/beta turn conformations at alpha-helical C-termini 22

Figure S9. Principal conformations of Schellman loop/beta turn combinations at alpha-helical C-termini 23

9 | Investigating the depth dependence of beta-turn geometry 24

Figure S10. Variation of the classical type distribution with depth 25

10 | Computing sequence motif overrepresentation and p-value 26

11 | Definitions of the $\alpha/\beta$ conformations 29

Table S8. Dihedral-angle ranges for the $\alpha/\beta$ conformations 29

References 30

### 1 | Exploring beta-turn geometry with Euclidean-space descriptors

Geometric turn descriptors[1,2] (see definitions in Section 5.3 of the main paper) complement the Ramachandran-space turn classification systems, enabling Euclidean-space structural discrimination within the classical turn types and BB clusters. ExploreTurns provides descriptor distribution plots (interpolated histograms) which support the exploration of turn BB geometry in Euclidean space, either in the global turn set or within each type or cluster. Figure S1 displays the descriptor plots for all classical types superimposed; to open a plot for any individual type or cluster, click the ***Distbn*** button in the descriptor's column after entering a type or cluster name in the ***β-turn type/cluster*** box (if this box is left blank, the descriptor's distribution in the global set of turns is displayed).

Figure 9 in Section 5 of the main paper uses the structures of turns with descriptor values near their limits to illustrate the descriptor definitions and demonstrate the modes of conformational variation which they characterize; Table S1 provides selection criteria for examples of these turns.

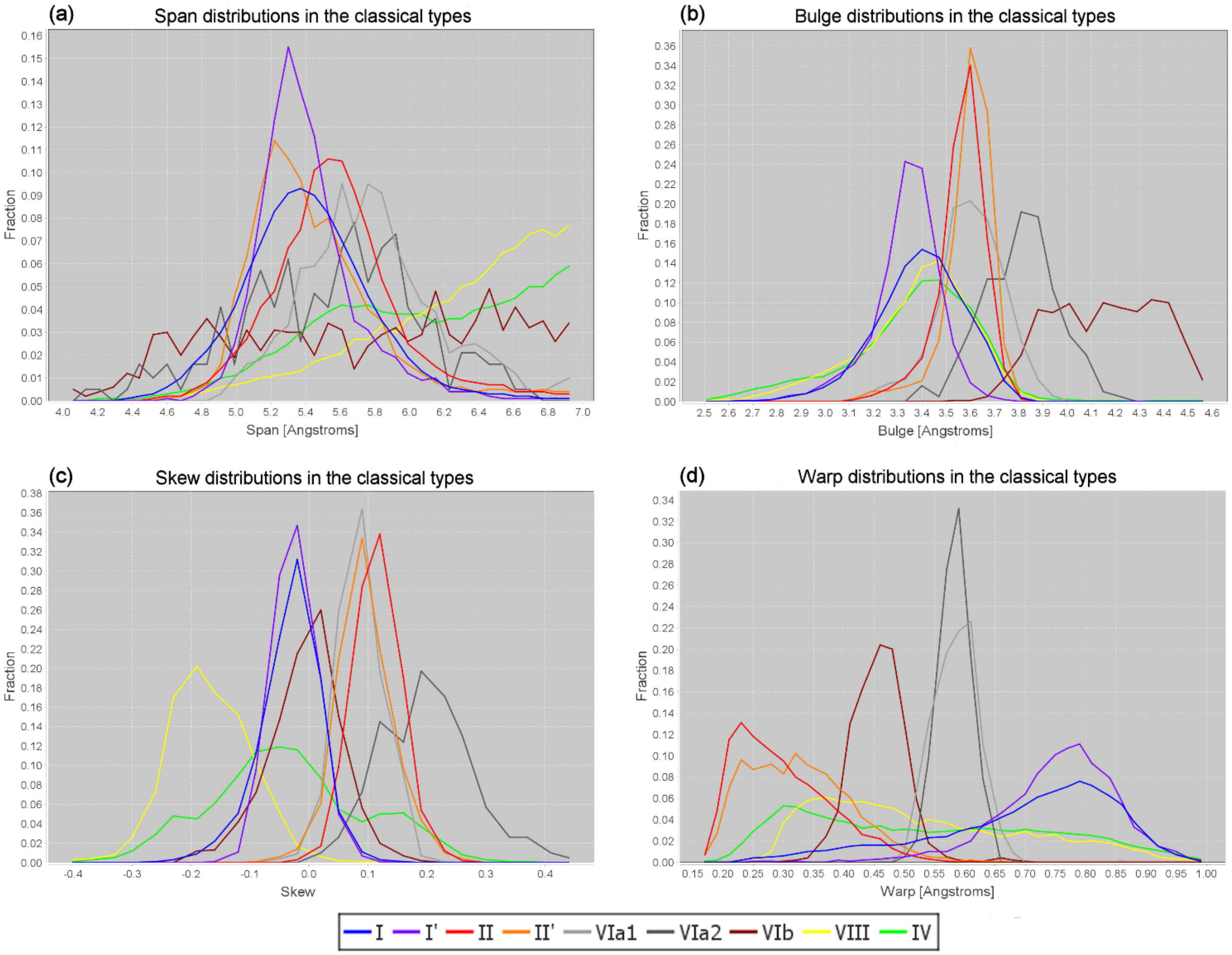


**Figure S1. Distributions of the geometric descriptors in the classical turn types.** Interpolated histograms of the distributions of the geometric descriptors[1,2] in the structurally classified classical turn types and the structurally unclassified type IV. The vertical axis measures turn fraction within each type. **(a)** Span measures $C_{\alpha 1} \rightarrow C_{\alpha 4}$ distance in the turn. The type I' span distribution peaks sharply near 5.3 Å, reflecting the common role of turns of this type as chain-reversers in 2:2 beta hairpins, and contrasting with the broader peak of the type I distribution, which is consistent with the wider range of type I roles. Types VIb, VIII and IV all show substantial abundance out to the 7.0 Å limit imposed by the beta-turn definition, reflecting the scarcity of the constraining 4>1 H-bond in these types. **(b)** Bulge measures the excursion the BB makes from a turn's span line (the line between $C_{\alpha 1}$ and $C_{\alpha 4}$) as it bends to form the turn. The bulge distributions for types VIa2 and VIb extend well to the right of all other types, and types II/II' show greater bulge than I/I'. All bulge distributions exhibit negative Pearson skewness[3], with more substantial tails on the left, indicating that the upper limit of bulge is "harder" than its lower limit: high-bulge/low-span (tighter) turns are more likely to be associated with conformational strain and steric clashes within the turn than low bulge/high span turns. **(c)** Skew measures a turn's N-terminal (negative) or C-terminal (positive) "lean" (asymmetry in the turn plane). The skew distributions for types {I, I', VIb} and {II, II', VIa1} can be formed into two loose groups that peak near skew 0.0 and 0.1 respectively, while types VIII and VIa2 peak to the left and right of these groups, and the structurally unclassified type IV covers the range of all structurally classified types. **(d)** The distributions of warp, which measures a turn's departure from flatness, form five groups from left to right, consisting of types {II, II'}, {IV, VIII}, VIb alone, {VIa1, VIa2}, and {I, I'}. Groups {II, II'} and {IV, VIII} exhibit positive Pearson skewness, with long tails to the right, while {I, I'} shows negative skewness. The similarity between the type IV and VIII distributions in the bulge and warp plots is notable.

### Table S1. Selecting turns with geometric descriptors near their limits

| Type of extreme | Selection criteria | Notes |
|---|---|---|
| Low span | ***Span Max***: 4 | |
| High span | ***Span Min***: 6.99 | High span (by definition, 7.0 Å is the limit for a beta turn). |
| Low N-terminal half-span | ***Span Max***: 1.4/ | Selects turns with a maximum N-terminal half-span of 1.4 Å. |
| High C-terminal half-span | ***Span Min***: /4.4 | Selects turns with a minimum C-terminal half-span of 4.4 Å. |
| Low bulge | ***Bulge Max***: 2.25 | |
| High bulge | ***Bulge Min***: 4.6 | |
| Negative skew | ***Skew Max:*** -.45 | |
| Positive skew | ***Skew Min:*** .4 | |
| Low warp | ***Warp Max***: .17 | |
| High warp | ***Warp Min***: 1.02 | |
| High warp in type II | ***β-turn type/cluster***: II<br>**Warp Min:** .59 | Although type II is strongly biased toward low warp (to see this, click ***Distbn*** in the *Warp* column), some turns of this type exhibit a substantial departure from flatness. |
| Low warp in type I | ***β-turn type/cluster***: I<br>**Warp Max:** .32 | Although type I is biased towards high warp, turns of this type display a very wide warp range; the 719 turns selected here are flatter than 39% of all type II turns. |

**Notes**: to view a set of structures, enter its selection criteria, click ***Load Selected Structures***, then click ***Browse Structures*** to step through the set by the interval specified in the ***Stepsize*** box (negative values specify back-steps). To change the radius for the display of structure external to the turn/tails, enter a new value in the ***Display radius*** box and click ***Browse Structures***. Click ***Clear All Criteria*** before entering a new example.

### 2 | Tuning turn geometry for compatibility with SC motifs

Since geometric descriptors enable Euclidean-space structural discrimination within the classical turn types and BB clusters, they yield major improvements in the precision of measurements of sequence preferences in turns. ExploreTurns provides plots of the fractional overrepresentation of sequence motifs vs. descriptor value for all descriptors for all single-AA and pair sequence motifs with sufficient abundance within turns and their two-residue BB neighborhoods. These plots, which are displayed by clicking ***Distbn*** in a descriptor's column after entering a sequence motif in the ***Sequence motif*** box and a type or BB cluster in the ***β-turn type/cluster*** box, identify the descriptor regimes that are most compatible with a sequence motif, and therefore most likely to be compatible with any associated SC interaction (SC motif). The plots support the tuning of turn BB geometry for compatibility with a particular SC motif, via the selection of turns with descriptor values that maximize the fractional overrepresentation of the associated sequence motif (see *Sequence motif selection…/Examples/Ex. 4* in the ExploreTurns feature summary).

Examples of motif overrepresentation plots are displayed in Figures S2 (for single-AA motifs) and S3 (for pair motifs). In many cases, the peaks and trends shown in these plots can be rationalized as compatibilities between the geometries specified by the descriptor values and the SC motifs associated with the sequence motifs (see the captions). Figure S3 displays example structures for three pair SC motifs, and Table S2 gives examples of warp-tuned selection criteria for several sequence motifs.

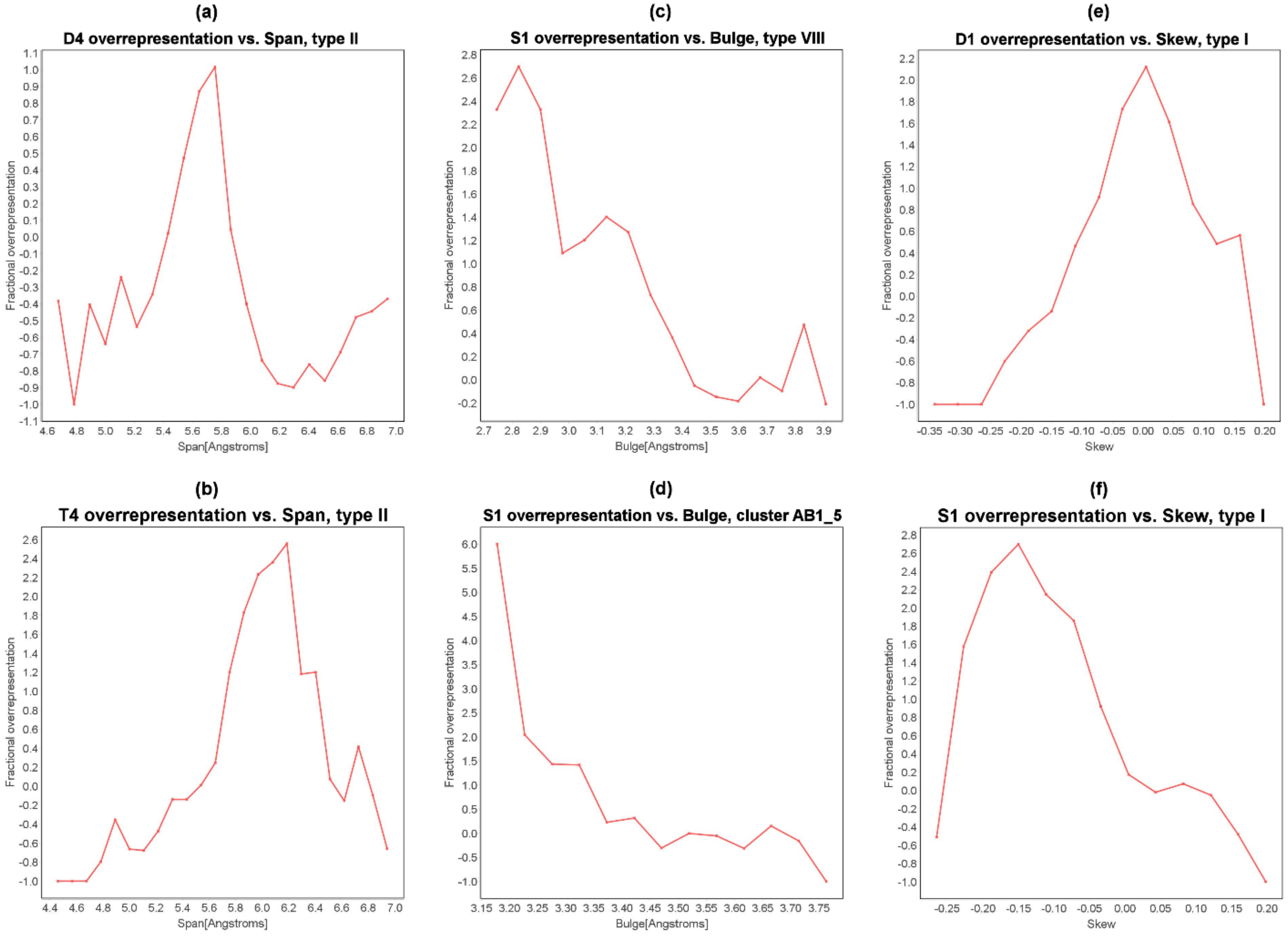


**Figure S2. Examples of sequence motif overrepresentation vs. descriptor value plots for single-AA motifs and the span, bulge and skew descriptors. (a, b)** Plots of fractional overrepresentation vs. span for Asp and Thr at turn position 4 (D4 and T4) in type II turns show prominent maxima that likely correspond to the optimal spans for SC/BB H-bonding with the main-chain NH group of the first turn residue (MCN1, for D4) or the main-chain carbonyl group of that residue (MCO1, for T4). **(c)** The overrepresentation of Ser at position 1 (S1) drops as bulge rises in type VIII, likely reflecting the lengthening and weakening of the H-bond between Ser's SC and MCN3 (adjacent to the turn center) in this ST turn/motif[4]. **(d)** S1's overrepresentation falls as bulge increases in the type VIII-associated BB cluster AB1_5, demonstrating that motifs are tunable within BB clusters as well as classical types. **(e, f)** While Asp at position 1 (D1) in type I turns shows peak overrepresentation near zero skew, Ser at position 1 (S1) shows a negative-skew bias; this contrast may reflect the suitability of a short N-terminal half-span (which promotes negative skew) for optimal H-bonding between Ser's shorter SC and the BB NH group at the turn's C-terminus (MCN4).

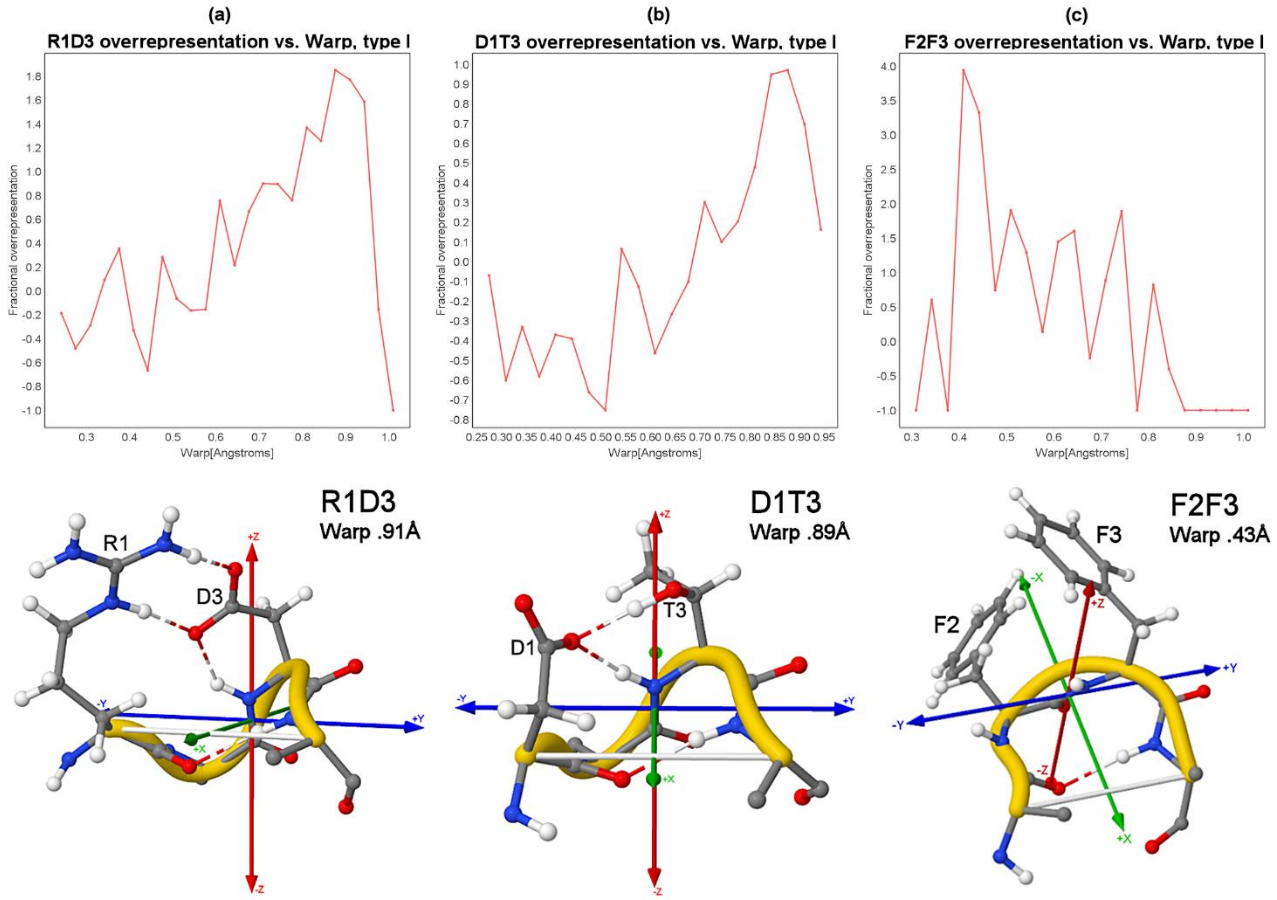


**Figure S3. Examples of sequence motif overrepresentation vs. descriptor value plots for pair motifs and the warp descriptor, with example structures for the associated SC motifs. (a, b) The** Arg1+Asp3 (R1D3) motif, commonly associated with salt bridges, and the Asp1+Thr3 (D1T3) motif, which often forms SC/SC H-bonds, show strong preferences for high warp, which likely optimizes SC positioning by raising the SC at position 3 (4A56_A_55 and 3AL2_A_1301). **(c)** Phe2+Phe3 (F2F3) favors lower warp, which brings the SCs closer together for perpendicular pi-stacking (3PYC_A_29).

### Table S2. Selecting sequence motifs tuned by warp

| Motif type | Selection criteria | Notes |
|---|---|---|
| Asx motif | **β-turn *type/cluster***: I<br>***Sequence motif***: D1<br>***BB H-bonds...***: 4>1<br>***Warp Max***: .85<br>***Warp Min***: .8 | The most common SC H-bonding motif in type I turns. Here the turn's BB geometry has been tuned by warp to maximize motif overrepresentation (and likely compatibility with SC/BB H-bonds). |
| ST motif | **β-turn *type/cluster***: I<br>***Sequence motif***: S1<br>***BB H-bonds...***: 4>1<br>***Warp Min***: 1.0 | Tuned by warp to maximize motif overrepresentation and SC/BB H-bonding. |
| SC/SC H-bonding | **β-turn *type/cluster***: I<br>***Sequence motif***: D1T3<br>***Warp Max***: .87<br>***Warp Min***: .85 | Tuned to high warp for compatibility with SC/SC H-bonding. |
| Salt bridge | **β-turn *type/cluster***: I<br>***Sequence motif***: D1R3<br>***Warp max***: 1.0<br>***Warp min***: .95 | Tuned to high warp for compatibility with salt bridging. |
| Pi-stacking interaction | **β-turn *type/cluster***: I<br>***Sequence motif***: F2F3<br>***Warp max***: .45<br>***Warp min***: .4 | Tuned to low warp, which can bring the SC rings closer together for perpendicular pi-stacking. |

**Notes**: see notes for Table S1. After entering selection criteria for a motif, click ***Distbn*** to display its plot of overrepresentation vs. warp, or click ***Map Motif*** to map its structure and H-bonding across all BB geometries[5].

### 3 | Ramachandran-space descriptor heatmaps

ExploreTurns provides heatmaps of the distributions of the turn descriptors across the BB dihedral-angle (Ramachandran) spaces of the two central residues of the turns in each type, BB cluster, or the global turn set. Descriptor heatmaps characterize the bond rotations associated with the variations of the geometric descriptors, and reveal the conformations that yield the lowest energies for the 4>1 beta-turn H-bond. To open a heatmap, click ***Heatmp*** in a descriptor's column after entering a type or cluster in the ***β-turn type/cluster*** box (if the box is left blank, then the map for the set of all turns is displayed). Figure S4 displays examples of descriptor heatmaps, with interpretations in the caption.

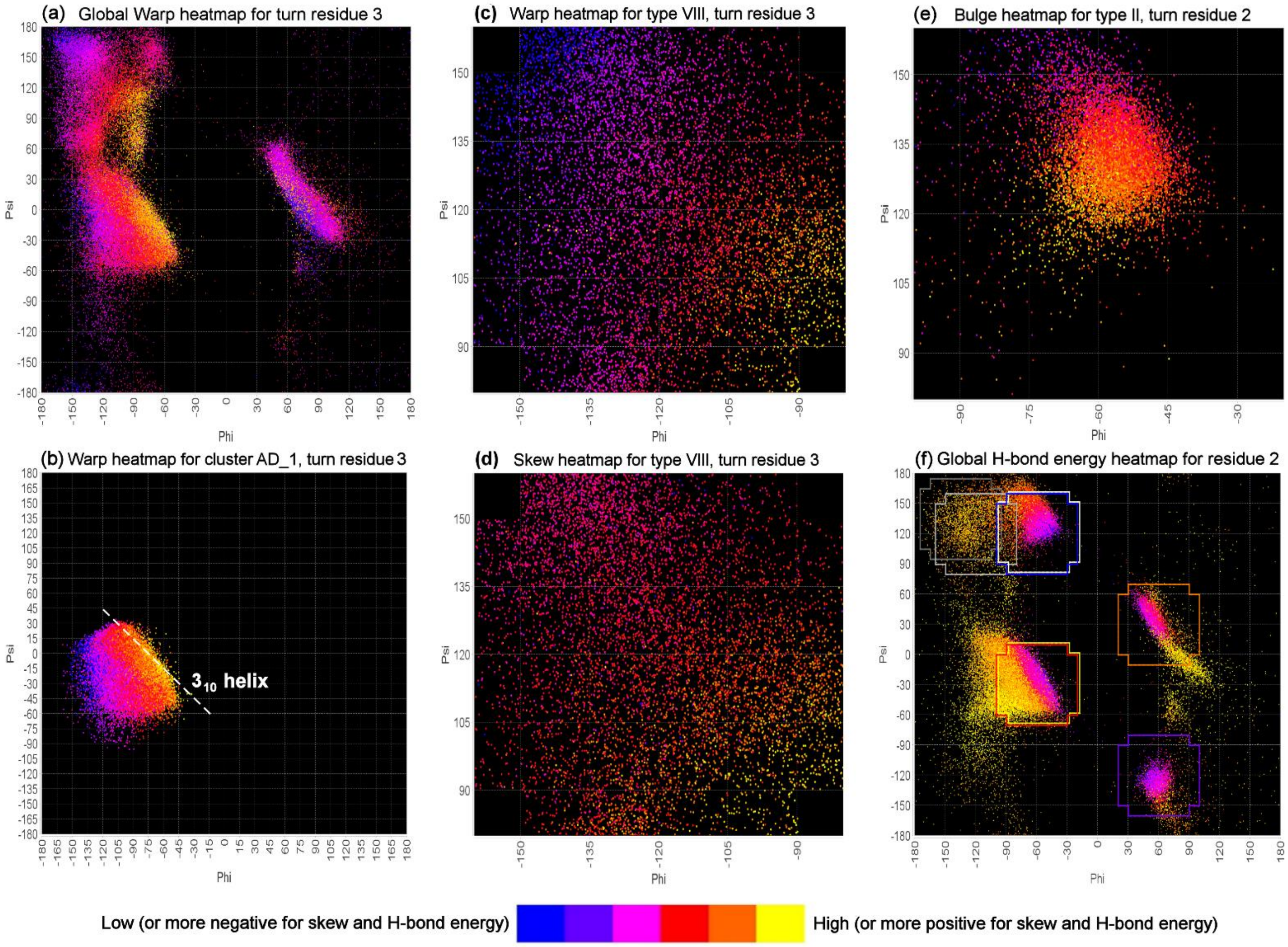


**Figure S4. Ramachandran-space heatmaps characterize the bond rotations associated with descriptor variation.** Descriptor values are colored on relative scales within each plot according to the legend; actual values in a Ramachandran-space region can be obtained from ExploreTurns. **(a)** The most characteristic feature of high-warp turns is the orientation of the second peptide bond approximately perpendicular to the turn plane, and the global warp distribution at residue 3 shows a rising gradient of warp over a wide range of $\psi_3$ as $\varphi_3$ rises towards -30° (becoming increasingly acute), accompanying the rotation of the peptide bond towards a perpendicular orientation and supporting the sharper BB reversals required at $C_{\alpha 3}$ to direct the BB back down towards the turn plane after the large z-dimensional excursions associated with high warp. **(b)** The warp heatmap for residue 3 in the type I-associated BB cluster AD_1, which contains about half of all beta turns, demonstrates the relationship between warp and helicity: warp increases as the conformation approaches the line along which $(\varphi, \psi)$ sums to -75°, ideal for a $3_{10}$ helix; the $3_{10}$ helix can be viewed as an extension of a (high-warp) beta turn[6]. **(c, d)** The warp and skew heatmaps for residue 3 in type VIII turns show broad, highly correlated gradients of falling warp and increasingly negative skew (from lower-right to upper-left in the plots) as the residue opens into a more extended conformation, both flattening the turn and increasing its C-terminal half-span (and consequently its N-terminal "lean", or negative skew). **(e)** At the second residue of type II turns, rising $\psi_2$ reduces bulge by rotating the $C_{\alpha 2} \rightarrow N_2$ bond "outwards and upwards" (towards -y, +z), effectively reducing the x components of the three main-chain-path bonds in the plane of the turn's first peptide bond. **(f)** The heatmap for the electrostatic energy of the 4>1 H-bond at residue 2 in the global turn set shows the energy minima at the cores of classical types I, I', II/VIa1, and II'. The frames outline the allowed conformations in each classical type, with color-coding: **Red**: I, **Orange**: I', **Blue**: II, **Purple**: II', **White**: VIa1, **Light Grey**: VIa2, **Dark Grey**: VIb, **Yellow**: VIII (frames that overlap are shifted slightly).

### 4 | Selecting supersecondary structures with context vectors

Table S3 lists selection criteria for five examples of supersecondary structures in which beta turns link helices/strands into particular relative orientations, and Figure S5 displays examples; a helix kink[7] is also included. The ExploreTurns "OR" mode for ***DSSP symbols*** entry, specified by vertical bars surrounding the turn instead of the angle brackets of the default "AND" mode, is used for all examples here except the kink, in order to increase the size of the selected sets by allowing flexibility in the start positions of the secondary structures adjacent to the turn. For example, ***HH|H**H|HH*** specifies that helices must be present before and after the turn, but provides flexibility in the helix start positions by requiring DSSP type 'H' either at the turn's terminal residues or within two residues of the turn (see *Secondary structure selection in the ExploreTurns frame* in the tool's feature summary, or *DSSP symbols* in the user guide).

Approximate orientations between secondary structure elements are selected by specifying the directions of the N- and C-tails using the ***Context vectors*** and ***Vector tolerances*** criteria (see *β-turn tail orientation using "context vectors"* in the feature summary). When an alpha helix begins within two residues of the turn, tail direction is computed as the direction of the helix axis; if the tail contains any other conformation (including strand), its orientation is determined directly from its alpha-carbon positions. The accuracy of context vectors is subject to the irregular nature of BB shape; the vectors are most accurate when the tails contain helices or low-curvature strands.

#### Table S3. Selection criteria for examples of supersecondary structures

<table>
<tr><th>Supersecondary structure</th><th>Selection criteria</th><th>Notes</th></tr>
<tr><td>helix↔helix junction<br>-y → +x</td><td>DSSP symbols: **HH|H**H|HH**<br>Context vectors: -90,0<>0,0<br>Vector tolerances: 30<>30</td><td>Alpha helix junction; N-tail towards -y, C-tail towards +x.</td></tr>
<tr><td>helix↔helix junction<br>-y → +z</td><td>DSSP symbols: **HH|H**H|HH**<br>Context vectors: -90,0<>0,90<br>Vector tolerances: 30<>30</td><td>Alpha helix junction; N-tail towards -y, C-tail towards +z. Contacts heme groups in cytochrome C's (e.g. 5B6Q_A_35, 1FS7_A_397).</td></tr>
<tr><td>helix↔helix junction<br>-z → +z</td><td>DSSP symbols: **HH|H**H|HH**<br>Context vectors: 0,-90<>0,90<br>Vector tolerances: 45<>45</td><td>Alpha helix junction; N-tail towards -z, C-tail towards +z. Binds the IHP cofactor in the RNA-editing enzyme ADAR2 (1ZY7_A_527).</td></tr>
<tr><td>strand↔helix junction<br>-y → +z</td><td>DSSP symbols: **EE|E**H|HH**<br>Context vectors: -90,0<>0,90<br>Vector tolerances: 35<>35</td><td>Strand-helix junction, N-tail towards -y, C-tail towards +z. Binds Mg/dTTP at the active site of DNA polymerase (4TQR_A_8).</td></tr>
<tr><td>strand↔strand junction<br>(+x, -y) → (+x, +y)</td><td>DSSP symbols: **EE|E**E|EE**<br>Context vectors: -45,0<>45,0<br>Vector tolerances: 30<>30</td><td>Strand-strand junction, N-tail towards (+x, -y), C-tail towards (+x, +y). Binds FMN in flavoredoxin (2D5M_A_33).</td></tr>
<tr><td>Helix kink[7]</td><td>DSSP symbols: ****<H**H>****<br>Context vectors: -90,20<>90,30<br>Vector tolerances: 20<>20</td><td>Helix "kink" formed by a beta turn.</td></tr>
</table>

**Notes:** (see notes for Table S1).

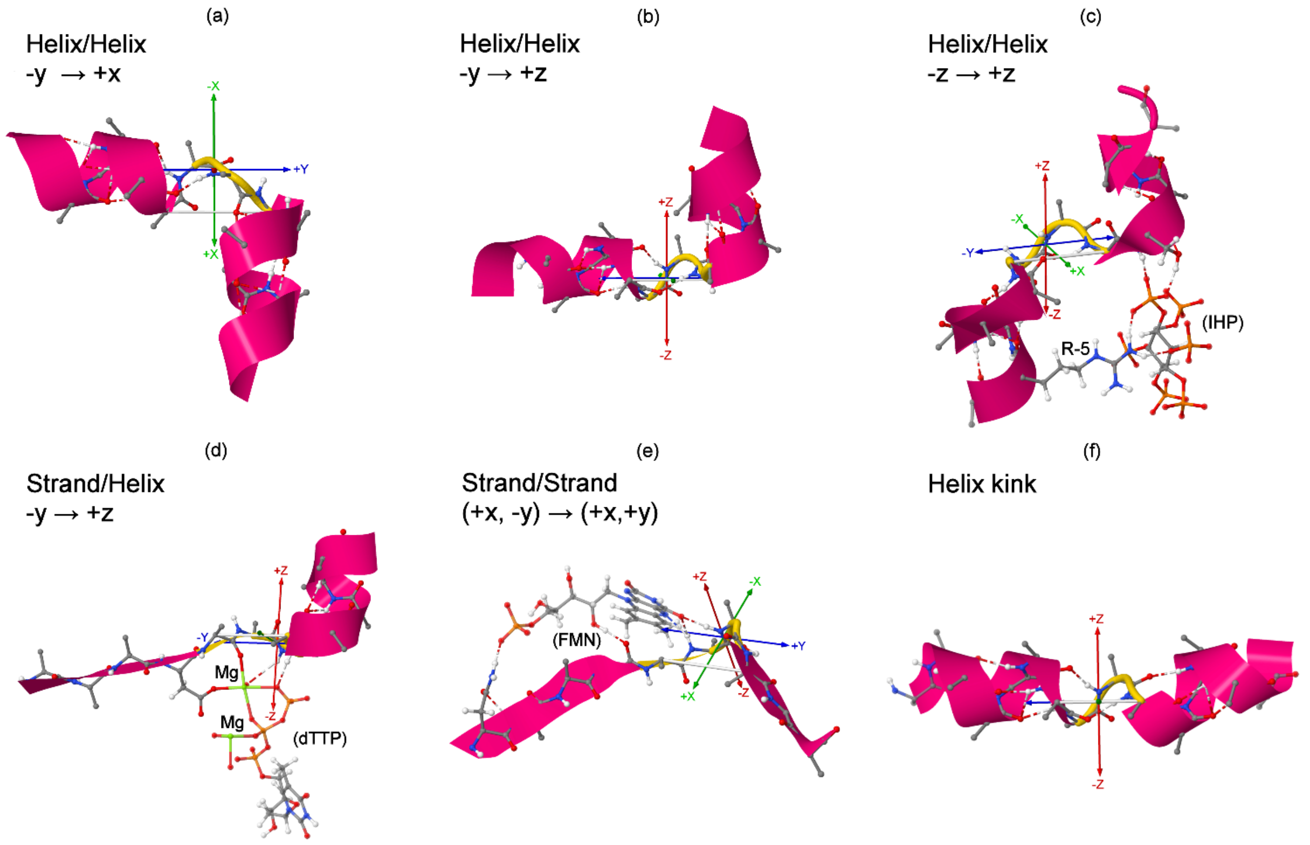


**Figure S5. Examples of supersecondary structures linked by beta turns**. To view the individual structures shown here, load the entire ExploreTurns database by clicking ***Load Selected Structures*** without any criteria, then enter the structure's address into the ***PDB address*** box and click ***Browse Structures***. **(a)** Helix-helix junction, -y → +x (4PP8_C_144). **(b)** Helix-helix junction, -y → +z (5B6Q_A_35). **(c)** Helix-helix junction, -z → +z, binding the cofactor IHP in the catalytic domain of the human RNA editing enzyme ADAR2 (1ZY7_A_527). **(d)** Strand-helix junction, -y → +z, binding Mg and dTTP at the active site of DNA polymerase (4TQR_A_8). **(e)** Strand-strand junction, (+x, -y) → (+x, +y), binding FMN in flavoredoxin (2D5M_A_33). **(f)** Helix kink[7] (5AQM_B_201).

### 5 | Classifying and exploring non-BBL H-bonded loop motifs and variants

Table S4 provides compound-turn labels and notes that support the exploration of non-BBL H-bonded loop motifs and variants. Figure 3 in the main text compares the structures of 12 of these motifs.

### Table S4. Selection criteria for non-BBL H-bonded loops and variants

| Motifs and variants, classified with the Greek CT nomenclature | Romanized CT labels | Additional notes |
|---|---|---|
| Alpha-beta loop[8]<br>α-β1 turn | **a-b1** All variants in most common frame<br>**p'*-a1-b1** *p'** extends the CT label to exclude a π'-turn, ruling out 6C1 BBLs | The name apparently refers to secondary structures linked by the motif, not H-bonded turn types. |
| Crown bridge loop[9]<br>π-β1 turn<br><u>Variants:</u><br>π-α1-β1 Triple nest<br><br>π-α1-β1-β'3 Crown pyramid[10] in Mg phosphatases | **@p-b1_*** *'@'* specifies permissive baseline mode, excluding 6C1, Schell.<br><br>**@p-a1-b1_*** *'@'* excludes 6C1 BBLs and the variant below<br>**p-a1-b1-b'3_3** *_3* selects the 3rd most common position frame, in which the β-turn at the center of the ExploreTurns window forms the pyramid's "base" | Additional H-bonds in this motif first form a triple nest[11], then an active site motif found in Mg phosphatases in all three kingdoms[10]. |
| Schellman loop[12,13]<br>π-β2 turn<br><br><u>Six-residue variants:</u><br>π-β2$\varphi^{3,4}\varphi_5$ SchellmanLH<br>π-β1-β2 Schellman+β[8]<br>π-β1-β2 Schellman310<br>π-α1-β2 Schellman+α[8]<br><u>Other forms:</u><br>ρ*-ρ'*-α1-π2-β3 α+Schellman[8]<br>ρ-π1-β2 Schellman+ρ<br>ρ-π1-α1-β2 Schellman+α+ρ<br>σ-π2-β3 Schellman extended<br>τ*-τ'*-π1-β2-π4-β5 Double Schell | **p-b2** All variants in most common frame<br>**p-p'*-b2_*** All variants except 6C1 BBL in all position frames (selected with '*')<br><br><br>**p-b2-\F3-\F4-\f5_*** LH turn/RH bend<br>**@p-p'*-b1-b2** Adds β-turn in loop<br>**p-b1-b2-\h1-\H2-\H3-\H4-\h5_*** $3_{10}$ hel<br>**P-p'*-a1-b2** Adds α-turn in loop<br><br>**R*-r'*-a1-p2-b3-\h1_*** overlaps α-turn<br>**R-p1-b2** Adds ρ-turn; forms nest[11]<br>**R-p1-a1-b2** Adds α and ρ; triple nest<br>**s-p2-b3_*** Embeds loop in a σ-turn<br>**t*-t'*-p1-b2-p4-b5_*** Double Schellman | Commonly forms dual helix C-caps with its defining pair of BB H-bonds. The common RH form combines an RH (type I) turn with an LH bend at its end; the LH form is a BB mirror image. Many Schellman loops are absent from the database due to helix overlap with the turn, but 2318 examples are available.<br>**p-b2-\H1-\H2 _*** selects C-caps only; **p-b2-\H5-\H6_*** N-caps only. The Schellman310 and the last 4 loops were detected with ExploreTurns. |
| Wide Schellman loop[14]<br>ρ-α2 turn | **r-a2_*** Adds a residue inside the β-turn, forming an α-turn. | Seven-residue loop. For dual C-caps only, enter **r-a2-\H1-\H2_*** |
| Short Schellman loop<br>α-γ2 turn<br><u>Dihedral variants</u><br>α-γ2$\varphi_3\varphi^4$<br><br>α-γ2$\varphi^3\varphi_4$ | **a-g2_***<br><br><br>**a-g2-\f3-\F4_*** Analogue of RH Schell<br><br>**a-g2-\F3-\f4_*** Analogue of LH Schell | Five-residue motif detected with ExploreTurns. Drops a residue from the Schellman loop. Select helix N-caps with **a-g2-\H4-\H5 _*** or C-caps with **a-g2-\H1-\H2 _*** |
| Pi-alpha loop<br>π-α1 turn<br>π-π'*-α1-β*1-γ*2-β*2<br>(Exclusions) | **p-a1**<br><br>**p-p'*-a1-b*1-g*2-b*2-\h2_*** *\h2* rules out loops within helices | Six-residue motif detected with ExploreTurns; adds a residue to the alpha-beta loop. {*p'**, *b*1*, *b*2*, g*2} rule out 6C1 BBLs, crown bridge loops, Schell + short Schell loops. |
| Beta-gamma loop<br>β-γ1 turn | **b-g1** | Detected with ExploreTurns; high frequency of PDB outliers. |
| Beta bracket<br>ρ*-ρ'*-β1-α'3-β4 | **r*-r'*-b1-a'3-b4**<br>**r*-r'*-b1-a'3-b4-\h5** excludes $3_{10}$ helix | β-turn pair constrained by an α'-turn, detected with ExploreTurns. |
| Long beta bracket<br>σ*-σ'*-β1-p'3-β4 | **s*-s'*-b1-p'3-b4_2**<br>**u*-u'*-b3-p'5-b6-\E1** Selects only "loopback" sheet exits in α/β hydrolases | A π'-turn replaces the bracket's α'-turn. Can form a conserved sheet-exit loop in α/β hydrolases. |

**Notes**: To select a loop's structures, enter its CT label into the ***BB H-bonds/Loop motif/Compound turn*** box and click ***Load Selected Structures***. Motifs may also be selected by entering the names given in the first column into the ***BB H-bonds...*** box (with Greek letters translated to English words), or the loops can be explored using the links in the loop motif browser, which is annotated with structural schematics. The H-bonds displayed by JSmol can differ from those selected in ExploreTurns due to differences between the JSmol and ExploreTurns H-bond definitions.

### 6 | Classifying and exploring beta-bulge loops

Tables S5 and S6 provide compound-turn labels and notes that support the exploration of the most abundant beta-bulge loops and variants with C-terminal bulges (Table S5) and N- or N/C-terminal bulges (Table S6). Table S7 provides labels and notes for a set of longer beta-bulge loops and variants.

#### Table S5. Beta-bulge loops with C-terminal bulges

| BBL types/Greek CT labels | Romanized CT labels (top entries select both dihedral variants, but exclude all BB H-bond variants) | Additional notes |
|---|---|---|
| **5C1** (type 1)<br>α'-β1 turn<br><u>Variants</u><br>**5C1+**: α'-β1$\varphi^4$ ($\varphi_4 > 0$)<br>**5C1-**: α'-β1$\varphi_4$ ($\varphi_4 < 0$)<br>**5C1 extended**: τ-α'3-β3 | **A'-b1**<br><br><br>**A'-b1-\F4** Very common; "reclined chair".<br>**a'-b1-\f4** Bulge projects towards -z.<br>**t-a'3-b3-\e1-\e2-\e8-\e9** (β-ladders excluded) | Capitalization of the BBL's framing α-turn (in **A'-b1**) excludes additional BB H-bonds in the loop, selecting its strict "baseline" form. |
| **6C1** (type 2)<br>π'-α1 turn<br><u>Variants</u><br>**6C1+**: π'-α1$\varphi^5$<br>**6C1-**: π'-α1$\varphi_5$<br>**6C1 extended**: υ-π'3-α3 | **P'-a1**<br><br><br>**P'-a1-\F5** "Upright chair".<br>**p'-a1-\f5** Commonly forms a "hook".<br>**u-p'3-a3-\e1-\e2-\e9-\e10** (β-ladders excluded) | Adds a residue to the 5C1 BBL's internal β-turn, forming an H-bonded α-turn. |
| **6C2**<br>π'-β1 turn<br><u>Variants</u><br>**6C2+**: π'-β1$\varphi^4$ ($\varphi_4 > 0$)<br>**6C2-**: π'-β1$\varphi_4$ ($\varphi_4 < 0$)<br>**6C2 extended**: υ-π'3-α*3-β3 | **P'-b1**<br><br><br>**P'-b1-\F4** Bulge projects toward +z.<br>**P'-b1-\f4** Bulge projects towards -z.<br>**u-p'3 -a*3-b3-\e1-\e2-\e9-\e10** (β-ladders excluded, *a*3* excludes 6C1 extended) | Adds a residue to the 5C1 BBL's bulge. |
| **7C3**<br>ρ'-β1 turn<br><u>Variants</u><br>**7C3+**: ρ'-β1$\varphi^4$ ($\varphi_4 > 0$)<br>**7C3-**: ρ'-β1$\varphi_4$ ($\varphi_4 < 0$) | **R'-b1**<br><br><br>**R'-b1-\F4** Bulge projects towards +z.<br>**R'-b1-\f4** Bulge projects towards -z. | Adds a residue to the 6C2 BBL's bulge. |

**Notes**: See notes for Table S4. Loops and variants can be selected with either the bolded BBL shorthands given in column 1 or the romanized CT labels given in column 2.

### Table S6. Beta-bulge loops with N- or N/C-terminal bulges

| BBL types/Greek CT labels | Romanized CT labels | Additional notes |
|---|---|---|
| **5N1**<br>α'-β2 turn<br><u>Variants</u><br>α'-β2-β'2<br>α'-β2-β'*2 | **A'-b2**<br><br><br>**a'-b2-b'2** Adds a reverse β-turn.<br>**a'-b2-b'*2** Excludes reverse β-turn. | In its beta hairpin context, this motif can be formed by the insertion of a residue just before the chain-reversing turn of a 2:2 hairpin. The **a'-b2-b'2** variant, which adds a reverse β-turn to the loop's defining normal β-turn, represents about half of all instances of the motif. |
| **6N2**<br>π'-β3 turn | **P'-b3** | Adds a residue to the 5N1 BBL's bulge. |
| **7N3**<br>ρ'-β*1-β4 turn | **r'-b*1-b4** | Adds a residue to the 6N2 BBL's bulge.<br>*b*1* excludes the H-bonded β-turn which would yield the 7NC3 BBL below. |
| **7NC3**<br>ρ'-β1-β4 turn | **R'-b1-b4** | Since this loop contains split pairs of BB H-bonds at both termini, it is classified as both N- and C-type. |

**Note**: See notes for Table S5.

### Table S7. Additional beta-bulge loops and variants

| BBL types and variants, with Greek CT labels | Romanized CT labels | Additional notes |
|---|---|---|
| **6N2C3**<br>π'-γ1-β3 turn<br>Variants<br>π'-γ1-β3$\varphi_6$ (C-tail orients to -z)<br>π'-γ1-β3$\varphi^6$ (C-tail orients to +z) | **p'-g1-b3**<br><br><br>**p'-g1-b3-\f6**<br>**p'-g1-b3-\F6** | Examples of the $\varphi_6$ variant bind RNA (1WPU_A_131, Figure 8d in the main text) and separate DNA strands in the helicase reaction (3AAF_A_1033, Figure 8e). |
| **6N3C2**<br>π'-β1-γ4 turn | **p'-b1-g4** | |
| **7C2**<br>ρ'-α1 turn | **r'-a1** | Adds a residue to the 6C1 BBL's bulge. Four examples bind metals. |
| **7C4**<br>ρ'-γ1 turn<br>Variants<br>ρ'-ρ*-γ1-β*3 (Excludes ρ, β H-bonds)<br>ρ'-ρ*-γ1-β3 (Includes β H-bond)<br>ρ'-ρ-γ1-β*3 (Includes ρ H-bond) | **r'-g1**<br><br><br>**r'-r*-g1-b*3** Excludes ρ,β H-bonds<br>**r'-r*-g1-b3** Includes β3 H-bond<br>**r'-r-g1-b*3** Includes ρ1 H-bond | |
| **7N3C1C3**<br>ρ'-π1-β1-β4 turn | **r'-p1-b1-b4**<br><br>**r'-p1-b1-b4-\h2** Excludes $3_{10}$ hel.<br>**r'-p1-b1-b2-b4** Adds β-turn, $3_{10}$ helices occur in all examples<br><br>**r'-p1-b1-b4-N1** Asx SC motif | Addition of a π1 H-bond to 7NC3 yields overlapping one- and three-residue C-terminal bulges and draws the β-turn "lobes" together. |
| **8C3**<br>σ'-α1-β*5 turn (β*5 excludes 8N4C3) | **s'-a1-b*5** | Adds a residue to 7C2's bulge. One example coordinates Ca with 5 bonds. |
| **8N4**<br>σ'-β*1-β5 (β*1 excludes 8NC4)<br>Variants<br>σ'-β*1-β2-β5 (β-turn in bulge)<br>σ'-β*1-γ'3-β5 (H-bond bridge)<br>$3_{10}$ helix in bulge<br>Excludes $3_{10}$ helix in bulge | **s'-b*1-b5**<br><br><br>**S'-b*1-b2-b5**<br>**s'-b*1-g'3-b5**<br>**s'-b5-\H3**<br>**s'-b5-\h3** | |
| **8C4**<br>σ'-β1 turn<br><br>Variant<br>σ'-β1-β4 (β-turn in bulge, not 8NC4) | **s'-b1-b*5** (b*5 excludes 8NC4)<br><br>**S'-b1** Excludes all variants<br><br>**S'-b1-b4** | |
| **8NC4**<br>σ'-β1-β5 turn | **s'-b1-b5** | Split pairs of BB H-bonds at both termini classify this motif as N- and C-type, with 4-residue bulges. Binds DNA (5CIY_A_232, Figure 8i). |
| **8N3C4**<br>σ'-β1-α4 turn | **s'-b1-a4** | Classified as both N-type/3-residue bulge and C-type/4-residue bulge. |
| **8N4C3**<br>σ'-α1-β5 turn | **s'-a1-b5** | Classified as both N-type/4-residue bulge and C-type/3-residue bulge. |

**Notes:** See notes for Table S5.

### 7 | Mapping Asx N-cap sequence preference vs. cap geometry

At the helix N-terminus, unsatisfied BB NH groups are commonly "capped" by H-bonding with the SCs of Asp or Asn (Asx) or Ser or Thr (ST) in Asx/ST N-caps[4,15-17]. Beta turns are also common at the helix N-terminus[16], and the Asx AAs in helix-terminal turns show greatest overrepresentation when they occur at the third turn position and this position coincides with the NCap residue at the start of the helix. ExploreTurns' "context vectors" specify the approximate directions of the turn's N- and C-terminal BB "tails" in the turn frame, and when a helix begins within two residues of the turn, tail direction is measured by the orientation of the helix axis, so the tool can be used to select sets of structures in which the helix has particular orientations with respect to the turn. Since the tool also computes sequence motif statistics for each selected set, it can be used to map N-cap sequence preference vs. helix/turn geometry, by evaluating motif overrepresentation and abundance in sets of structures spanning the range of helix/turn orientations.

The context vector for each tail is specified in a (longitude, latitude) format, with the "equator" lying in the x-y plane (the "turn plane"[1]), the zero of longitude at the equator corresponding to the +x direction, positive longitudes occurring in the +y half-space, and positive latitudes in the +z half-space (see *β-turn tail orientation using "context vectors"* in the feature summary, and Section 1.5 of the user guide). In this analysis, sets of structures are selected by specifying C-tail context vectors at 10° intervals of longitude and latitude, and the ***Vector tolerances*** field restricts the selected structures to those lying within 5° of a specified context vector.

Figure S6 displays an example of an Asp3 helix N-cap in a beta turn, and Figure S7 plots the abundance and overrepresentation of the Asp3 N-cap sequence motif vs. helix/turn orientation. The plot shows that Asp3's peak overrepresentation occurs at zero longitude, where the helix axis is parallel to the x-z plane, and latitudes just above the turn plane; overrepresentation maintains very high values as the helix rotates upward to 50° above the plane. As the helix rotates into negative longitudes, towards -y and across the mouth of the turn, overrepresentation and abundance fall off past -10°, while at positive longitudes, where the helix rotates towards +y and away from the turn, the two measures remain high as the helix projects above the turn plane at mid-latitudes (~45°), until ~50° longitude, where overrepresentation drops as the helix rotates downwards, crossing below the plane, and Asp3's SC loses access to the helix-terminal NH groups.

Figure S8 maps abundance and overrepresentation for the Asn3 N-cap motif, and shows that peak overrepresentation occurs not when the helix orients at zero longitude, as seen for Asp3, but when it straddles the meridian at longitudes of +/-10°, and peak overrepresentation is more focused in latitude. Figures S7 and S8 can guide ExploreTurns investigations of the structure and H-bonding associated with Asx N-caps formed by beta turns.

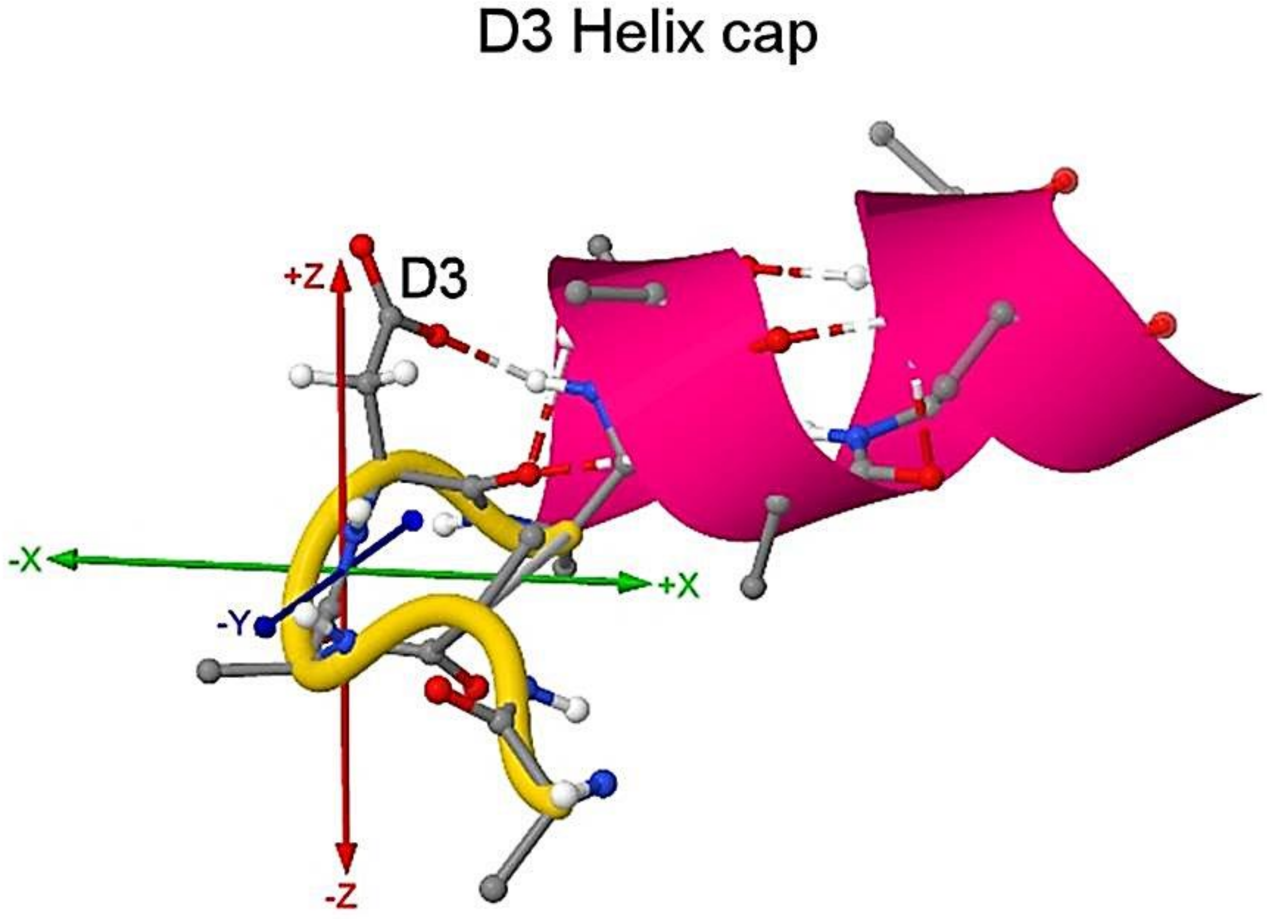


**Figure S6. Example of an Asp3 helix N-cap.** An Asp3 (D3) SC motif in a type VIII beta turn forms an Asx N-cap by H-bonding with an unsatisfied BB NH group at the helix terminus (5EJY_A_86).

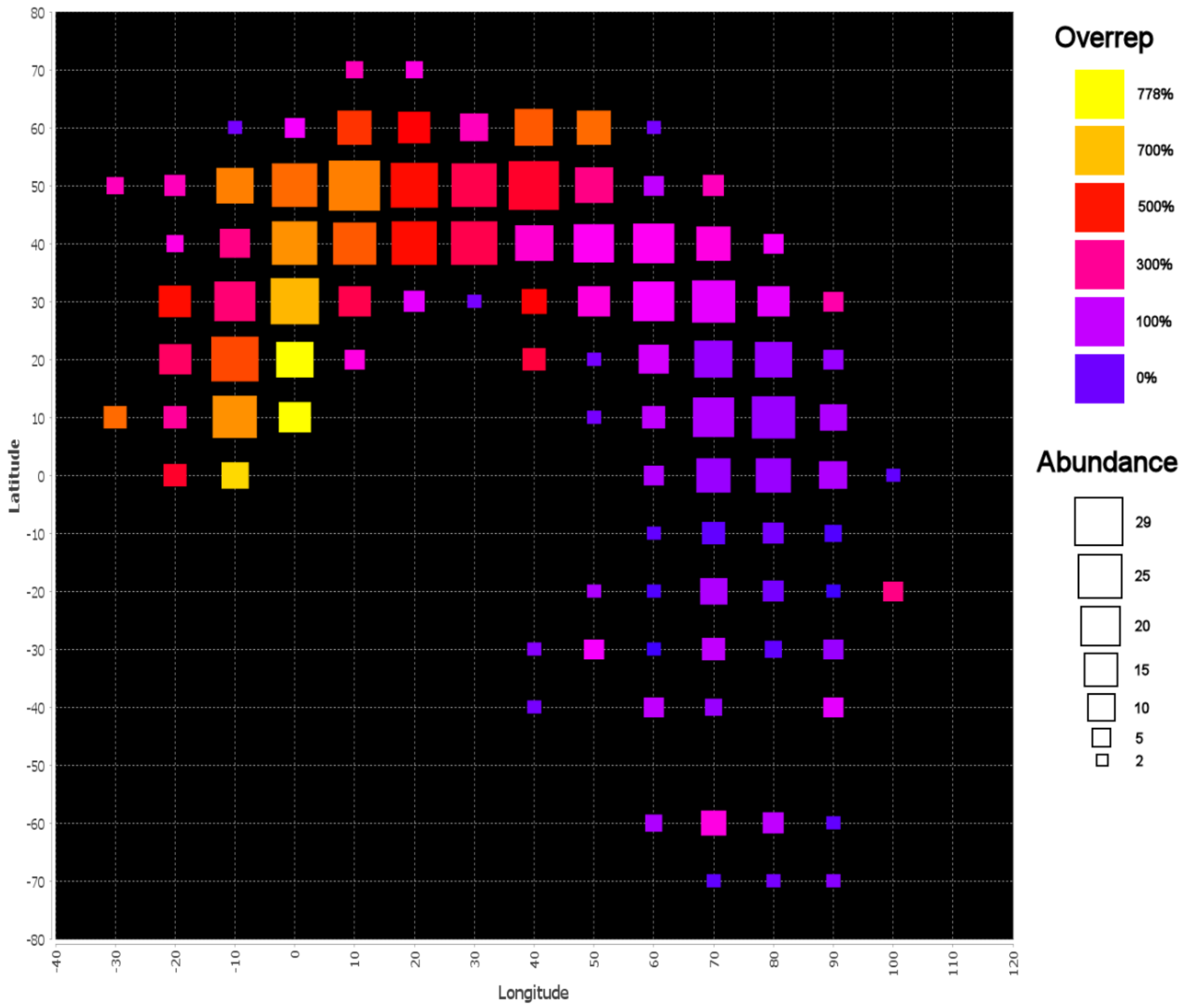


**Figure S7. Mapping Asp3 N-cap sequence preference vs. cap geometry.** Fractional overrepresentation and abundance are mapped vs. helix/turn orientation for the sequence motif that specifies Asp at turn position 3 at the helix N-terminus. Beta turns in which the third turn residue lies at the NCap position in an $\alpha$-helix are selected using the ***DSSP symbols*** entry ****<***H>****, and the Asp3 motif is specified with the ***Sequence motif*** entry *D3*. The orientations of the helix with respect to the turn are sampled at 10° increments by varying the direction of the C-tail (here the helix axis), using ***Context vectors*** entries of the form *,*<>longitude,latitude. The ***Vector tolerances*** entry *<>5 is applied to select structures with helix axes that lie within 5° of each context vector. Squares are plotted for all orientations that yield at least 10 structures, with at least one structure containing Asp3. Asp3's fractional overrepresentation is depicted as a heatmap colored according to the upper legend, while the area of each square is set proportional to the occurrence of Asp3 in each orientation according to the lower legend. Not all orientations with high Asp3 overrepresentation show frequent SC capping of helix BB groups, since Asp's hydrophilicity is also favorable at this commonly exposed position, and SC/SC H-bonding as well as non-capping SC/BB H-bonding in the structures also likely contribute to the preference for the motif.

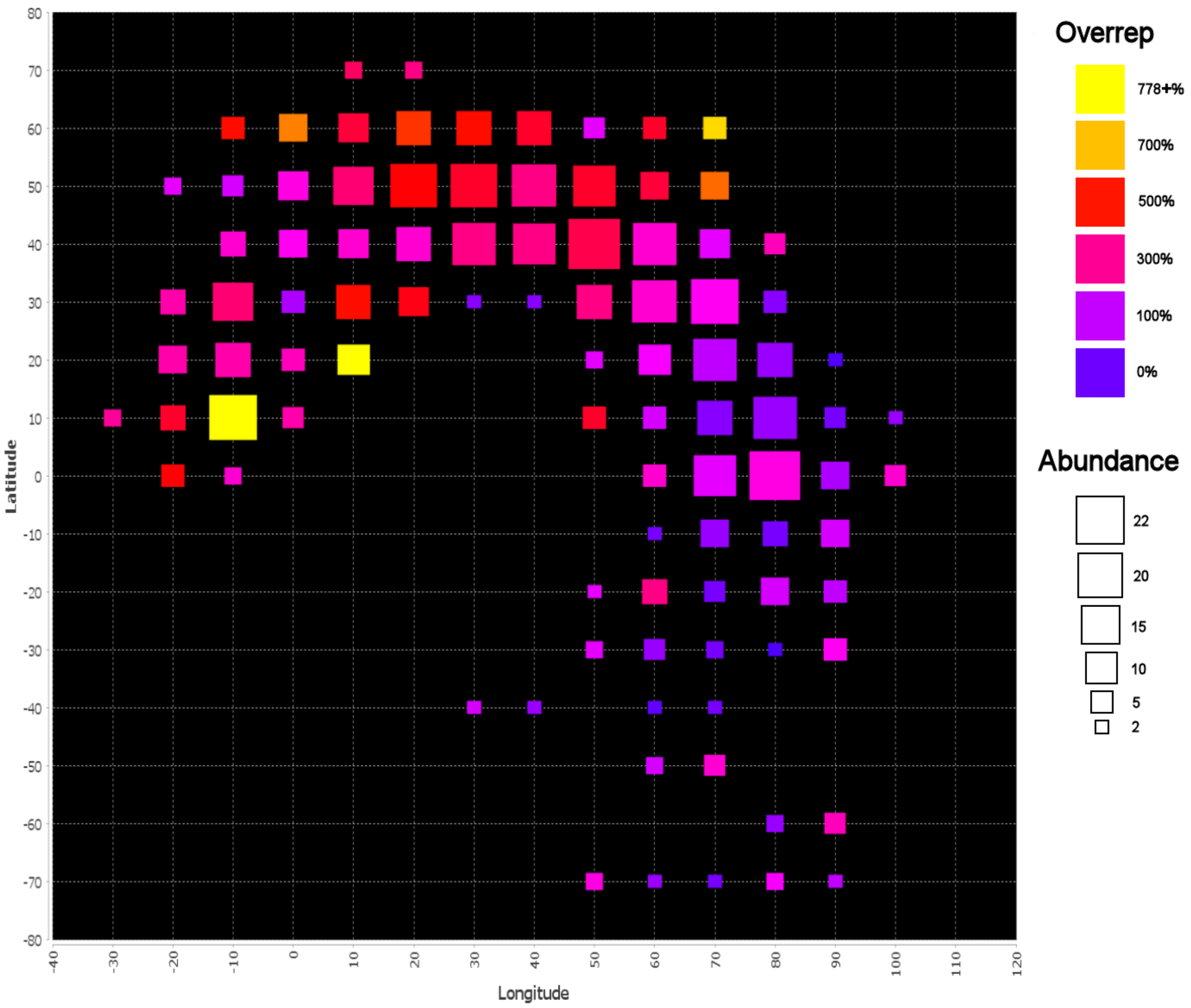


**Figure S8. Mapping Asn3 N-cap sequence preference vs. cap geometry.** See the Figure S7 caption for the methodology used to select the data in ExploreTurns (N3 is substituted for D3). In contrast to D3, N3 shows its highest overrepresentation not when the helix axis orients at zero longitude (parallel to the x-z plane), but when it straddles the meridian, at +/-10° longitude, and peak overrepresentation is more focused on lower latitude.

### 8 | Profiling Schellman loop/beta turn conformations at alpha-helical C-termini

The Schellman loop[12,13] (see CT labels in Table S4) is a six-residue motif defined by a nested pair of (6>1, 5>2) BB H-bonds (in the loop frame) that is commonly enabled by a +φ conformation at the fifth loop residue. The loop is frequently found at the helix C-terminus, where its defining H-bonds provide a dual C-cap[15,16]. The motif's inner nested 5→2 H-bond is ($i$ +3) → $i$, like the beta-turn H-bond, so helix-terminal Schellman loops in which the helix does not overlap with the central residues of the loop's 2→5 segment (thus ruling out a beta-turn classification for that segment) contain beta turns with 4→1 H-bonds that correspond to the loop's 5→2 H-bond. The ExploreTurns database contains examples of these structures (Figure S9d), as well as examples of structures in which beta turns overlap Schellman loops at different registrations in the loop frame, including at BB positions shifted C-terminally from the 2→5 segment (Figure S9a-c).

The constraints imposed by the Schellman loop's characteristic pair of H-bonds (along with other BB H-bonds that may occur in the motif, see Table S4) ensure that the orientation of the helix with respect to a beta turn at its terminus depends largely on two factors: the position within the turn of the loop's +φ residue, which determines the loop's registration in the turn frame, and the beta-turn geometry, which determines the orientations of the turn's BB H-bond donors. Analysis with ExploreTurns shows that of these factors, the position of the +φ residue is primary, since the distribution of turn BB geometries for each of the four possible +φ positions is characterized by a majority geometry which establishes a principal approximate helix/turn orientation. Figure S9 presents example structures and ExploreTurns selection criteria for these four conformations, classified by the position of the +φ residue in the beta turn.

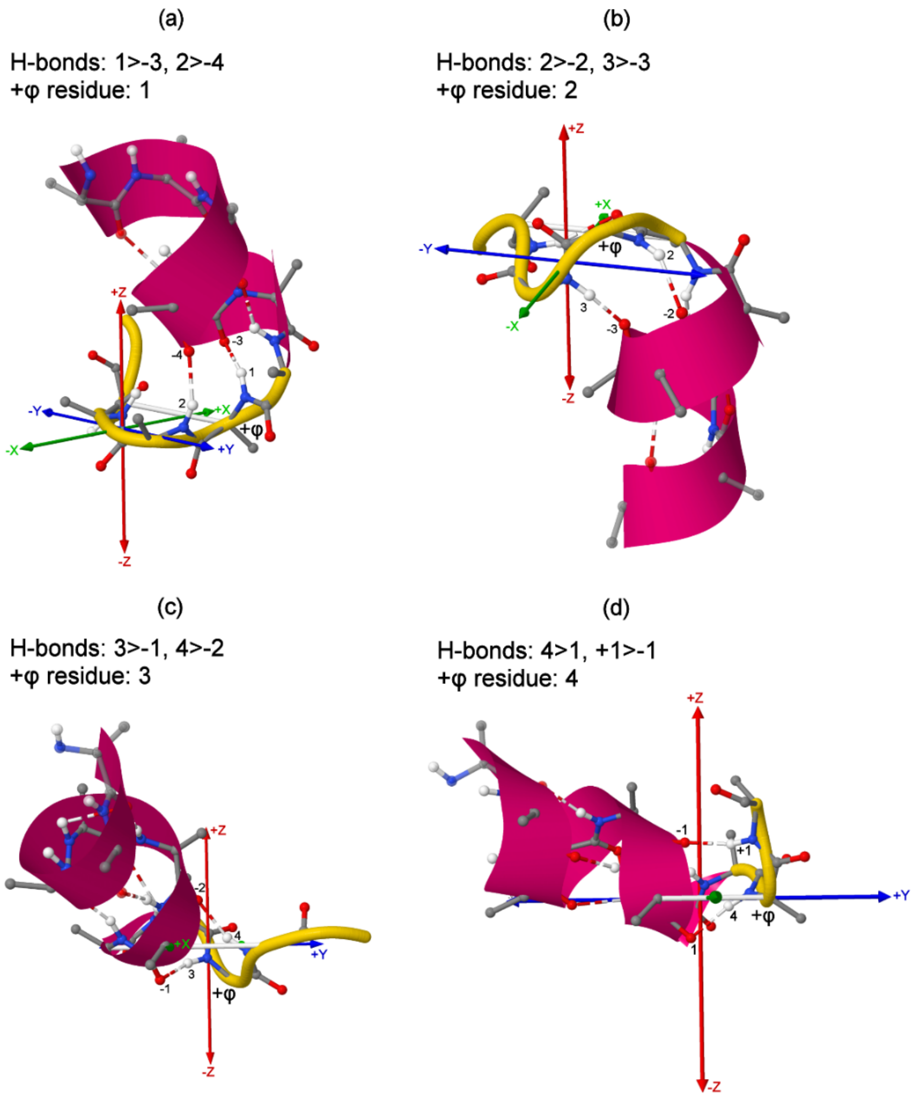


**Figure S9. Principal conformations of Schellman loop/beta turn combinations at alpha-helix C-termini.** The helix/turn orientation in a Schellman loop/beta turn combination depends on both the loop's registration within the turn frame, which can be measured by the position of the loop's +φ residue (which enables the loop's defining (6>1, 5>2) nested H-bond pair), and the turn geometry, but each +φ position is associated with a majority BB geometry that establishes a principal (approximate) helix/turn conformation. In (a)-(c), the +φ position advances C-terminally, bringing the helix into the turn, and in (d) the loop's inner nested 5>2 H-bond corresponds to the turn's 4>1 H-bond. Example structures for the four conformations are shown here; for clarity, the only helix-terminal H-bonds displayed are the loop's defining nested pair. ExploreTurns selection criteria are given for each conformation; before entering a new set of criteria, click ***Clear All Criteria*** to erase the previous set. **(a)** Turn residue 1 is +φ, and a type VIII turn BB geometry orients the helix towards (-x, +z) [***β-turn type/cluster***: *AB*, ***DSSP symbols***: **Hx<x***>****, ***BB H-bonds***: *1>-3, 2>-4*]. **(b)** Turn residue 2 is +φ, and a type I' turn geometry (which requires a +φ conformation for residue 3 as well) orients the helix towards -z [***β-turn type/cluster***: *I'*, ***DSSP symbols***: ***H<x***>****, ***BB H-bonds***: *2>-2, 3>-3*]. **(c)** Turn residue 3 is +φ, and a combination of right- and left-handed helical conformations at the turn's two central residues orients the helix towards (+x, +z) [***β-turn type/cluster***: *Dd* or *Ad*, ***DSSP symbols***: ****<H***>****, ***BB H-bonds***: *3>-1, 4>-2*]. **(d)** Turn residue 4 is +φ, and an almost exclusive type I turn geometry orients the helix towards -y (with a +z component). The loop's inner nested 5>2 H-bond corresponds to the turn's 4>1 H-bond [***DSSP symbols***: ****<H***>****, ***BB H-bonds***: *4>1, +1>-1*].

### 9 | Investigating the depth dependence of beta-turn geometry

The extent to which beta-turn BB geometry is influenced by the structures in a protein external to turns is an open question. In the classical type system[18], turns are partitioned by the BB dihedral angles of their central residues into eight structurally classified types {I, I', II, II', VIa1, VIa2, VIb, VIII} and a catch-all category (type IV) for outliers. Type IV turns frequently exhibit (φ, ψ) values which lie just outside the allowed ranges for the structurally classified types, and this observation has led to the description of many type IV turns as distortions of these types[19]. If turns are substantially distorted due to interactions with other structures in a protein, then the distribution of their geometries might be expected to change with increasing depth beneath the solvent-accessible surface, since the frequency, nature or impact of the interactions may change with depth. Since ExploreTurns includes depth as a selection criterion and also reports the type distributions of the selected structure sets, the tool can be used to profile changes in the turn type distribution with depth, and Figure S10 plots type fractions vs. depth.

The interpretation of Figure S10 is not straightforward, due to complications associated with depth measurement. ExploreTurns' depth measurements are taken from the BB atoms of the two central turn residues, since including SC atoms would introduce uncertainties related to the variability of SC structures and potential SC positional uncertainty. However, this procedure introduces a bias towards shallower depths for classical types I', II, and II', because these types each show a very strong preference for Gly at one of the central turn positions (due to requirements for +φ conformations at these positions), and Gly's lack of a SC reduces the average measured depth. This bias should be considered an artifact rather than a structurally meaningful finding, since it is due entirely to well-understood differences in SC content between types, and Figure S10 includes a measure of the fraction of Gly residues at the two central turn positions to serve as a type-independent proxy for the effect.

Biases due to preferences in turn types for SCs other than Gly also likely affect the results in Figure S10, but a comprehensive evaluation of these is beyond the scope of this work. The figure does, however, show a trend in the type distribution that is unlikely to result from SC bias: the fraction of structurally unclassified turns increases proportionally by 52% with depth, due to an effective "transfer" of turn fraction from classical type I to type IV (see the figure caption for details). When turns are classified by BB cluster medoids rather than classical types, this effect is also seen: the type I-associated BB cluster loses an overall turn fraction of 8.5%, while the outlier category gains an 8.5% fraction, quadrupling its share of all turns to 11.3%.

The apparent transfer of a substantial fraction of all turns, as depth increases, from type I to the structurally unclassified category may reflect geometric distortions due to increasing and/or more impactful contacts with adjacent components of the protein, increasing stress transmitted through the BB by neighboring structures or structures that incorporate beta turns, or increased binding of ligands or prosthetic groups by turns. The pause in type IV's rise between 6.5 Å and 9.0 Å may reflect a saturation of interactions between turns and external structures reached on average near 6.5 Å, the effects of cavities in proteins, or some combination of these or other effects. Note that the structural "bins" in the histogram become wider at greater depth, since the count of structures falls with depth; just 4% of turns occur at depths greater than 9.0 Å.

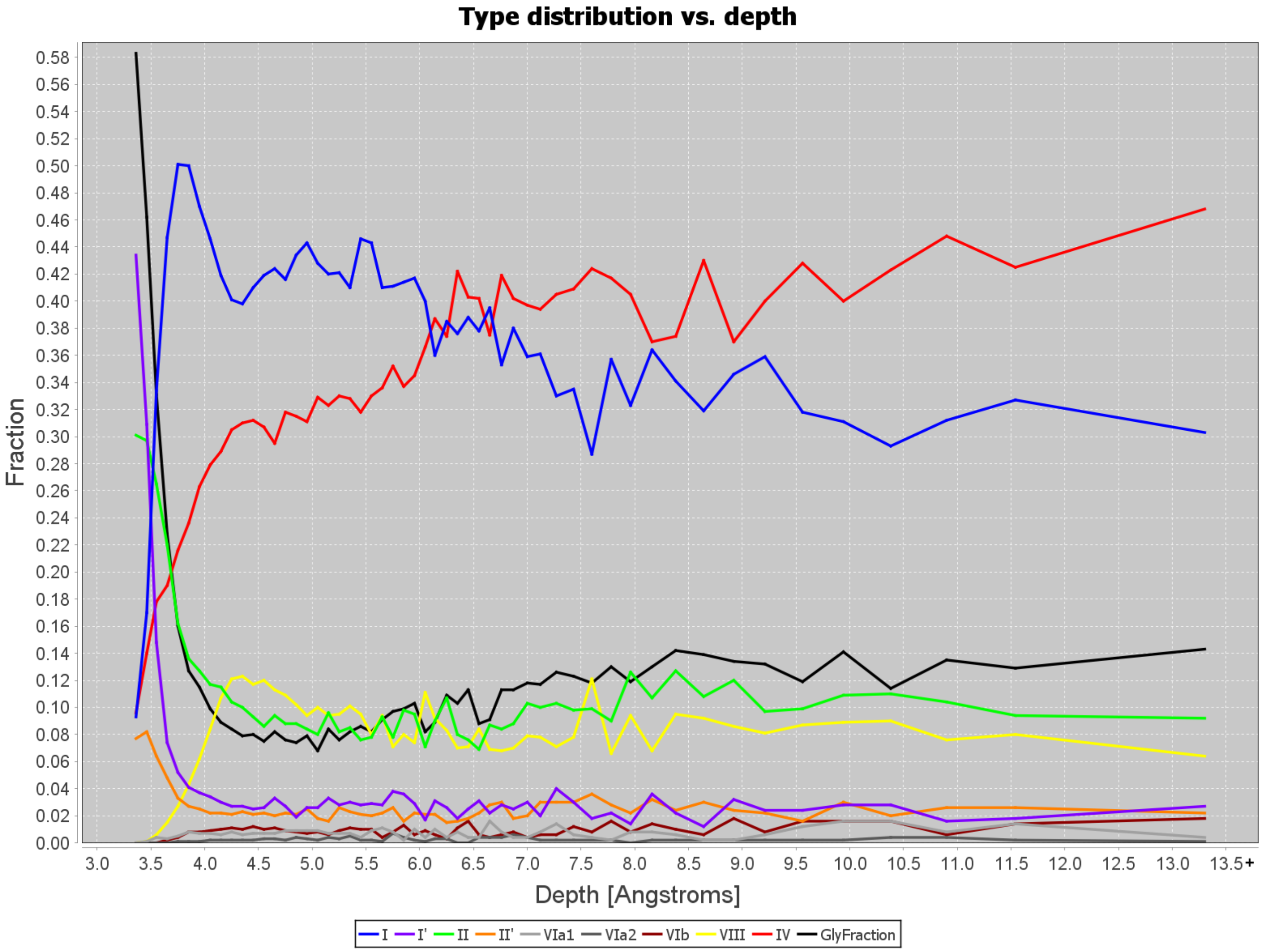


**Figure S10. Variation of the classical type distribution with depth beneath the solvent-accessible surface.** Abundance fractions of the classical types in the beta-turn dataset are plotted vs. depth beneath the solvent-accessible surface[20]. The fraction of Gly residues at the two central turn positions is also plotted (black), as a type-independent proxy for the bias toward shallower depths due to high Gly frequencies in types {I', II, II'}, which require a +φ conformation at a central residue. This bias, together with other possible SC-related biases, complicates the interpretation of the plot; the type fractions shown can reflect combinations of genuine structural preferences (such as the tendency for the chain-reversing turns of 2:2 beta hairpins, which are commonly of type I' or II', to occur near the surface) and SC bias. However, the graph shows a trend that is unlikely to be the product of SC bias: near a depth of 4.5 Å, beyond the point at which the Gly bias has subsided and the fraction of structurally unclassified (type IV, red) turns has "recovered" from its bias-related suppression, type IV begins to rise again, with increasing steepness, until it exceeds the abundance of type I (blue) near 6.0 Å. Type IV then fluctuates, with no net gain, until near 9.0 Å, where it begins to rise again. By maximum depth, type IV's share of all turns has increased from 31% to 47% (a 52% proportional increase), largely at the expense of type I. Note that the structural "bins" in this histogram become wider at greater depth, since the count of structures falls with depth.

### 10 | Computing sequence motif overrepresentation and p-value

Statistical tools[1,2,5,16,21,22] are used in ExploreTurns to evaluate the over/under-representation and the significance of sequence motifs which specify up to three AAs at three positions anywhere in the turn or tails (for motifs that specify more than three AAs, only the motif's count and its abundance fraction in the selected set is reported). The degree of over/under-representation of a motif is measured by its fractional overrepresentation $(O - E)/E$, where $O$ is the motif's observed count in a set of structures and $E$ is its expected count under a suitable null model. The statistical significance of a motif in a structure set is measured by its p-value, which represents the probability that the motif's departure from its expected count would be as large or larger than its actual value by chance alone under the null model. Lower p-values indicate a smaller chance that the null model is consistent with the data, and consequently a greater chance that the model is incorrect and the motif is significantly over- or underrepresented.

For a single-AA sequence motif, the null model specifies that the probability $P$ of the motif's occurrence in the set is equal to the position-independent probability of the occurrence of the motif's AA anywhere in proteins, which is computed as the overall abundance fraction of the AA in the complete dataset of protein chains used in the study: $P = C_{AA}/T$, where $C_{AA}$ is the count of the AA in the set of all chains and $T$ is the total number of residues in the chain set. The motif's expected count in the structure set is then $E = NP$, where $N$ is the size of the set, and its fractional overrepresentation is $(O - NP)/NP$. The p-value is computed as the combined area underneath the two tails of the binomial distribution with parameters $N$ and $P$ that represents the null model of random motif occurrence, where the tails represent the probability that the motif's observed count would equal or exceed its value by chance alone under the model.

A sequence motif which specifies two amino acids at two positions (pair motif) is evaluated by measuring the significance of the two-factor effect in a 2x2 contingency table[21] in which the rows of the table indicate the presence or absence of the AA specified at the motif's first position and the columns indicate the presence/absence of the AA at its second position. For example, the following contingency table might be used to evaluate the pair motif D1P2 that specifies Asp at turn position 1 and Pro at turn position 2 in a set of 100 turns:

| | $Pro2$ | $\overline{Pro2}$ |
|---|---|---|
| $Asp1$ | 10 | 15 |
| $\overline{Asp1}$ | 20 | 55 |

Here the labels indicate the presence or absence (barred label) of each AA at its position, and the table contains the observed counts $O_{ij}$ for each of the four possible combinations of presence/absence of the two AAs at positions 1 and 2; when D1P2 is present in a turn, it is counted in cell (1,1) of this table.

The null model for a pair motif specifies independent occurrence of each AA at each position, which indicates no pair synergy between the motif's individual components. Under this model, the probability that a turn falls into cell $(i, j)$ of the 2x2 observed table, where $i$ indexes the rows and $j$ the columns, is:

$$P_{ij} = P_{1(i)} \times P_{2(j)} = \frac{M_{1(i)}}{N} \times \frac{M_{2(j)}}{N}$$

where $P_{1(i)}$ represents the probabilities of the turn either containing the first AA specified by the motif at its particular turn position ($i = 1$) or not containing that AA at that position ($i = 2$), and $P_{2(j)}$ represents the independent probability of the presence or absence of the second AA at its position. $M_{1(i)}$ and $M_{2(j)}$ are the table's margin totals for its rows ($M_{1(i)}$), formed by summing the table over its columns, and its columns ($M_{2(j)}$), formed by summation over its rows, and $N$ represents the total number of structures in the set.

The expected counts for the cells of the 2x2 table under the null hypothesis of independence are:

$$E_{ij} = N \times P_{ij} = \frac{M_{1(i)} \times M_{2(j)}}{N}$$

so the expected count for the motif, in cell (1,1) of its table, is:

$$E_{11} = \frac{M_{1(1)} \times M_{2(1)}}{N}$$

and its fractional overrepresentation is:

$$\frac{(O_{11} - E_{11})}{E_{11}}$$

where $O_{11}$ is the motif's observed count.

To compute the motif's p-value, a chi-squared metric is computed which measures the distance between the observed counts in the four cells of the motif's 2x2 table and their corresponding expected counts:

$$\chi^2 = \sum_{i=1}^{2} \sum_{j=1}^{2} \frac{\left(O_{ij} - E_{ij}\right)^2}{E_{ij}}$$

The p-value is obtained by comparison of this metric with the chi-squared distribution with one degree of freedom, which represents the (single) two-factor effect, or pair synergy, present in the observed table. P-values are not corrected for multiple testing error so that they may be used as reference values for individual motifs of interest.

Triplet motifs are evaluated using a method analogous to that applied for pair motifs. For a triplet motif in a turn set, the null model specifies no three-factor effect in the 2x2x2 contingency table for the motif, in which each dimension represents the presence or absence of one of the three individual AA components in the motif. This model specifies that there is no triplet synergy associated with the motif; that is, that there is no association between any pair of AAs in a motif and the remaining, single AA.

The expected counts for the null model of no three-factor effect in a triplet motif's 2x2x2 observed table cannot be computed using a closed-form expression such as that used to calculate $E_{ij}$ in the 2x2 tables of pair motifs, but instead must be calculated via iterative proportional fitting (IPF)[21], which begins with a table of uniform estimated expected counts and applies, at each iteration, each of the three 2x2 tables of margin totals computed from the motif's 2x2x2 observed table to this table of estimates, modifying the estimates in each step until they converge to a desired level of precision. The resulting expected-count estimates are consistent with all three observed 2x2 margin tables, so they contain all three two-factor effects present in the data (along with the three single-factor effects associated with the individual AAs), and they represent the maximum likelihood estimates for the table's expected counts under the null model which lacks only the three-factor effect.

When a triplet motif is present in a turn, it is tabulated in cell (1, 1, 1) of its observed 2x2x2 table, so the motif's expected count is $E_{111}$, and its fractional overrepresentation is:

$$\frac{(O_{111} - E_{111})}{E_{111}}$$

where $O_{111}$is the motif's observed count.

To compute the motif's p-value, a chi-squared metric is computed to measure the distance between the observed counts in the eight cells of the motif's 2x2x2 observed table and their corresponding expected counts:

$$\chi^2 = \sum_{i=1}^{2}\sum_{j=1}^{2}\sum_{k=1}^{2}\frac{\left(O_{ijk} - E_{ijk}\right)^2}{E_{ijk}}$$

The motif's p-value is obtained by comparison of this metric with the chi-squared distribution with one degree of freedom, which represents the (single) three-factor effect, or triplet synergy present in the observed table. P-values are not corrected for multiple testing error so that they may be used as reference values for individual motifs of interest.

Note that the absence of synergy in a pair motif means only that each AA in the motif occurs independently of the other, and that the motif is therefore not likely to represent a significant interaction between the two AAs. This means that the effect of the simultaneous occurrence of the two AAs is likely to be no more important than the sum of their individual effects, but it does not mean that the occurrence of the pair has no importance, since the pair is an instance of both single-AA motifs, each of which may themselves be significant. In order to fully evaluate the importance of a pair motif, it is therefore necessary to consider the importance of its components; for example, the D1R3 motif is overrepresented in type I turns by 45%, likely reflecting its associated intra-turn salt bridges, but its D1 component is overrepresented in the type by 159%, reflecting the SC/BB H-bonding that makes D1 the most important of all H-bond motifs in the type. The analogous consideration applies to triplet motifs: the absence of triplet synergy means only that the motif is not likely to represent a significant interaction between its pair components and the single-AA components which complement them; it does not necessarily mean that the occurrence of the triplet has no importance, since it is also an instance of its underlying pair and single-AA motifs.

### 11 | Definitions of the α/β conformations

Table S8 defines the "right-handed" and "left-handed" (αR, αL, βR, βL) BB conformations, which provide a simplification of Ramachandran space useful for selecting conformations in compound-turn labels or in the ***α/β or (φ,ψ) BB Conf. (A|a|B|b|F|f|Y|y)N*** box. The romanized labels used to select each conformation are also given.

**Table S8. Dihedral-angle ranges for the α/β conformations**

| **Conformation** | $\varphi_{min}$ | $\varphi_{max}$ | $\psi_{min}$ | $\psi_{max}$ |
|---|---|---|---|---|
| **αR/A** | -150° | -20° | -90° | 40° |
| **αL/a** | 20° | 150° | -40° | 90° |
| **βR/B** | -180° | -20° | 80° | 180° |
| **βL/b** | 20° | 180° | -180° | -80° |
